# Natural hybridization reveals incompatible alleles that cause melanoma in swordtail fish

**DOI:** 10.1101/2019.12.12.874586

**Authors:** Daniel L. Powell, Mateo Garcia, Mackenzie Keegan, Patrick Reilly, Kang Du, Alejandra P. Díaz-Loyo, Shreya Banerjee, Danielle Blakkan, David Reich, Peter Andolfatto, Gil Rosenthal, Manfred Schartl, Molly Schumer

## Abstract

The establishment of reproductive barriers between populations is the key process that fuels the evolution of new species. A genetic framework for this process was proposed over 80 years ago, which posits “incompatible” interactions between genes that result in reduced survival or reproduction in hybrids. Despite this foundational work, progress has been slow in identifying individual genes that underlie hybrid incompatibilities, with only a handful known to date. Here, we use a combination of approaches to precisely map the genes that drive the development of a melanoma incompatibility in swordtail fish hybrids. We find that one of the genes involved in this incompatibility also causes melanoma in hybrids between distantly related species. Moreover, we show that this melanoma reduces survival in the wild, likely due to progressive degradation of the fin. Together, this work represents only the second case where the genes underlying a vertebrate hybrid incompatibility have been identified and provides the first glimpse into the action of these genes in natural hybrid populations.

**One sentence summary:** Using a combination of mapping approaches, we identify interacting genes that lead to melanoma in hybrids and characterize their effects in natural hybrid populations.

## Main text

The emergence of reproductive barriers between populations is the first step in the process of speciation and is a driving force behind the diversity of life on earth, yet we still know surprisingly little about how it occurs at the genetic level. One of the foundational theories in evolutionary biology, the Dobzhansky-Muller model of hybrid incompatibility (*1–3*), posits that new mutations which arise in diverging species can interact negatively in hybrids, generating lower hybrid viability or causing hybrid sterility. Despite a large body of empirical work that provides support for the general predictions of this model (reviewed in *4*), progress in this area has been limited by a lack of knowledge about which genes interact to generate hybrid incompatibilities. Although substantial effort has been devoted to this problem, only a dozen incompatible interactions have been mapped to the single-gene level (*5*). With so few known cases it has been difficult to identify common genetic and evolutionary mechanisms underlying the emergence of incompatibilities, though several have been proposed (*6–8*). This represents a major gap in our understanding of the origin of reproductive barriers between species.

Understanding mechanisms of isolation has become more pressing in recent years as it has become evident that hybridization between species is common. Over the last decade, evidence of previously unknown hybridization events have been discovered in the genomes of species from across the tree of life, including our own (e.g. *9–13*). This has led to an increasing consensus that hybridization is a ubiquitous force in the evolutionary histories of diverse groups of species. With this realization has come a renewed interest in identifying hybrid incompatibilities and understanding how these genes act as barriers in nature.

Of hybrid incompatibilities that have been mapped to the single-gene level, almost all of them have been identified using crosses between deeply diverged species (*5*), in part because such species have been more tractable to study in a lab environment (*4*). However, the majority of these species no longer hybridize in nature. As a result, it is unclear whether these mapped incompatibilities were important in the initial divergence between species or arose long after they ceased to exchange genes.

One such example is the *xmrk* gene in swordtail fish (genus *Xiphophorus). Xmrk* is one of only two genes in vertebrates known to drive hybrid incompatibility (the other being the regulator of mammalian recombination hotspots, *prdm9*; *14*), and was one of the earliest described hybrid incompatibilities (*15*). In crosses between *X. maculatus* and *X. hellerii*, a malignant melanoma develops in a subset of F_2_ hybrids, emanating from natural pigmentation spots on the body and fins. Decades of research have revealed that this hybrid incompatibility is the result of an interaction between the *xmrk* gene derived from *X. maculatus* and an unknown interacting locus derived from *X. hellerii* (*16*).

Despite past work demonstrating the role of *xmrk* in the development of hybrid melanomas, its importance as a barrier between species has been debated (*17*). The reasons for this are two-fold. First, this work has focused on deeply diverged species (~3 million years diverged) that do not naturally hybridize (*18*), making it unclear whether the *xmrk* incompatibility plays a role in reproductive isolation in natural populations. Second, given that melanoma in *X. maculatus* x *X. hellerii* laboratory hybrids develops later in life, it is unclear if it has a major effect on survival and reproduction (*17*).

We identified a phenotypically similar melanoma in natural hybrids formed between the swordtail fish species *X. birchmanni* and *X. malinche*. Because *X. birchmanni* and *X. malinche* are closely related and hybridize in the wild (~0.5% differences per base pair and ~250,000 generations diverged; *19*), this system provides an exciting opportunity to characterize the effects of melanoma in natural populations during the early stages of speciation. Notably, in some populations, hybrids develop melanoma early in life (13 ± 4% of males develop melanoma before sexual maturity; Fig. S1). While a subset of natural hybrids between *X. birchmanni* and *X. malinche* are viable and fertile, there is evidence for many segregating incompatibilities (*20, 21*) and genome-wide signatures of selection against hybrids (*22*). Like in other species, few incompatibilities have been directly mapped and the individual genes underlying hybrid incompatibilities are largely unknown.

Melanoma in *X. birchmanni* x *X. malinche* hybrids develops from a phenotype derived from *X. birchmanni* called the “spotted caudal,” which is a dark blotch on the caudal fin generated by clusters of macromelanophore cells (Fig. 1, Fig. S2; *23*). The spotted caudal trait occurs at intermediate frequencies in *X. birchmanni* but is absent from *X. malinche* populations (Fig. 1, Fig. S3, Fig. S4). Some hybrid populations have a high frequency of the trait (Fig. 1) and exhibit phenotypes that extend far beyond the range of those observed in *X. birchmanni* (Fig. 1). Long-term tracking of hybrids in the lab documents the progression of the trait from a phenotype typical of the *X. birchmanni* spot to the unusually expanded trait found in some hybrids (Fig. 1, Fig. S1; Supporting Information 1.1).

**Figure 1.**
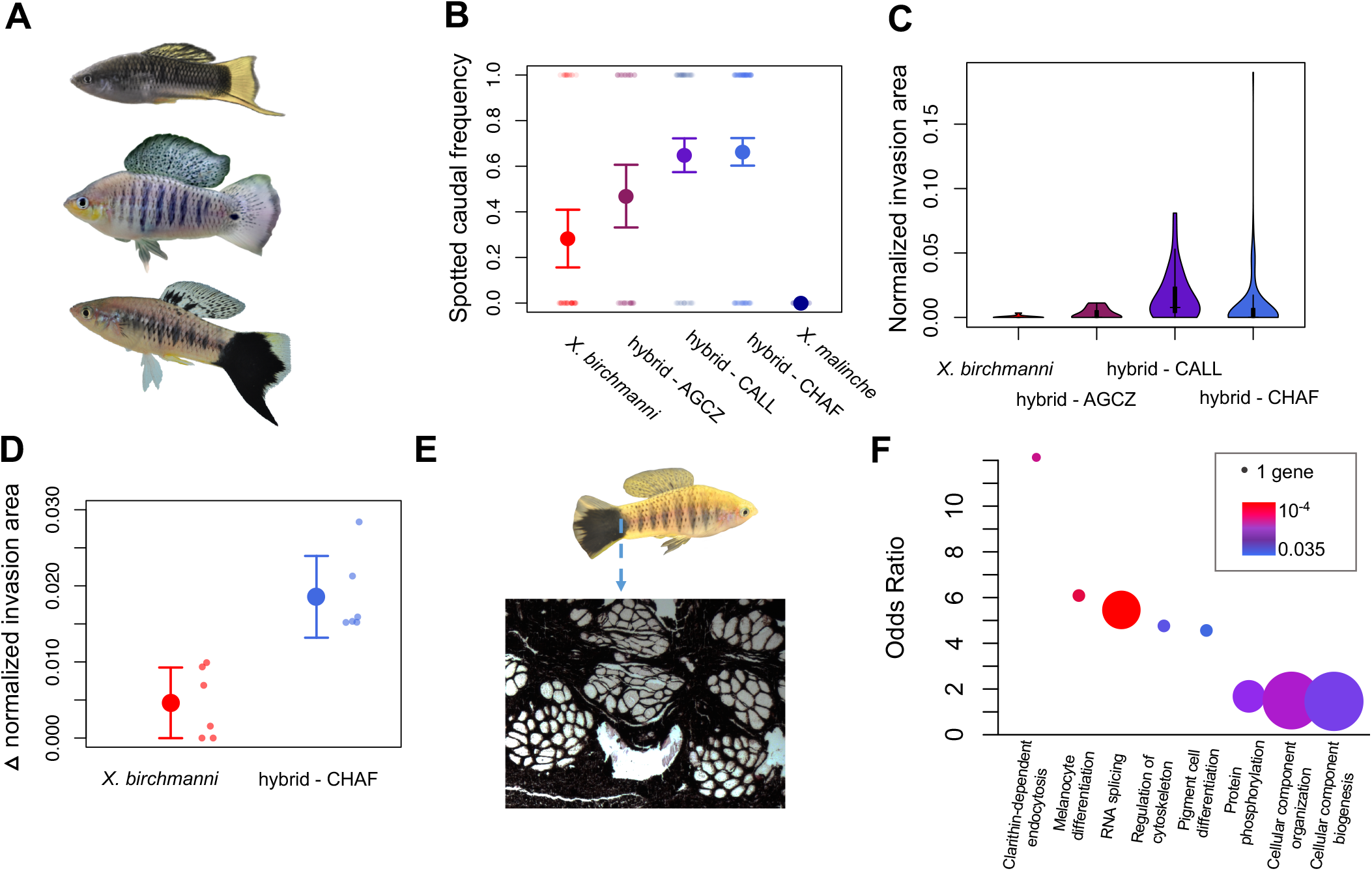
Hybridization generates a high incidence of melanoma. **A**) Naturally hybridizing species *X. malinche* (top) and *X. birchmanni* (middle) differ in a number of morphological traits including the presence of a melanin pigment spot that is polymorphic in *X. birchmanni*. A hybrid in which this spotting phenotype has transformed into a melanoma can be seen at the bottom of panel A. **B**) While *X. birchmanni* populations segregate for the presence of this spot, the trait is absent in *X. malinche* populations. Notably, hybrid populations tend to have high frequencies of this trait. **C**) Not only is the trait at higher frequencies in hybrid populations, it is also covers more of the body. Shown here is invasion area, or the melanized body surface area outside of the caudal fin (normalized for body size). **D**) Monitoring of hybrid and parental individuals in the lab over six-months indicates that the spot expands more over time in hybrids. **E**) Histological results showing a cross section of the caudal peduncle from a Chahuaco falls hybrid with melanoma (at 10X magnification). Melanoma cells invading the body and muscle bundles are visually evident. **F**) Gene ontology categories enriched in melanoma tissue compared to normal caudal tissue (Supporting Information 1.2; thinned with REVIGO; *43*). The size of the dots reflects the number of genes identified and the color corresponds to the p-value. Categories with undefined odds ratios are not plotted here but are listed in Table S1. In **B** and **D**, solid dots show the mean and whiskers indicate two standard errors of the mean. Transparent dots show the raw data for each individual.

Histological sections from hybrid individuals revealed penetration of melanocytes into the musculature and invasion of surrounding tissues, indicative of a malignant melanoma (Fig. 1, Fig. S5). We also performed mRNA-sequencing of hybrid individuals who varied in the degree of expansion of their spot (Fig. S2, S6). Functional enrichment analysis indicated changes in the regulation of a number of melanoma-associated gene categories, such as pigment cell differentiation and regulation of cytoskeletal organization, including several implicated in melanoma in other fish species (Fig. 1, Table S1, Supporting Information 1.2; *24*).

In contrast to what is observed in hybrids, melanoma emanating from the spotted caudal is extremely rare in non-hybrid individuals (Fig. 1; *6*). In hundreds of individuals collected from parental populations, we have not identified a single wild-caught *X. birchmanni* male with melanoma (1296 males collected from 2017-2019, 0 with melanoma). Lab-reared individuals of both groups indicate that environmental differences (e.g. levels of ultraviolet irradiance) do not underlie differences in the frequency of melanoma between hybrid and parental populations (Supporting Information 1.1). The presence of melanoma in hybrids but not in the parental species suggests that this melanoma is likely to be a hybrid incompatibility generated by interactions between alleles in the *X. birchmanni* and *X. malinche* genomes.

### Precise mapping of the melanoma incompatibility

We sought to map the genetic basis of the spotted caudal in *X. birchmanni* and the genetic basis of melanoma in interspecific hybrids. To facilitate mapping, we first generated and annotated chromosome-level *de novo* assemblies for both species using a 10X-based linked read approach followed by assembly into chromosomes with Hi-C data. The resulting assemblies contained 24 large scaffolds corresponding to the known chromosomes in *Xiphophorus (26;* Supporting Information 1.3). We annotated these assemblies using RNAseq data from eight organ types derived from F1 hybrids and with data from F1 embryos (Supporting Information 1.3). BUSCO analysis of the resulting annotated assemblies indicated high assembly completeness (~97% for both assemblies, Supporting Information 1.3; *27*).

Leveraging these new assemblies, we collected low coverage whole genome sequence data for 392 adult male *X. birchmanni* individuals from a single collection site and performed a genome-wide association study (GWAS) scanning for allele frequency differences between spotted cases and unspotted controls. We evaluated possible technical issues associated with population structure and low-coverage data (Supporting Information 1.4). Our analyses identified a strong association between the spotting pattern and allele frequency differences on chromosome 21 at an estimated false positive rate of 5% (Fig. 2, Fig. S7, S8; Supporting Information 1.4). Upon closer examination, two distinct signals are evident on this chromosome, approximately 5 Mb apart (Supporting Information 1.4, 1.5). The first peak is centered on the *xmrk* gene (Fig. S9; *28*), which arose through duplication of a gene homologous to the mammalian epidermal growth factor receptor approximately 3 million years ago (*11, 29, 30*). This finding is striking because *xmrk* has been previously shown to control pigmentation patterns as well as drive hybrid melanomas in distant relatives of *X. birchmanni* and *X. malinche* (*16*). The signal at the second peak on chromosome 21 contains a single gene, the melanosome transporter gene *myrip* (Fig. 2; see Supporting Information 1.4; *31*).

**Figure 2.**
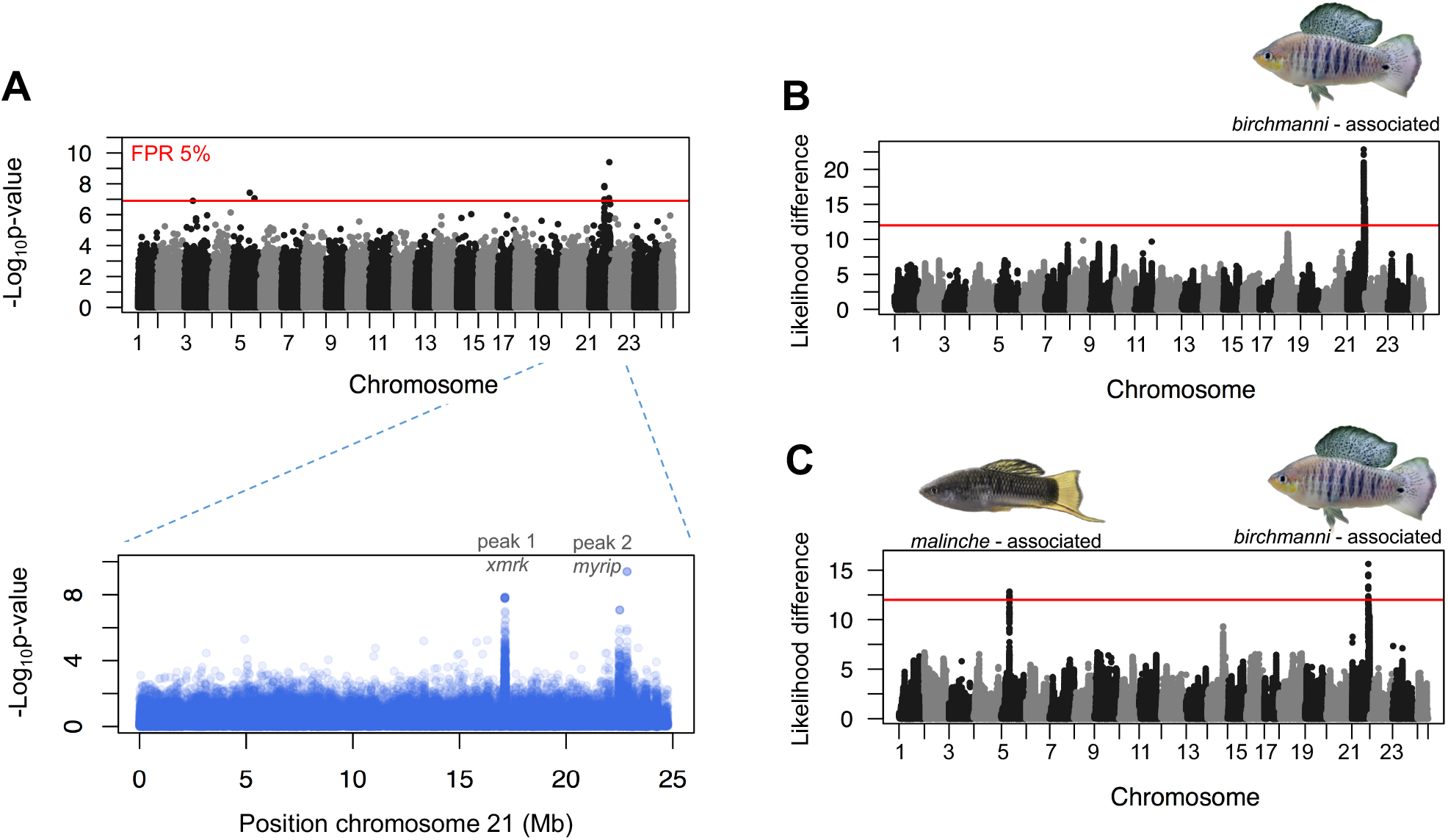
Combined genome-wide association and admixture mapping approaches identify the genetic basis of the melanoma hybrid incompatibility. **A**) Results of genome-wide association scan for allele frequency differences between spotted cases and unspotted controls. In the upper panel, results can be seen for all chromosomes and the red line shows the genome-wide significance threshold, determined by permutation (Supporting Information 1.4). In the bottom panel the results from chromosome 21 are plotted. Chromosome 21 has the strongest association with the presence of the spotting pattern and there are two regions on this chromosome associated with the spot. **B**) Admixture mapping in hybrids identifies a strong association between *X. birchmanni* ancestry on chromosome 21 and spot presence, as predicted by GWAS results. Plotted here are log likelihood differences between models with and without ancestry at the focal site included as a covariate. The red line shows the genome-wide significance threshold, determined by permutation (Supporting Information 1.6). **C**) When we treated melanin invasion as the focal trait and map associations with ancestry, we again identified associations with *X. birchmanni* ancestry on chromosome 21 but also identified a second region on chromosome 5 associated with *X. malinche* ancestry.

To characterize the genetic basis of the melanoma phenotype in hybrids, we used an admixture mapping approach. We focused our mapping efforts on a hybrid population with high incidence of melanoma (19 ± 3% of adult males, the Chahuaco falls population, Fig. S3). To infer local ancestry patterns, we generated ~1X low-coverage whole genome sequence data for 209 adult males from this population and inferred local ancestry by applying a hidden Markov model to 680,291 ancestry informative sites genome-wide (*20, 22, 32*; ~1 ancestry informative site per kb, Supporting Information 1.6). Simulations and analyses of known crosses indicate that we expect to have high accuracy in local ancestry inference (Fig. S10-S13, Supporting Information 1.6).

Using these data, we performed admixture mapping for two traits: spot presence and melanocyte invasion of the body (59% of individuals had spots and 19% of individuals had melanoma). Admixture mapping for the presence of the spotted caudal revealed one highly significant region on chromosome 21 associated with *X. birchmanni* ancestry (Fig. 2; Supporting Information 1.6). In part because of the lower resolution of admixture mapping this peak is broad but the signal occurs in the same regions identified by our GWAS scan (i.e. near *xmrk* and *myrip;* see Supporting Information 1.6), confirming that the genetic basis of the spot is the same in *X. birchmanni* and in hybrids. Simulations accounting for effect size inflations due to the winner’s curse (*33*) suggest that *X. birchmanni* ancestry in this region explains approximately 75% of the variation in spot presence or absence (Supporting Information 1.6).

Admixture mapping for melanoma (melanocyte invasion) identified an additional significant region, this time on chromosome 5. Instead of being associated with *X. birchmanni* ancestry, melanocyte invasion was associated with *X. malinche* ancestry (Fig. 2). The presence of an *X. birchmanni* associated region for the spotted caudal and an *X. malinche* associated region for melanocyte invasion suggests that the melanoma phenotype could be a classic hybrid incompatibility, where hybrid breakdown occurs as a result of combining these two regions of distinct ancestry. To evaluate this hypothesis, we used a contingency test to ask whether the presence of melanoma is dependent on the combination of *X. birchmanni* ancestry at the chromosome 21 peak and *X. malinche* ancestry at the chromosome 5 peak. We found a non-random association between ancestry at the chromosome 21 and chromosome 5 regions and the prevalence of melanoma (Fisher’s exact test, p=0.0005, Fig. 3). Individuals heterozygous for *X. malinche* ancestry at this region on chromosome 5 appear to have a lower risk of melanoma (Fig. 3). Moreover, regardless of melanoma phenotype, spotted individuals that were heterozygous in this region had smaller spots than individuals that were homozygous (t-test on log-normalized area p=0.007; Fig. S14).

**Figure 3.**
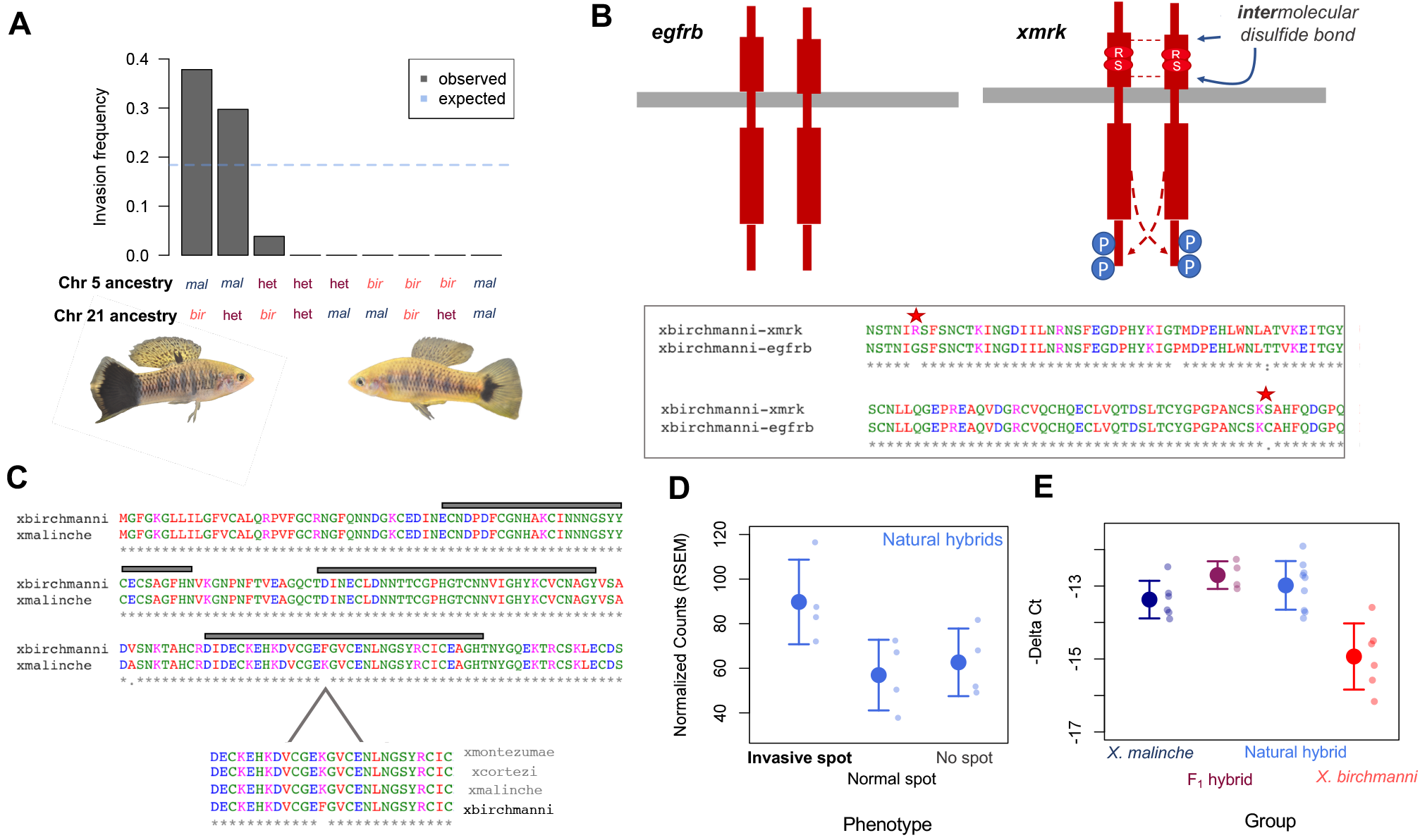
Interactions between chromosome 5 and 21 are associated with melanoma in hybrids. **A**) Individuals with *X. malinche* ancestry in the associated region on chromosome 5 and *X. birchmanni* ancestry on chromosome 21 are more likely to develop melanoma (Fisher’s exact test: p=0.0005). The blue line shows the expected proportion of cases if melanoma risk were equally distributed among spotted individuals (i.e. among individuals with at least one *birchmanni* allele at chromosome 21). Note that we only had one observation for the *bir-bir* and *bir-het* genotypes. **B**) The *xmrk* sequence in *X. birchmanni* harbors two mutations (G364R and C582S) that transform *xmrk* to a constitutively active state (*34, 44*). The schematic compares the ancestral form of the protein (*egfrb*) to the predicted structure of *xmrk* in *X. birchmanni*. Proteins are shown in red and the cell membrane in gray. In *xmrk*, residues R364 and S582 promote intramolecular disulfide bonds that cause protein dimerization and phosphorylation (blue circles; *34, 44*). The inset shows a partial clustal alignment of *X. birchmanni egfrb* and *xmrk* with these substitutions highlighted. Colors indicate properties of the amino acid, stars indicate locations where the amino acid sequences are identical. **C**) Clustal alignment showing the N terminus of *cd97* in *X. birchmanni* and *X. malinche*. In addition to other non-synonymous substitutions, we observe a change in a conserved epidermal growth factor binding domain (gray rectangles). The inset shows that the substitution found in *X. birchmanni* is not present in closely related species. **D**) *cd97* is upregulated in melanoma tissue from Chahuaco falls hybrids compared to nonmelanoma tissue. **E**) Real-time qPCR of caudal fin tissue from *X. malinche, X. birchmanni*, and natural and F1 hybrids indicates that hybrids have*X. malinche-like* expression of *cd97*. Expression was significantly different between *X. birchmanni* and all other groups (Tukey post-hoc test, corrected p<0.005) but not between *X. malinche* and hybrids (natural hybrids p=0.4; F1s p=0.2). In **D** and **E**, solid dots show the mean and whiskers indicate two standard errors of the mean. Transparent dots show the raw data for each individual.

Our GWAS identified associations between the spotted caudal and both *xmrk* and *myrip* (Fig. 2), making it initially unclear whether either or both these genes interacts with *malinche* ancestry on chromosome 5 to produce melanoma. Although both are likely important in the sense that they are associated with the spotting pattern that precedes melanoma, several lines of evidence suggest that we should focus on *xmrk* as the driver of melanoma in hybrids. First, *myrip* is not expressed in the caudal tissue of adults where melanoma develops, nor is it expressed in the melanoma itself (Fig. S15). In contrast, *xmrk* is expressed in caudal tissue and has higher expression in spotted than unspotted tissues (Fig. S15). In addition, functional studies have strongly linked *xmrk* to the development of melanoma. We identified two amino acid substitutions in *X. birchmanni* that fall within the extracellular domain of *xmrk* and have been shown to drive the oncogenic properties of *xmrk* in vitro (Fig. 3; *34*), and transgenic studies have demonstrated that overexpression of *xmrk* causes the formation of tumors (*24*, *35*, *36*). While *myrip* may not be directly involved in the development of melanoma, past work in other swordtail species has suggested the presence of “patterning” loci linked to *xmrk* (reviewed in *23*). Given *myrip”* s role in melanosome transport, we speculate that it could play a role in pigmentation patterning, which occurs in the first several weeks of life.

The region on chromosome 5 associated with *X. malinche* ancestry and melanoma contains only two genes (Fig. 2), a gene called *cd97* and a fatty acid transporter gene. Although this is unusually high resolution given our admixture-mapping approach, subsampling the data indicated that this scale of resolution is not due to noise in the data and instead is likely the result of high recombination rates in this region (Supporting Information 1.6). We therefore sought to better characterize the two genes in this region and investigate which was most likely to be involved in the development of melanoma in hybrids. The ortholog of *cd97* in mammals has known roles in epithelial metastasis and is associated with tumor invasiveness in several cancers (*37–39*). Accordingly, we find that *cd97* is upregulated in melanotic tissue in hybrids while the fatty acid transporter gene is not (Fig. 3); nor is this pattern of upregulation observed in any other gene within 100 kb of this region (Supporting Information 1.7). In addition, of five amino acid changes observed between *X. birchmanni* and *X. malinche* in *cd97*, one occurs in a conserved epidermal growth factor-like calcium binding site (Fig. 3).

Motivated by the observed expression differences across melanoma phenotypes in hybrids, we investigated differences in expression of *cd97* across genotypes. We found that *cd97* was expressed at low levels in the caudal fin tissue of *X. birchmanni*, regardless of spotting phenotype, but at similarly high levels in *X. malinche* and in natural and artificial hybrids (Fig. 3; ANOVA p-value =1^-5^, Tukey post-hoc: all groups different from *X. birchmanni* at p<0.005). Interestingly, higher expression of *cd97* in *X. malinche* and hybrids is not tissue-specific and does not appear to be driven by *cis-*regulatory differences (Fig. S16; Supporting Information 1.7). We do not know whether the link between *X. malinche* ancestry at *cd97* and melanoma is driven by coding or regulatory differences (see Supporting Information 1.7). However, in mammals, overexpression of *cd97* has been linked to tumor metastasis (*37–39*), hinting that it could play a similar role in this hybrid incompatibility.

Our results suggest that interactions between *xmrk* and *cd97* are linked to melanoma in hybrids between *X. birchmanni* and *X. malinche*. Because *xmrk* is involved in a melanoma hybrid incompatibility in two evolutionarily distinct cases (here and in *X. maculatus* x *X. helleriĩ*), it is natural to ask whether the interacting region is the same. Although the role of *xmrk* in the *maculatus-hellerii* hybrid incompatibility has been known for thirty years (*40*), the identity of the interacting gene has remained elusive. Decades of lab crosses have narrowed the interacting locus to a ~5 Mb region on chromosome 5 but have not yet identified the underlying gene, although candidates have been proposed (*16*). We too map the gene interacting with *xmrk* to chromosome 5, but the region we identify is distinct, found more than 7 Mb away from the region identified in the *hellerii*-*maculatus* cross. Alignments of chromosome 5 confirm that *cd97* is found in the same location in all four species (Fig. S17) and linkage disequilibrium between *cd97* and the region identified in *hellerii*-*maculatus* decays to background levels in hybrids (Supporting Information 1.6). Moreover, simulations show that we can rule out a lack of power to detect an association between this region and melanoma, assuming a similar effect size to that seen in the *hellerii-maculatus* cross (Fig. S18). Together, these results indicate that the melanoma hybrid incompatibility has a distinct genetic basis in the two cases, with different genes interacting with *xmrk*.

### Repeated evolution of a melanoma incompatibility

These mapping results are surprising because they suggest that a melanoma incompatibility involving *xmrk* emerged independently in two distinct lineages. Despite the evolutionary distance between these species (Fig. S19), it is possible that the melanoma incompatibility arose through similar evolutionary paths in both cases. Past work has indicated that *X. hellerii* and its relatives lack *xmrk* (*40*), either because the lineage leading to this clade diverged before *xmrk* arose (Fig. S19) or because of an ancient loss of *xmrk*. In contrast, many species in the lineage leading to *X. birchmanni* and *X. malinche* retained *xmrk* (Fig. S19; Supporting Information 1.8). Using analysis of local coverage around *xmrk* in *X. malinche*, *X. birchmanni*, and their relatives, we find that *xmrk* has been deleted in *X. malinche* since its divergence from *X. birchmanni* (Fig. S20; Supporting Information 1.8). We speculate that the loss of *xmrk* in *X. malinche* could have changed the level of constraint on interacting genes in this lineage, and if so, that similar mechanisms could be at play in *X. hellerii*.

### Selection on melanoma in natural populations

Although the melanoma that forms in *birchmanni* x *malinche* hybrids appears to be deleterious based on its development early in life (Fig. S1) and its malignancy (Fig. 1, Fig. S5), we wanted to evaluate its impact in natural hybrid populations. Sampling over several years, we found shifts in the frequency of the spotted caudal trait between juvenile and adult males (Supporting Information 1.1, 1.9). Specifically, in hybrid populations with high incidence of melanoma, juvenile males had a significantly higher frequency of the spotted caudal trait than adult males (Fig. 4). In contrast, this pattern was not observed in the *X. birchmanni* parental population or in a hybrid population with a low incidence of melanoma (Fig. 4). Phenotype tracking of lab-raised individuals shows that once it appears, the spotted area always expands over time, indicating that we do not expect spotted individuals to disappear due to phenotypic plasticity (Fig. 1; Supporting Information 1.1). We also did not find evidence for systematic shifts in ancestry between the juvenile and adult male life stages that could explain this pattern (Supporting Information 1.9). However, we do see an unusually strong shift towards *X. birchmanni* ancestry at the melanoma risk locus (in the top 1% of changes genome-wide, Fig. S21; Supporting Information 1.9).

**Figure 4.**
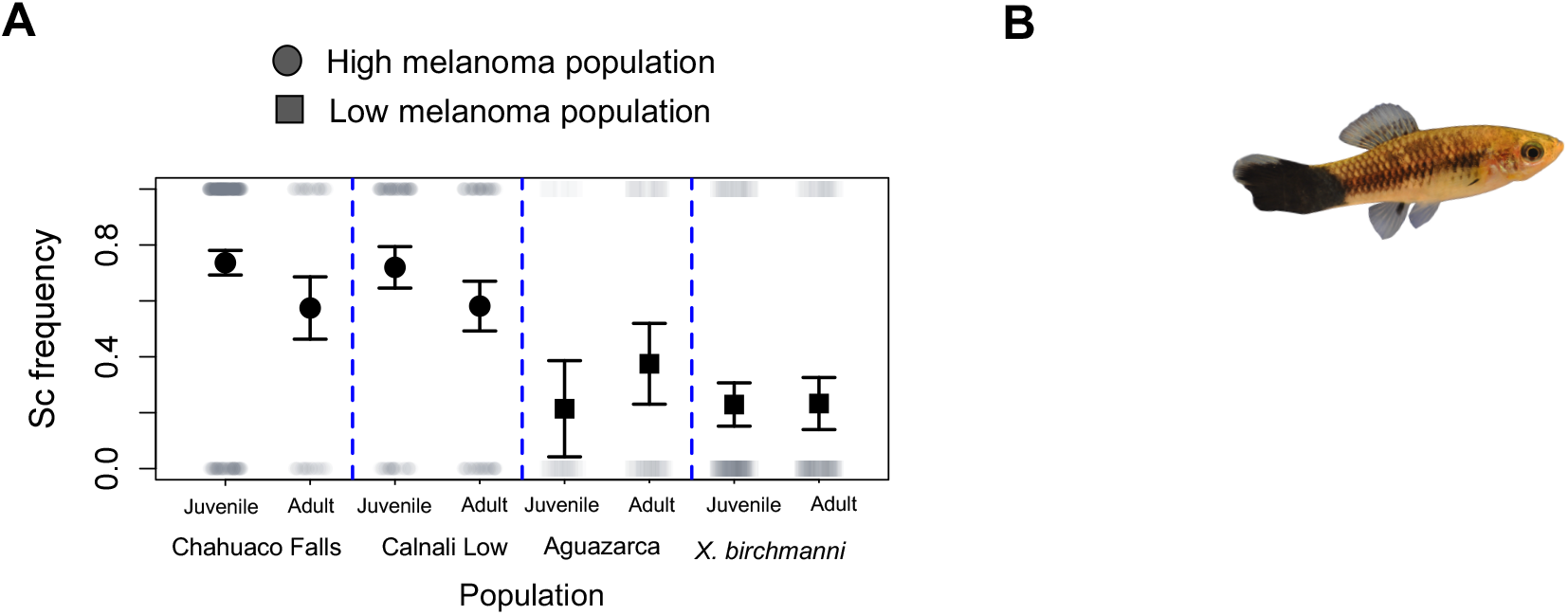
Impact of the spotted caudal melanoma in natural hybrid populations. **A**) We observe a lower frequency of adult males with the spotted caudal phenotype than juvenile males in two populations where melanoma is common (two proportions Z-test: Chahuaco falls - p=0.006, Calnali low - p=0.02). In contrast, this pattern is not observed in a hybrid population where melanoma is rare (Aguazarca - p=0.24) or in the *X. birchmanni* parental population (Coacuilco - p=1). **B**) A possible source of selection on hybrids with melanoma in natural populations is its impact on swimming ability or visual detectability to predators. Because of where the melanoma develops, it can cause the degradation of a fin essential in swimming (top) or the growth of tumors on the fin (bottom, overhead and side view of the same individual). Exaggerated spots can also make fish easier to see underwater (Supplementary Video 2). **C**) Visualization of the difference in fast-start response between individuals with low and high melanoma invasion (upper and lower 25% quantiles shown here). Note that this representation is for visualization only; statistical analysis is based on the linear model described in the text. **D**) ABC simulations indicate that the change in frequency of the spotting phenotype between juvenile and adult males is consistent with strong viability selection (Supporting Information 1.9). Shown here are posterior distributions of viability selection coefficients consistent with the observed frequency change data in Chahuaco falls (left) and Calnali low (right). Simulation details can be found in Supporting Information 1.9.

The paired shifts in spotting phenotype and ancestry at the melanoma risk locus, in the absence of substantial shifts in genome-wide ancestry, point to viability selection acting against hybrids with the spotted caudal during maturation (Supporting Information 1.9). We used an approximate Bayesian approach to infer the strength of viability selection consistent with the phenotypic shifts observed in hybrid populations. These simulations are consistent with extremely strong viability selection against spotted individuals in the two hybrid populations where melanoma is common (maximum a posterior estimate of *s* ~ 0.2; 95% confidence intervals 0.05-0.44 and 0.04-0.38; Fig. 4).

Because of where the hybrid melanoma develops on the fish, we hypothesized that it may impact swimming ability. Specifically, histology results showed substantial degradation of the muscle tissue that connects to the caudal fin in advanced melanomas (Fig. 1, Fig. 4, Fig. S5; Supporting Video 1). Motivated by this observation, we measured the impact of invasive spotted caudal on swimming performance using two approaches. We did not find differences between phenotypes in ability to swim against a current (Supporting Information 1.10). However, we did find that individuals with 3D melanoma had slower escape responses when startled (t=-2.6, p=0.014; Supporting Information 1.10). This result is intriguing since fish with expanded spotting are likely more visually obvious (Supporting Video 2), which could impact rates of detection by both avian and piscine predators.

Although the melanoma hybrid incompatibility appears to reduce survival, there may also be countervailing forces maintaining it in the population. Given the evidence for reduced survival of spotted individuals in populations with high rates of melanoma, it is surprising that this trait is still segregating in some hybrid populations (Fig. 4; Fig. S22, Supporting Information 1.9). Simulations suggest that high levels of migration from *X. birchmanni* paired with low effective population sizes would be required to maintain it at observed frequencies (Fig. S22, Supporting Information 1.9). However, because our inferences are based on only one component of fitness (viability) rather than a direct measure of fitness, it is important to stress that there may be much weaker effects of melanoma on overall fitness. Alternatively, other factors may explain its maintenance, such as mating advantages in individuals with large spots (*41*).

### Implications

The involvement of *xmrk* in a melanoma hybrid incompatibility in two distantly related swordtail species pairs raises the question of whether certain genetic interactions are particularly prone to becoming involved in incompatibilities. Intuitively, genes that interact with many others or those that evolve rapidly may be especially likely to breakdown in hybrids (*5, 6*). Indeed, the only other know hybrid incompatibility in vertebrates, the recombination hotspot regulator *prdm9*, causes hybrid sterility in multiple crosses in mice (*42*). Whether unifying molecular or evolutionary forces drive the evolution of hybrid incompatibilities will become clearer as more incompatibilities are mapped to the single-gene level.

## Acknowledgements

We thank Erin Calfee, Hunter Fraser, Bernard Kim, Mark Lipson, Priya Moorjani, Molly Przeworski, and members of the Reich, Rosenthal, and Schumer labs for helpful feedback on this work. We appreciate the help of Alejandro Moyaho-Martinez in the design of swim trials, Joseph Lim and members of the Rosenthal lab in collecting samples and Oscar Juárez-Mora for help conducting swim trials. We thank the Federal Government of Mexico for permission to collect fish. Stanford University and the Stanford Research Computing Center provided computational resources and support for this project. This work was supported by an NSF LTREB 1354172 to G. Rosenthal, funding from the Hagler Institute for advanced study to M. Schartl, and a Hanna H. Gray fellowship, L’Oreal for Women in Science grant, and NIH 1R35GM133774 grant to M. Schumer.

## Supplementary Materials

Materials and Methods

Appendix of computational commands

Table S1-S3

Figures S1-S41

Video S1

Video S2

