## Supporting Information for "Natural hybridization reveals incompatible alleles that cause melanoma in swordtail fish"

### Supplementary Online Material

Daniel L. Powell, Mateo Garcia, Mackenzie Keegan, Patrick Reilly, Kang Du, Alejandra P. Díaz-Loyo, Shreya Banerjee, Danielle Blakkan, David Reich, Peter Andolfatto, Gil Rosenthal, Manfred Schartl, Molly Schumer

### 1. Materials and Methods

#### 1.1 Analysis of spotted caudal phenotype in parental species and hybrids

To characterize phenotypic differences in the spotted caudal between *X. birchmanni*, *X. malinche*, and their naturally occurring hybrids, we began by quantifying spotting phenotypes in wild-caught males from multiple hybrid and parental populations. We focus our collections on males because the spotted caudal trait is rare in females.

1.1.1 Phenotyping methods

All fish were collected using baited minnow traps. Fish were anesthetized in MS-222 (Texas A&M IACUC protocol #2016-0190; Stanford APLAC protocol #33071) and photographed against a grid background with caudal fins spread using a Nikon d90 DSLR digital camera equipped with a macro lens. Each fish was scored for the presence of the spotted caudal pattern, invasion of melanin containing cells into the body of the fish, as well as a light colored margin surrounding the melanized area. Invasion was scored as the extension of the melanized area into caudal peduncle (i.e. anterior to the hypural plates). The presence or absence of three-dimensional growth was noted for fish that had invasion of melanin containing cells into the caudal peduncle. Melanized area, standard length, and body depth were also measured from photographs using ImageJ (*1*).

1.1.2 Spotted caudal progression methods

Given the striking differences in phenotype we observed between *X. birchmanni* and natural hybrids (Fig. 1; Fig. S2), we wanted to confirm that this was driven by genetic rather than environmental differences. We raised wild caught juveniles from the Chahuaco falls hybrid population ($n = 6$) and the Coacuilco *X. birchmanni* population ($n = 6$) under controlled laboratory conditions. We tracked changes in the melanized area over time.

We photographed hybrid and parental juvenile fish with the spotted caudal phenotype at the time of collection (December 2018) and approximately six months later (May 2019). This allowed us to quantify the expansion of melanized area in hybrid and parental individuals. We normalized measures of total melanized area and invasion area by standard length for each individual at both time points and averaged measurements taken from both sides of the fish. We then compared the expansion of the spotted caudal phenotype across groups using two sample t-tests. We defined the change in normalized melanized area as the response variable and population as the predictor variable.

The change in normalized invasion area was greater in hybrid fish from the Chahuaco falls population than in *X. birchmanni* from Coacuilco (Fig. 1; two-sample t-test: $t = 4.81$, $p = 0.0008$), a pattern that was also observed for total melanized area (two-sample t-test: $t = 2.66$, $p = 0.031$).

1.1.3 Spotted caudal frequencies in wild populations

Since a subset of hybrids with the spotted caudal trait will develop melanoma (e.g. Fig. S1), which may reduce their probability of survival, we tracked the frequency of the spotted caudal trait in juvenile and adult male individuals in three natural hybrid populations. Due to the unpredictability of progression, we were unable to determine which juveniles will eventually develop melanoma and which will not. However, we can assume that a subset of hybrid individuals with the spot will develop melanoma.

We tracked juvenile and adult male spotting frequency in two hybrid populations where melanoma occurs at high frequency, in ~30% of spotted adults (Chahuaco falls and Calnali low), and one hybrid population with low incidence of melanoma in spotted adults (8%; Aguazarca). We also collected this information in a pure *X. birchmanni* population (Coacuilco) where 0% of sampled spotted individuals had melanoma.

In each population, we collected individuals in baited minnow traps 1-3 times a year and recorded the number of juvenile and adult males with and without the spotted caudal phenotype. We used morphological characteristics to distinguish between juvenile and adult males. Male *Xiphophorus* possess a sexual organ used for sperm transfer called a gonopodium which develops at sexual maturity. We distinguished adult males from juveniles based on the completed development of the gonopodium, which is characterized by stiffening of rays 3-5 of the anal fin and presence of hooks and serrae at the distal end of these rays. Any fish showing incomplete development of the gonopodium were scored as juveniles.

We observed marked shifts between juvenile and adult male frequencies in populations with high melanoma incidence (Fig. 4). Although we do not have sufficient samples from all populations to perform the analysis on a per-year basis, we performed a Mantel-Haenszel Chi-Squared Test to test whether there were differences between years in juvenile and adult spotting frequencies. The only population in which we observe an effect of collection year is the Calnali low population ($p=0.01$; all other populations $p>0.18$).

#### 1.2 Histological and gene expression analysis of spotted caudal melanoma

##### 1.2.1 Histology of tissue from hybrid individuals

We prepared histological sections from seven wild-caught hybrid males from the Chahuaco falls population that had invasion of melanocyte containing cells into the body to evaluate them for evidence of malignancy. The peduncle and tail fin were fixed in a 4% formaldehyde solution for 24 hours at 4 °C. Subsequently, the tissues were dehydrated, embedded in paraffin and then serially sectioned at 5μm thickness. The sections were counterstained with hematoxylin and eosin.

Individuals spanned a range of externally visible invasion phenotypes; normalized invasion area in the sampled individuals was 0.025-0.16 cm (the range observed in overall population was 0-0.16 cm). However, all of the individuals evaluated histologically showed signs of an advanced malignant melanoma, including cell invasion, tissue destruction, and focal epidermal infiltration. Examples of these malignant features can be seen in Fig. 1 and Fig. S5. These results indicate that even individuals with less severe invasion phenotypes in Chahuaco falls have melanoma.

##### 1.2.2 Gene expression data collection

To ask if cancer-related genes were upregulated in individuals with invasive spotted caudal, we collected caudal fin tissue from four hybrid individuals with expanded melanized area, four with normal spots, and four without a spot. We also collected low-coverage whole genome sequencing data (see below *1.4.1 DNA extraction and Tn5 library preparation* for details) to confirm that there were not systematic difference in ancestry between groups which could confound expression analyses (Fig. S6; ANOVA $p=0.12$).

We extracted RNA and prepared mRNA-seq libraries. RNA was extracted using the Qiagen RNeasy Mini Kit (Qiagen, Valencia, CA). Briefly, dissected caudal fin tissue was combined with Buffer RLT and DTT and homogenized thoroughly using a pestle. We then followed manufacturer's instructions for RNA extraction from whole tissue, including on-column incubation with DNAse I. RNAseq libraries were prepared using the KAPA mRNA HyperPrep Kit (Roche, Palo Alto, CA) with 700-1000 ng of input RNA. Sample preparation followed manufacturer's instructions. Briefly, RNA was combined with washed mRNA Capture Beads to select molecules with polyA tails, which were chemically fragmented after incubation and washing steps. First and second strand cDNA synthesis were followed by A-tailing and ligation of unique Illumina IDT adapters. Libraries were bead purified and amplified in individual PCR reactions for 12 cycles. Amplified libraries were purified using KAPA Pure Beads and quantified using a Qubit fluorometer (Thermo Scientific, Wilmington, DE). Library size distribution and quality were visualized using Agilent 4200 Tapestation (Agilent, Santa Clara, CA). Libraries were sequenced on the Illumina NextSeq 4000 across four lanes to collect paired-end 75 basepair reads. Gene expression data is available through NCBI's sequence read archive (SRAXXXXX, pending).

1.2.3 Gene expression and enrichment analysis

Since gene expression data from hybrids represents a mixture of *birchmanni* and *malinche* derived transcripts, mapping to one of the species' reference genome could result in reference biases. To avoid issues stemming from reference bias, we instead mapped to the *X. maculatus* reference genome (*2*) using STAR (*3*), because *X. maculatus* is equally divergent from *X. birchmanni* and *X. malinche* (*4*). We used the RSEM pipeline to quantify expression of each transcript annotated in the *X. maculatus* reference genome and test for differential expression between groups (*5*).

We identified genes that were differentially expressed between individuals with melanoma and those with normal or no spots at a false discovery rate of 5%. To test for enrichment of particular functional categories, we used the biomart package in R (*6*) to retrieve gene ontology (GO) annotations and performed enrichment analysis using the GOstats package (*7*). We defined the "gene universe" as the set of genes included in the RSEM analysis. We tested for significantly over-represented GO categories using a hypergeometric test and a p-value threshold of 0.05. We used REVIGO (*8*) to remove redundant GO terms. The results of this analysis are shown in Fig. 1. The unthinned GO results can be found in Table S1.

#### 1.3 Chromosome scale *de novo* assemblies for *X. birchmanni* and *X. malinche*

Past work in the *X. birchmanni* x *X. malinche* system has relied on mapping to pseudoreferences developed from high coverage Illumina data and the *X. maculatus* reference genome (*9–11*). Concerned that the use of these references might impact the completeness of our mapping results, we constructed and annotated *de novo* assemblies for both species.

1.3.1 10X genomics and PacBio draft assemblies

High molecular weight DNA extraction and 10X Chromium library preparation were performed by the Genomic Services Lab at the HudsonAlpha Institute for Biotechnology. Briefly, high molecular weight DNA was extracted from samples were using the Genome Reagent Kit protocol following the manufacturer's instructions. DNA was diluted to 10 ng/μl, quantified using a Qubit, and then further diluted to working solutions of 0.4 ng/µl, which is the recommended concentration based on the *Xiphophorus* genome size. These working solutions were used as described in the Chromium library preparation protocol to begin the emulsion phase. The emulsion phase was broken as directed by the protocol, and bead purification was performed in 96-well plates. Final libraries were quantified using a Qubit and library size was evaluated on a bioanalyzer.

Genome assembly was performed using the supernova software v2.0.1 with the maximum reads used parameter set to 280 million and the output style specified as pseudohap; otherwise recommended parameters were used. This resulted in a draft assembly with an N50 of 12.2 Mb for *X. birchmanni* and 860 kb for *X. malinche*.

For *X. birchmanni*, which has much higher within-species polymorphism than *X. malinche* (*9*), we also collected high coverage PacBio data. High molecular weight DNA was extracted using the Genomic Tip 100/G protocol (Qiagen, Valencia, CA) following manufacturer's instructions. DNA was sent to the core facility at Mt. Sinai School of Medicine where PacBio libraries were prepared and sequenced to 75X coverage. The resulting data was assembled with *canu* (*12*) specifying a genome size of 700 Mb. The N50 of the resulting assembly was 2.0 Mb.

1.3.2 Dovetail assembly improvement

To generate a chromosome-level assembly, Chicago and Hi-C libraries were generated for both species by Dovetail (Santa Cruz, CA). Chicago libraries were prepared as described in Putnam *et al.* 2016 (*13*). Approximately 500 ng of high molecular weight DNA (mean fragment length = 100 kb) was reconstituted into chromatin *in vitro* and fixed with formaldehyde. Fixed chromatin was digested with DpnII, the 5' overhangs filled in with biotinylated nucleotides, and then free blunt ends were ligated. After ligation, crosslinks were reversed and the DNA was purified. This purified DNA was then treated to remove any biotin that was not internal to the ligated fragments. Following this step, the DNA was sheared to an average fragment size of 350 basepairs and libraries were generated using NEBNext Ultra enzymes and Illumina-compatible adapters. Biotin-containing fragments were isolated using streptavidin beads before PCR enrichment of each library. The libraries were sequenced on an Illumina HiSeq X to produce 232 million 2x150 basepair paired-end reads for *X. malinche* and 165 million 2x150 basepair paired-end reads for *X. birchmanni*, or 233X and 163X genome-wide coverage respectively.

Dovetail HiC libraries were prepared for each species as described previously (*14*). For each library, chromatin was fixed with formaldehyde in the nucleus and then extracted. Fixed chromatin was digested with DpnII, the 5' overhangs filled in with biotinylated nucleotides, and then free blunt ends were ligated as described above. After ligation, crosslinks were reversed and the DNA was purified from protein and purified DNA was treated to remove biotin not internal to the ligated fragments. Prior to library preparation with NEBNext Ultra enzymes and Illumina-compatible adapters, DNA was sheared to an average size of 350 basepairs. Streptavidin beads were used to isolate biotin-containing fragments prior to PCR enrichment of each library. The libraries were sequenced on an Illumina HiSeq X to produce 244 million 2x150 basepair paired end reads for *X. malinche* and 143 million 2x150 basepair paired end reads for *X. birchmanni*.

For each species, the 10X draft *de novo* assembly, reads from that assembly, Chicago library reads, and Dovetail Hi-C library reads were used as input data for HiRise, a software pipeline designed specifically for using proximity ligation data to scaffold genome assemblies (*13*). The assembly software uses an iterative analysis. First, shotgun and Chicago library sequences were aligned to the 10X draft *de novo* assembly using a modified version of SNAP read mapper (http://snap.cs.berkeley.edu). The separation of Chicago read pairs mapped within draft scaffolds was analyzed by HiRise to produce a likelihood model for genomic distance between read pairs, which was used to identify and break putative miss-joins, score prospective joins, and perform joins above a particular threshold. Next, Dovetail Hi-C reads were aligned and scaffolded following the same method. Finally, shotgun sequences were used to close gaps between contigs.

1.3.3 Annotation of completed assemblies

Before annotation, the initial quality of each assembly was assessed using BUSCO and the bony fishes (*Actinopterygii* odb9) database (*15-16*). We used the parameter –long so that the predicted *Actinopterygii* conserved genes would be used to train AUGUSTUS 3.2.3 (*17*). BUSCO analysis indicated high levels of completeness for both initial assemblies (*birchmanni* 97.2%; *malinche* 97.2%).

Next, repetitive elements were identified using blastx 2.2.28+ and RepeatModeler (http://www.repeatmasker.org/). The results were combined with a custom fish repeat database incorporating previously identified repeat libraries from poeciliid species (relatives of *Xiphophorus*) as well as repeat libraries from a larger fish repeat database (*18*). This custom library was transferred to RepeatMasker to identify repetitive elements in the *X. birchmanni* and *X. malinche* assemblies.

Following repeat masking, the genome assemblies were annotated using RNAseq data collected for this study as well as homology annotation and *de novo* annotation approaches. For homology annotation, we used exonerate2.2.0 (http://www.ebi.ac.uk/about/vertebrate-genomics/software/exonerate, *19*) and Genewise2-2-0 (*20*). The protein database for homology annotation contained 544,476 non-redundant proteins from Swiss-Prot (www.uniprot.org) and 13 Ensembl genomes (version 95, http://www.ensembl.org): human, mouse, sea lamprey, coelacanth, spotted gar, zebrafish, cod, tilapia, medaka, platyfish, fugu, tetraodon, and stickleback.

For RNA-seq based annotation, we collected data from $F_1$ hybrids between *X. birchmanni* and *X. malinche*. To maximize the number of expressed genes detected, we sampled eight organs: liver, testis, ovaries, brain, gills, eyes, muscle, and caudal fin. We also sampled developing embryos. We collected between 10,312,406-34,758,066 reads per sample. Reads were mapped and gene models were identified using Tophat and cufflinks 2.1.1 (*21*). In parallel, HISAT2 2.1.0, Trinity 2.4.0, and PASA 2.2.0 were used to assemble transcripts, which were then mapped to the reference genome to identify gene models (*22-23*). For *de novo* annotation, we used SNAP (http://korflab.ucdavis.edu) and GeneMark-ES (*24*).

We extracted the gene models confirmed by all of the methods described above using EVidenceModeler1.1.1 (*25*). These high quality gene models were then used to train AUGUSTUS for each species respectively. Following this training, AUGUSTUS was run using these prior models to produce a final set of gene models. From this final set of gene models, we retained high quality gene models that were present in a reference database (BUSCO, Pfam, or Swiss-prot) and/or had a predicted amino acid sequence with an Augustus score greater than 80. To assign gene symbols, we blasted each gene to Swiss-Prot (www.uniprot.org/e) using blastp2.2.28 (*26*), and assigned the gene to the best hit symbol. This resulted in 24,577 annotated genes for *X. birchmanni* and 24,416 annotated genes for *X. malinche*. Final genome assemblies and annotations are available on Dryad (Accession doi: XXXX, pending).

##### 1.4 A genome wide association scan for spotted caudal presence or absence

The spotted caudal trait in hybrids is derived from the *X. birchmanni* parental species. Within *X. birchmanni* it is polymorphic, which allowed us to search for variants associated with the spot in population samples of *X. birchmanni*. This design is advantageous because lower linkage disequilibrium (LD) within *X. birchmanni* (as compared to hybrids which have high LD due to recent admixture) allowed us to precisely map variants associated with the spot.

###### 1.4.1 DNA extraction and Tn5 library preparation

DNA was extracted using the Agencourt DNAdvance kit (Beckman Coulter, Brea, California). The protocol followed manufacturer's instructions except that we used half-reactions. Briefly, ~10 mg of tissue from the caudal fin was mixed with lysis buffer, DTT, and proteinase K and incubated for 16-18 hours at 55 ºC. Lysed tissue was mixed with 50 µl Bind 1 buffer and Bind 2 buffer to bind the DNA to magnetic beads, then washed three times with 70% ethanol. DNA was eluted in 50 µl elution buffer. Extracted DNA was quantified using the Nanodrop 8000 (Thermo Fisher Scientific, Waltham, MA).

Ten to thirty nanograms of DNA were mixed with Tn5 transposase enzyme pre-charged with adapters and incubated at 55 ºC for 7 minutes to enzymatically shear DNA. The reaction was stopped by adding 2.5 µl 0.2% SDS and incubating at 55 ºC for 7 minutes. Three microliters of sheared DNA were combined with one plate-level i5 index and one of 96 custom i7 indices in an individual PCR reaction. After amplification, 10 µl of each library were pooled with other libraries and purified using 18% SPRI beads (*27*). Libraries were quantified using a Qubit fluorimeter (Thermo Scientific, Wilmington, DE). Library size distribution and quality were visualized using an Agilent Tapestation (Agilent, Santa Clara, CA). Libraries were sequenced on the Illumina NextSeq 4000 or the Illumina HiSeq 4000 to collect paired-end 75 basepair reads and 150 basepair reads respectively. This data is available through NCBI's sequence read archive (SRAXXXXX, pending).

###### 1.4.2 Identifying variants associated with the spotted caudal in *X. birchmanni*

To identify variants associated with the spotted caudal phenotype within *X. birchmanni*, we performed low coverage whole genome sequencing (~1X average coverage) on 233 unspotted controls and 159 spotted cases from the *X. birchmanni* Coacuilco population to generate data for a genome-wide association study (GWAS).

We used the samtools-legacy program to estimate allele frequency differences between cases and controls and assign p-values to these differences between the two groups assuming a one-degree $\chi^2$ distribution (*28*; https://github.com/lh3/samtools-legacy/blob/master/samtools.1). We used this legacy version instead of the current release of samtools because it retains the ability to perform this analysis (Heng Li, personal communication). We note that were unable to use standard logistic regression approaches that explicitly accounted for population structure because of the low-coverage nature of our data. However, we did evaluate the possible impact of population structure between cases and controls using a series of analyses (see *1.4.3 Evaluating the impact of population structure on GWAS results*).

Before applying samtools-legacy, we wanted to determine its expected accuracy in simulations. We used *macs* (*29*) to generate 1 Mb diploid sequences for 392 individuals. We assumed $\theta = 0.0012$ per site (matching previous estimates for *X. birchmanni*; *9*) and used the previously inferred recombination map for *X. birchmanni* for the first megabase of chromosome

1. We input the *macs* output into seq-gen (*30*) to generate nucleotide sequences with the observed base composition of *X. birchmanni*. Next, we combined pairs of simulated haploid sequences to generate diploid individuals and generated fastq files for each individual using the program wgsim (https://github.com/lh3/wgsim). We simulated 1X coverage of this region, matching the average coverage in our data. We then randomly defined cases and controls among these 392 simulated individuals, such that the total number matched the numbers in our real data, mapped simulated reads with bwa and ran samtools-legacy to quantify allele frequency differences between the simulated cases and controls. Finally, we compared the observed allele frequency difference to the true allele frequency difference at each SNP to evaluate the accuracy of the pipeline (Fig. S23). We repeated this procedure 100 times. Across 100 simulations, the average difference between true and inferred allele frequencies was 1.2% (Fig. S23, based on the absolute value of the difference). The upper 95% quantile was 5%. These simulations suggested that application of samtools-legacy will results in accurate quantifications of allele frequency differences between cases and controls in our data.

For the real data, we followed the same procedure. Briefly, for each individual we mapped reads with bwa-mem, sorted bam files with samtools, removed reads with mapping quality scores less than 30, and ran samtools-legacy to estimate allele frequency differences between cases and controls. We only performed allele frequency analysis at high quality SNPs ascertained in high coverage whole genome sequence data from 26 individuals from the *X. birchmanni* Coacuilco population (*9*). This resulted in allele frequency estimates and associated p-values for 1,254,071 SNPs genome-wide.

To determine the appropriate genome-wide significance threshold for allele frequency differences between spotted and unspotted individuals, we used a permutation-based approach. We randomly shuffled phenotypes and repeated the genome-wide scan for allele frequency differences between cases and controls 500 times. Based on these permutations, we set our genome-wide significance threshold at $10^{-7}$ since fewer than 5% of permutations had associations at or lower than this threshold. Applying this threshold to our real data, we identify several GWAS hits associated with the spotted caudal phenotype (Fig. 2). We focus on the association found on chromosome 21 because upon closer inspection the two other regions we identified contained only a single associated SNP. Moreover, SNPs in strong LD with them ($R^2$>0.8) were not associated with the trait.

Examination of the signal on chromosome 21 revealed two distinct peaks (Fig. 2). The first of the two significant regions was centered on an ortholog of the epidermal growth factor receptor *b* (*egfrb*; Fig. S9). *egfrb* is known to have duplicated in the ancestor of some swordtail species; the two resulting copies are typically referred to as *egfrb* and *xmrk* (*31*). Indeed, our *X. birchmanni* assembly contains two genes closely related to *egfrb*, approximately 7 Mb apart. To determine whether our GWAS peak contained *egfrb* or *xmrk*, we built a maximum likelihood phylogeny of *egfrb* and *xmrk* sequences from *X. maculatus* using RAxML (*32*) as well as the orthologs we identified in *X. birchmanni* and *X. malinche*. This analysis indicated that the gene in our first GWAS peak is most closely related to the *xmrk* gene (Fig. S9).

<u>1.4.3 Evaluating the impact of population structure on GWAS results</u>

The case-control design that we use for GWAS can be susceptible to high false positive rates in the presence of population structure, since population structure can generate allele frequency differences between cases and controls that are independent of the trait of interest.

Because our data is low coverage we were not able to use a standard approach to control for population structure, but we investigated its possible impacts in the analyses described below.

To evaluate evidence for population structure correlated with the spotted caudal phenotype, we generated "pseudo-haploid" calls for each individual by sampling a read from each position in the pileup file and assigning the allele supported by that read to the individual. We used these pseudo-haploid calls to perform a PCA in all individuals using plink (*33*). We then asked whether there was a correlation between spotting phenotype and any of the first 20 PCs. The results of this analysis suggested subtle population structure between spotted and unspotted individuals (Fig. S8). We observed correlations between spotting phenotype and PC1 and PC2 loadings (both $p<0.001$), but not with any of the other top 20 PCs. Together, PC1 and PC2 explain 1.5% of the observed genome-wide SNP variation.

To understand whether population structure could be driving the signal we see at our two GWAS hits, we performed several analyses. We wanted to evaluate whether GWAS associations were substantially weakened when we accounted for population structure using pseudohaploid calls and PCs based on these calls. However, because pseudohaploid calls lower our power by discarding information, we expected to see a weaker association between the spot and the focal SNP a priori. To address this, we first evaluated the association between the spotted caudal trait and pseudohaploid calls at the peak SNPs within our two GWAS hits on chromosome 21. To do so, we generated 100 replicate sets of pseudohaploid calls at the peak SNP in each of the two associated regions on chromosome 21 and tested for an association with spotting phenotype. As expected due to reduced power, p-values at both peak SNPs rose but still remained associated with the trait (range of p-values for peak 1 from 100 replicates: $1^{-8}$ - $5^{-6}$; peak 2: $2^{-13}$ - $3^{-10}$).

Next, we repeated the analysis, this time adding PC1 and PC2 as covariates in the analysis to account for genome-wide differences between spotted and unspotted individuals. We find that the range of p-values for associations between these regions and the spotted caudal are similar to the analysis that did not account for population structure (range of p-values for peak 1 from 100 replicates: $2^{-7}$ - $3^{-6}$; peak 2: $2^{-11}$ - $1^{-8}$). This suggests that the subtle population structure we observe is not driving our GWAS results.

As a second approach to verify this, we excluded 16 outlier individuals based on their PCA loadings (PC1 > 0 and PC2 > 0.01; Fig. S8). Following this filtering the associations between phenotype and PC1 and PC2 were no longer significant ($p=0.26$ and $p=0.06$ respectively) but we observed similarly strong associations at the chromosome 21 GWAS hits (peak 1: $1^{-8}$; peak 2: $8^{-8}$).

#### 1.5 Structural differences between *X. birchmanni* and *X. malinche* in GWAS hit region

##### 1.5.1 Comparison of chromosome structure between *X. birchmanni* and *X. malinche*

To evaluate the quality of our assemblies and identify structural differences between the *de novo X. birchmanni* and *X. malinche* assemblies, we aligned all *X. birchmanni* and *X. malinche* chromosomes using MUMmer (*34*). We confirmed the presence of previously known inversions on chromosome 17 and chromosome 24 in these alignments (Fig. S24).

In addition, we identified a previously unknown inversion on chromosome 21, that overlaps a 6.5 Mb region that includes both of our GWAS hits. We aligned chromosome 21 from *X. birchmanni* and *X. malinche* to two outgroup sequences, *X. maculatus* and *X. hellerii*. These alignments demonstrated that the configuration observed in the *X. birchmanni* genome assembly is derived (Fig. S25).

Given that we had not detected this inversion in past studies (*9*, *10*), we wanted to characterize it further. To determine whether the *X. birchmanni* individual used to generate Hi-C libraries was homozygous or heterozygous for the inverted state, we used HiCExplorer (*35*) to examine the Hi-C contact map. The lack of substantial off-diagonal signal in the Hi-C contact map indicates that the Hi-C individual was likely homozygous for the inverted state (Fig. S26).

Because we do not observe high LD along this region of chromosome 21 in *X. birchmanni* population samples (Fig. S27), we hypothesized that the inversion might be fixed within *X. birchmanni*. However, this would make it surprising that we had not previously detected it. To investigate this further, we examined ancestry transitions in this region of chromosome 21 in hybrids. Notably, we observe ancestry transitions in this interval in hybrids from the Chahuaco falls population (Fig. S28), suggesting that recombination is not entirely suppressed between *X. birchmanni* and *X. malinche* haplotypes in this region. This is in contrast to previously identified fixed inversions (*9*, *10*) where no ancestry transitions are observed in hybrids (Fig. S28). Examining patterns of admixture LD decay in this region, there is some evidence for unexpected patterns compared to other regions of the genome, but the deviations are modest (Fig. S29). This suggests that many of *birchmanni* and *malinche* haplotypes in the hybrid population are co-linear in this region.

These observations are consistent with the presence of a segregating inversion in *X. birchmanni* but could also be generated by a misassembly. To confirm that this was not the case, this we turned to PacBio data that we had previously collected and assembled with *canu* (*12*). We identified a contig in our *canu* assembly (tig00017293, Fig. S26) that spanned the left inversion breakpoint. Mapping the PacBio reads to the Hi-C assembly with *minimap2* (*36*), we see evidence that the PacBio individual is heterozygous for the inversion (Fig. S30). This supports the presence of a segregating inversion in *X. birchmanni* rather than a misassembly.

Given these results, we wanted to ask about the frequency of the inversion in the *X. birchmanni* parental population. Because we have high coverage paired-end read data for dozens of *X. birchmanni* individuals (*9*), we took advantage of methods that use information about the mapping locations of paired-end read to evaluate support for the presence or absence of the chromosome 21 inversion in *X. birchmanni* samples.

Based on previous analyses that suggested better performance of the program lumpy than other paired-end read based methods for inversion detection (*37*), we decided to test the expected performance of this method on our data (*38*). We generated 25 sequences heterozygous for large inversions using the chromosome 21 sequence from the *X. birchmanni* reference genome. To do so, we randomly sampled a start coordinate and an end coordinate 6.5 Mb away, extracted this region using the program fastahack (https://github.com/ekg/fastahack), and inverted and joined this sequence with the co-linear portions of the chromosome using custom scripts (https://github.com/Schumerlab). We next generated paired-end reads using wgsim (https://github.com/lh3/wgsim) with a 0.12% polymorphism rate and the default indel rate. We chose this polymorphism rate because it matches the per basepair rate in *X. birchmanni*. The number of reads simulated resulted in 15X coverage chromosome-wide, which matches the lower end of the coverage distribution in our empirical data for *X. birchmanni*.

After generating these data, we mapped the simulated reads back to the *X. birchmanni* reference with bwa-mem (*39*) and removed duplicates and realigned indels in the bam files with picard tools and GATK respectively (*40*). Finally, we ran lumpy and evaluated the accuracy of inversion detection in each simulation. This analysis supported accurate and sensitive detection of inversions of this size with lumpy. Overall, 92% of inversions were detected in heterozygous individuals (with an "INV" or "BND" assignment from lumpy) and there were no false positives in the simulated samples. We note, however, that real data will have complexities not reflected in the simulated data, so this should be viewed as an upper bound on accuracy and lower bound on rates of false detection of structural differences.

We next analyzed the real data using the same approach. We used paired-end data from 25 unrelated moderate to high coverage individuals (15-50X) that we had previously collected (*9*) or collected for this project. We mapped each individual to the *X. birchmanni* assembly and processed files as described above. For each individual we determined whether lumpy identified an inversion ("INV") or unclassified structural rearrangement ("BND") localizing within 100 kb of each edge of the known inversion. This analysis resulted in an estimate that 24% of individuals in our *X. birchmanni* population sample harbored an inverted chromosome. We did not attempt to distinguish whether individuals were heterozygous or homozygous for the inversion. The inversion appears to be segregating in the non-spotted background because of the individuals with known phenotypes it was identified only in non-spotted individuals.

1.5.2 Considering possible technical issues generated by a segregating inversion

Concerned that the co-localization of our GWAS hits with the inversion might indicate the presence of a technical artifact, we performed a series of analyses. First, we examined the distance between the *xmrk* and *myrip* GWAS peaks and the inferred breakpoints of the segregating inversion. We found that our GWAS hits on chromosome 21 were ~400-800 kb away from the closest inversion breakpoint, suggesting that mapping difficulties around boundaries of this segregating inversion are unlikely to generate technical problems. To confirm this, we masked bases within 200 kb of the inferred inversion breakpoints and repeated the analysis. Doing so, we found that the results at peak SNPs on chromosome 21 were identical. However, to be cautious of potential issues near the inversion breakpoints, we excluded SNPs within 200 kb of inversion breakpoints in subsequent analyses.

Because the inversion is segregating in *X. birchmanni* and we happened to choose an individual with the inversion as the reference individual, we wanted to demonstrate that our results were not sensitive to this choice. As such, we repeated the GWAS analysis described above but used the *X. maculatus* (*2*) and *X. malinche* reference genomes respectively, which do not harbor an inversion in this region.

This analysis confirmed the results we obtained mapping to the *X. birchmanni* reference. We were able to replicate signals overlapping both *xmrk* and *myrip* in *X. maculatus* (at *xmrk* $p=1.8x10^{-7}$; at *myrip* $p=5x10^{-6}$). The *X. malinche* reference only contains one *egfrb* ortholog (see 1.8 Evolutionary history of genes linked to the spotted caudal). Using the *malinche* reference, we replicated signals at both this ortholog and at *myrip* ($p=8x10^{-8}$; at *myrip* $p=9x10^{-9}$).

1.5.3 Chromosome 21 is likely the X-chromosome in *X. birchmanni* and *X. malinche*

It is interesting to note that chromosome 21 is thought to be the sex chromosome in *X. birchmanni* and *X. malinche* due to homology to related species (*2*). The high frequency of structural rearrangements observed across species on this chromosome could be a result of this (Fig. S24). Interestingly, we do see much lower frequencies of the spotted caudal trait in females, both in hybrid and *X. birchmanni* populations. However, it is unlikely that this is due to males being hemizygous since our GWAS hits are outside the non-recombining region of the X chromosome (M. Schartl, personal observation). Instead, it may be due to androgen-dependent trait expression, as is common for sexually dimorphic traits in *Xiphophorus*.

#### 1.6 Admixture mapping identifies the genetic basis of melanoma in *X. birchmanni* x *X. malinche* hybrids

Having identified loci associated with the spotted caudal in *X. birchmanni*, we next set out to identify loci associated with melanoma in hybrids. To this end, we used an admixture mapping approach to detect associations between ancestry at sites across the genome and spotting and melanoma phenotypes.

##### 1.6.1 Accuracy of local ancestry inference approach using *de novo* assemblies

Previous work on these natural hybrid populations has shown that given the density of fixed ancestry informative sites between species and the time since initial admixture, local ancestry can be inferred with high accuracy (*9*, *10*, *41*). However, past work used local ancestry calling with co-linear pseudogenomes (*9*, *10*, *41*) and an ancestry inference approach called multiplexed-shotgun genotyping (*42*). This approach is predicted to have high accuracy but is computationally intensive, especially in large datasets (*33*).

We chose to implement local ancestry calling by developing a new pipeline (*43*) that accommodates *de novo* assemblies and a recently developed hidden Markov model based approach called AncestryHMM, which is efficient for large datasets (*44*). AncestryHMM takes as input data counts for each hybrid individual at ancestry informative markers as well as counts for the ancestry informative alleles in population samples of the parental species. The program also requires information about the physical location of the markers and recombination distance between markers in one of the parental genome assemblies.

We identified and validated ancestry informative markers in our *de novo* assemblies. We focused on defining ancestry informative markers in terms of the coordinates of the *X. birchmanni* genome assembly for ease of comparison to the GWAS results. As a first approach, we mapped high-coverage whole genome resequencing data from 26 *X. birchmanni* individuals and 4 *X. malinche* individuals (*9*) to both the *birchmanni* and *malinche* de novo assemblies using bwa. Although fewer *X. malinche* samples were used, the sequence diversity of *X. malinche* is approximately 4X lower than *X. birchmanni* (*9*, *10*); thus, we require fewer individuals to identify candidate ancestry informative sites. After removing duplicates and realigning indels, we excluded any read that did not map uniquely to both *de novo* assemblies to remove reads that could be impacted by reference bias. We then called and filtered variants as previously described (*9*) and identified fixed differences between *X. birchmanni* and *X. malinche*. These sites were treated as candidate ancestry informative sites for further analysis.

Next, we repeated this approach with low-coverage whole genome data for 153 parental individuals (124 from two parental *X. birchmanni* and 29 from two parental *X. malinche* populations). Using these data, we identified and removed ancestry informative sites that were not fixed in our low-coverage parental panels, defined as <98% frequency for the previously identified *X. birchmanni* or *X. malinche* allele in each population panel respectively. We also removed ancestry informative sites that were not covered in the low-coverage parental sequencing data.

We next developed a pipeline to extract read count data at ancestry informative sites for individual hybrids, format data for input into the AncestryHMM program, and run the HMM on all individuals. This pipeline is available on github (https://github.com/Schumerlab/ancestryinfer) and described in detail elsewhere (*43*). Briefly, reads were mapped to the genomes of the two parental species independently with bwa mem (*39*), and bam files were sorted with samtools (*28*). Next, reads that did not map uniquely to either parental genome were identified with ngsutils (*45*) and filtered from the two bam files for each individual. After filtering these reads, the mpileup command from bcftools was used in combination with custom scripts to generate a table of counts for each allele at all ancestry informative sites and calculate expected recombination distances between sites before applying ancestry inference using AncestryHMM (*44*). In our implementation we used a uniform recombination prior.

We tested the expected performance of this pipeline using several approaches. First, we took advantage of previously generated lab crosses where we had clear expectations for local ancestry patterns to evaluate performance. We ran the pipeline and evaluated inferred genome-wide ancestry in 200 $F_1$ hybrids. Applying a posterior probability threshold of 0.9, we found that estimated genome-wide ancestry in these individuals ranged from 49.6-50.1% *X. malinche* and genome-wide ancestry heterozygosity ranged from 99-100%, implying a low error rate (Fig. S10). Furthermore, $F_2$ individuals show expected patterns of local ancestry (Fig. S11). To further improve our ancestry inference, sites that had higher than 0.9 posterior probability for a homozygous ancestry state in $F_1$s were removed. Thus, in practice we expect that our ancestry inference accuracy will be even higher. The final number of ancestry informative sites that passed this filtering was 680,291 million.

Because lab crosses have large ancestry tracts and may represent a less challenging case of local ancestry inference than natural hybrids, we also performed simulations to evaluate performance under simulated scenarios that matched what is known about the hybrid population we focused on for admixture mapping. We developed a hybrid population simulator called *mixnmatch* (*43*) to simulate admixed individuals. Briefly, *mixnmatch* uses coalescent simulations to generate parental haplotypes and expectations about ancestry tract lengths for the simulated hybrid populations to generate hybrid chromosomes. This simulator is publicly available (https://github.com/Schumerlab/mixnmatch) and described in more detail elsewhere (*43*).

We performed two sets of simulations of a single chromosome for 100 diploid individuals. In both simulations we set the expected mixture proportion to 32% *birchmanni* and 68% *malinche*, matching the observed mixture proportions in our mapping population (see below). We simulated reads such that coverage was ~1X, as in our real data. We set the number of fixed ancestry informative sites for simulations based on observed values for chromosome 1 in our population samples of *X. birchmanni* and *X. malinche* and used the local recombination map for chromosome 1. In the first simulation, we set the number of generations since initial admixture to 45 which matches previous estimates for populations upstream of the Chahuaco falls population (*10*). Because performance could be negatively impacted if admixture is older than expected (and thus ancestry tracts are shorter), we also performed simulations doubling the time since initial admixture. Results of these simulations indicate that we expect to have high accuracy in local ancestry with our pipeline. Applying a posterior probability threshold of 0.9, average error rates were 0.05% and 0.12% in the simulations of 45 and 90 generations since initial admixture respectively (Fig. S13).

*mixnmatch* simulates hybrid chromosomes using co-linear genomes, whereas our real data originates from hybrids whose parental genomes have both small and large structural differences. To incorporate this into our simulations, we introduced structural variants into hybrid genomes at the read generation step before performing ancestry inference. To select the appropriate parameters, we examined the observed frequency and distribution of small indels detected by GATK (as described in *9*) when reads from a pure *X. malinche* individual were mapped to the *X. birchmanni* reference sequence (Fig. S12). We found that we could approximately mimic the observed frequency and size distribution with wgsim-generated reads by setting wgsim's -X parameter (the probability that an indel is extended) to 0.85, and the per site indel frequency to 0.0012. We simulated reads using this approach and then applied our ancestry inference pipeline as described above, simulating 45 generations since initial admixture.

The results of this simulation suggested that we can infer ancestry with high accuracy even when incorporating realistic rates of structural differences between species. The average error rate in this simulation was 0.09% (Fig. S13). Although these simulations do not capture all of the complexities of structural rearrangements in real genomes, paired with our results for lab-generated hybrids they suggest that our approach yields accurate local ancestry information.

1.6.2 Local ancestry analysis in natural hybrids

We collected 232 hybrids from a wild population with a high incidence of melanoma, the Chahuaco falls population. We photographed each fish and phenotyped individuals for the spotted caudal as described above (*1.1 Analysis of spotted caudal phenotype in parental species and hybrids*). For each individual, we collected low-coverage whole genome sequence data with an average of 1X coverage genome wide following the DNA extraction, library preparation and sequencing methods described above (see *1.4.1 DNA extraction and Tn5 library preparation*).

For local ancestry inference in hybrids, raw data was parsed by barcode and trimmed to remove low quality basepairs (Phred <20). To prevent long computational times, we limited the number of reads per individual to 16 million (~3X coverage genome-wide). Individuals with fewer than 300,000 reads were excluded based on simulation results which suggested that local ancestry inference is less accurate for these individuals (*41*, *43*). This resulted in the inclusion of 209 Chahuaco falls hybrids in our final analysis.

The parameters used in our HMM analysis were based on prior work on swordtail hybrid populations (*9–11*). We set the number of generations of admixture to 45, the recombination rate to an average of 1 cM/500 kb, and the per-site error rate to 0.02. To determine the appropriate priors for genome-wide mixture proportions, we initially ran the HMM with a uniform prior for the possible ancestry states. Based on this initial run, we estimated the average proportion of the genome derived from the *malinche* parent species in the Chahuaco falls population to be approximately 68% (Fig. S4). We then re-ran the HMM using these priors.

AncestryHMM outputs ancestry in the form of posterior probabilities. We converted these to hard ancestry calls by requiring a posterior probability of 90% or greater to assign a site to a given ancestry state. This threshold was chosen based on simulations which suggested that imposing this threshold results in an accurate set of hard calls (see *1.6.1 Accuracy of local ancestry inference approach using de novo assemblies*; *41*, *43*). Sites with lower than 90% posterior probability for any ancestry state were masked, as were sites that were covered in fewer than 50% of individuals in the sample. This procedure resulted in 605,745 ancestry informative sites in 209 individuals included in our final analysis. Because this number of ancestry informative markers is more than sufficient to tag all ancestry transitions in hybrids from this population, we thinned our data before performing admixture mapping analyses (see *1.6.3 Admixture mapping of two spotted caudal phenotypes*) to exclude adjacent columns with redundant information.

1.6.3 Admixture mapping of two spotted caudal phenotypes

Leveraging our local ancestry calls, we next used an admixture mapping approach to scan for associations between local ancestry and the phenotypes of interest. To map regions of the genome associated with melanoma, we focused on the phenotype in our dataset that is most tightly linked with melanoma: visible invasion of melanin containing cells into the caudal peduncle (see *1.2 Histology of tissues from hybrid individuals*). For each genomic position we compared two linear models:

model1 = glm(invasion_presence_absence ~ proportion_genome_*malinche* + body_length)

model2 = glm(invasion_presence_absence ~ ancestry_focal_site + proportion_genome_*malinche* + body_length)

We used the difference in log likelihood between these two models as a measure of the association between ancestry at a focal site and the invasion phenotype. Body length, which correlates with age (*46*, *47*), was included as a covariate since observations of lab-raised individuals indicated that the likelihood of invasion increases with age (Fig. 1).

To generate a genome-wide significance threshold, we performed 1,000 permutations. For each permutation, we jointly shuffled the invasion and body length phenotypes, and performed the genome-wide scan. To determine the appropriate significance threshold, we asked what likelihood difference threshold was required such that fewer than 50 of the 1,000 simulations contained associations that exceeded the threshold.

We identified two regions associated with the invasion phenotype, a *birchmanni*-associated quantitative trait locus (QTL) in the same region of chromosome 21 we identified previously (log likelihood difference = 15.6) and a *malinche*-associated QTL on chromosome 5 (log likelihood difference = 12.8; Fig. S31). When we repeated this procedure for the spotted caudal alone we only identified the QTL on chromosome 21 (log likelihood difference = 23; Fig. S31). This suggests that the *malinche*-associated peak is specifically associated with the melanoma phenotype.

Recent work has raised concerns about the use of permutation-based approaches for significance thresholds in the context of polygenic traits (*48*). As a result, we evaluated whether either the melanoma or spotted caudal traits were associated with genome-wide ancestry in our sample of natural hybrids. We did not see evidence of such an association, suggesting that a permutation-based approach is valid for the phenotypes we mapped (spotted caudal - Pearson's correlation coefficient: -0.10, p=0.14; invasion - Pearson's correlation coefficient: 0.0002, p=1.00).

1.6.4 Estimating the effect size of the QTL associated with the spotted caudal

We wanted to estimate the effect size of *X. birchmanni* ancestry at chromosome 21 on spotting phenotype (for similar analyses of the melanoma phenotype see *1.6.5 No evidence for involvement of a previously mapped melanoma risk region*). This is in principle straightforward since the effect size can be extracted from the logistic regressions described above. However, past work has demonstrated that effect size estimates in QTL association studies are likely to be inflated if the experiment is underpowered (*49*). In other words, conditional on observing an association in a study with low power, the estimated effect size is likely to be inflated. We thus performed approximate Bayesian computation (ABC) simulations to determine what effect sizes are consistent with our observed signal at the QTL on chromosome 21.

To simplify the parameter space explored in these simulations we made several assumptions. First, we assumed that the presence or absence of the spotted caudal is completely controlled by genetic factors derived from the *X. birchmanni* genome, with no environmental contribution. We also assumed that *X. birchmanni* ancestry in the focal region on chromosome 21 was fully dominant; this is consistent with observed phenotypes in heterozygous individuals in our mapping results (Fig. 3). Although the assumption of no environmental contribution may be incorrect, this would result in an underestimate of the true effect of *X. birchmanni* ancestry at the chromosome 21 QTL and thus be conservative.

We extracted genotypes for each Chahuaco falls individual at the peak associated marker on chromosome 21 and used these genotypes and genome-wide ancestry for each individual in simulations. For each replicate simulation, we drew the probability of being spotted as a function of *X. birchmanni* ancestry at the QTL peak as $h_{chr21}$ = random_uniform(0-1), and defined the probability of being spotted that is determined by *X. birchmanni* ancestry at other loci in the genome as $h_{other} = (1- h_{chr21})$. For each set of parameters, we generated simulated phenotypes and performed ABC simulations following the steps described below:

1) We first generated simulated phenotypes for each individual:
    a. For individuals with 1 or 2 alleles derived from *X. birchmanni* at the chromosome 21 QTL peak, we used binomial sampling with probability $h_{chr21}$ to assign individuals to the spotted or non-spotted phenotype.
    b. Individuals that were assigned an unspotted phenotype based on their ancestry at chromosome 21 could have a spot due to variance contributed by other loci in the genome. For these individuals we re-drew phenotypes using binomial sampling with probability $h_{other.}$
    c. For individuals with zero *X. birchmanni* alleles at the QTL peak, the probability of being spotted is completely determined by $h_{other.}$ We drew phenotypes for these individuals using binomial sampling with probability $h_{other.}$

2) Using these simulated phenotypes, we performed a linear regression in R for models that included genotype at the chromosome 21 QTL peak as a covariate and those that did not, as we had for the real data. We determined the log likelihood difference between these models and performed rejection sampling (with a 10% tolerance).

3) This procedure was repeated until 5,000 simulations had been accepted.

These simulations resulted in a well-resolved posterior distribution for the possible effect size of the chromosome 21 QTL on spotting phenotype. The maximum a posteriori (MAP) estimate of this distribution was 0.75 (95% confidence intervals: 0.57-0.86). These simulations suggest that the QTL on chromosome 21 explains the majority of observed variation in spotting phenotype (Fig. S32).

##### 1.6.5 No evidence for involvement of a previously mapped melanoma risk region

Artificial crosses between *Xiphophorus* species that are distantly related to *X. birchmanni* and *X. malinche* (~3 million years diverged; *4*, *50*) generate hybrids with melanoma. This melanoma can manifest on the dorsal fin, body, or caudal fin and tends to occur later in life (*51*).

Although there are a number of differences between this melanoma and that observed in *birchmanni* x *malinche* hybrids, it is striking that the *xmrk* gene appears to be involved in both (*52*). This raises the question of whether the interacting region, which maps to chromosome 5 in both cases (*53*), is the same.

The interacting region identified in the *X. hellerii* x *X. maculatus* cross is not precisely resolved, in part because mapping has relied on artificial crosses between the two species. We aligned the associated 5 Mb region from the *X. maculatus* genome to both the *X. birchmanni* and *X. malinche* assemblies using BLAST and confirmed that this region did not overlap with our admixture mapping peak and instead localized to a region at least 6 Mb away. Although published mapping efforts have not been successful in narrowing this large associated region, functional work suggests that the gene *cdkn2a/b* is the strongest candidate in this interval (*54*).

We next evaluated whether there was evidence of rearrangements that would result in closer physical linkage of the two melanoma associated regions. Pairwise alignments of chromosome 5 between *X. birchmanni, X. malinche, X. maculatus* and *X. hellerii* with MUMmer did not reveal rearrangements near *cdkn2a/b* or *cd97*. In all four assemblies, *cdkn2a/b* and *cd97* were separated by ~7 Mb (Fig. S21). Consistent with these observations, we also see no evidence for LD between these two regions in the Chahuaco falls hybrids ($R^2$=0.05), and the frequency of observed ancestry transitions in the region is also normal (Fig. S33). We did not find evidence that these genes are functionally similar in their domain structure (using CDD blast; *55*) or that they have known interactions (based on available annotations in the string database, *56*).

It is also possible that the region associated with melanoma risk in *X. hellerii* x *X. maculatus* hybrids is involved in the melanoma we observe in *X. birchmanni* x *X. malinche* hybrids but that we lack sufficient power to detect it. To evaluate this, we first asked if ancestry at the *cdkn2a/b* locus was biased toward *birchmanni* or *malinche*, which would reduce power to detect an association in our modest sample of individuals. We see no evidence of departures from expectations based on genome-wide ancestry in the region of chromosome 5 containing *cdkn2a/b* (Fig. S18). Examination of log likelihood differences in this region did not reveal any sub-significant signals near the *cdkn2a/b* region (Fig. S18).

Next, we performed simulations to ask what effect size for *cdkn2a/b* could be consistent with the observed admixture mapping signal in this region using a similar ABC approach to that described above (<u>*1.6.4 Estimating the effect size of the QTL associated with the spotted caudal*</u>). If this region is involved in melanoma but we lack the power to detect it, we would expect to accept a broad range of possible effect sizes from these simulations.

To make these simulations tractable, we made several simplifying assumptions. First, we assumed that individuals without the spotted caudal would not develop melanoma regardless of their genotype at the melanoma risk locus and that only individuals homozygous for *malinche* ancestry would develop melanoma (based on previous results from *X. hellerii* x *X. maculatus* and our results; Fig. 3, *52*). Second, we assumed that the development of melanoma was completely controlled by *X. malinche* ancestry and did not have any environmental contributions.

To perform ABC simulations, we selected the ancestry informative marker closest to the start of the *cdkn2a/b* gene (885 bp away from the gene) and extracted genotypes for all Chahuaco falls individuals at this locus and their spotted caudal phenotype. For each simulation, we drew the probability of developing melanoma as a function of *X. malinche* ancestry at *cdkn2a/b* as $h_{cdkn2a/b}$ = random_uniform(0-1) and defined the probability of developing melanoma that is determined by *X. malinche* ancestry at other loci in the genome as $h_{other} = (1- h_{cdkn2a/b})$. For each set of parameters, we performed ABC simulations following the steps described below:

1) We first generated simulated phenotypes for each individual:
   a. We set the risk of melanoma to 0 if the individual did not have the spotted caudal.
   b. For individuals with the spotted caudal and two *malinche* alleles at *cdkn2a/b*, we used binomial sampling with probability $h_{cdkn2a/b}$ to assign spotted individuals to melanoma or no-melanoma categories.
   c. Spotted individuals that were assigned a no-melanoma phenotype based on their ancestry at *cdkn2a/b* could still have melanoma due to risk contributed by other loci in the genome. For these individuals we re-drew phenotypes using binomial sampling with probability $h_{other.}$
   d. For spotted individuals with zero or one *X. malinche* alleles at *cdkn2a/b*, the probability of having melanoma in our simulations is determine entirely by $h_{other.}$ We drew phenotypes for these individuals using binomial sampling with probability $h_{other.}$

2) Using these simulated phenotypes, observed genotypes, and observed genome-wide ancestry, we performed a linear regression in R for models that included genotype at *cdkn2a/b* as a covariate and those that did not, as we had done for the real data. We determined the log likelihood difference between these models and performed rejection sampling (with a 10% tolerance).

3) This procedure was repeated until 5,000 simulations had been accepted.

These simulations resulted in well-resolved posterior distributions for possible effect size of the *cdkn2a/b* region. The posterior distribution of the effect of *cdkn2a/b* based on accepted simulations has a maximum a posteriori (MAP) estimate of 0.1 (determined by the mode, 95 confidence intervals 0.013-0.43; Fig. S18). Because the impact of this region on melanoma development is mendelian in *hellerii* x *maculatus* hybrids (*53*), these result suggest that we can reject the hypothesis that the melanoma has the same genetic basis in the two crosses. However, these simulations also highlight that we are underpowered to detect regions with weaker effects on melanoma-risk given our sample size.

We next repeated the above procedure to estimate the effect size of the melanoma risk region that we mapped in this study. We assumed melanoma risk was recessive given observed patterns in individuals heterozygous for *malinche* ancestry in this region and selected the ancestry informative marker with the highest likelihood difference from our QTL peak (at *cd97* on chromosome 5; Fig. 3, S18). Otherwise simulations were performed as described above. These simulations resulted in well-resolved posterior distributions for the impact of *malinche* ancestry at the *cd97* region on melanoma risk. The MAP estimate for the effect of the QTL region on chromosome 5 on melanoma risk is 0.5 with 95% confidence intervals ranging from 0.26-0.7. We note that these results are consistent with other loci contributing to melanoma risk, or with important environmental contributions to the development of melanoma.

Together, these analyses suggest that the melanoma hybrid incompatibility in *birchmanni* x *malinche* hybrids and *hellerii* x *maculatus* hybrids has a partially shared (*xmrk*) and partially distinct (chromosome 5 regions) genetic basis, raising questions about how it arose in both cases.

1.6.6 Complexity of admixture mapping signal at chromosome 21

In theory, admixture mapping results for spotted caudal presence and absence in Chahuaco falls hybrids should mirror results obtained from GWAS in *X. birchmanni*, assuming that the genetic basis of the spot is the same in *X. birchmanni* as it is in hybrids, as seems sensible. However, the peak in hybrids falls at 20.8 Mb, directly between the two associated regions identified in our GWAS scan (17.2 and 22.4 Mb respectively). Although this may initially seem surprising, differences between the GWAS and admixture mapping approaches could be expected to drive these differences. Specifically, recent admixture leads to high LD in hybrids (Fig. S29), a problem that is exacerbated in this region by the presence of segregating structural rearrangements in *X. birchmanni* (Fig. S24-S26). This high LD, in combination with the two nearby GWAS hits for the spotted caudal, could reasonably be expected to generate a broad signal with a peak intermediate to the previously identified GWAS peaks.

In addition, because *xmrk* is deleted in *X. malinche* (see *1.5.1 Comparison of chromosome structure between X. birchmanni and X. malinche*; *57*) we do not expect to map spotted caudal presence precisely to the *xmrk* region in hybrids. This is because our local ancestry inference method should not report ancestry within deletions, since we require that all reads included in variant counts for the HMM map uniquely to both parental species' genomes. Indeed, nearly all hybrid individuals (97%) are missing ancestry calls in the region surrounding *xmrk*.

To test our hypothesis that the two GWAS peaks identified in *X. birchmanni* could generate the intermediate peak observed in hybrid individuals, particularly in the context of missing data near the *xmrk* region and high admixture LD, we performed simulations. We note that because of unknown parameters, we simply use these simulations to understand whether the GWAS signal on chromosome 21 could generate the broad peak we observe in the admixture mapping data.

We performed simple simulations using the genotypes observed in our data and simulated phenotypes. We selected genotypes at the closest ancestry informative marker to *xmrk* (17.1 Mb) that had <50% missing data. We confirmed that this ancestry informative marker was surrounded by SNPs in high LD with *xmrk* in the *X. birchmanni* population data ($R^2$ >0.8). We extracted individual hybrid genotypes at this marker, the ancestry informative marker closest to the second GWAS peak at 22.4 Mb, and all intervening markers.

Next, we simulated phenotypes based on observed genotypes at the ancestry informative markers closest to the GWAS peaks. For simplicity we assumed that *birchmanni* ancestry in each region contributed equally to the probability of being spotted, that *birchmanni* ancestry was dominant, and that having *birchmanni* ancestry in both regions conferred a probability of being spotted of 0.75 (see *1.6.4 Estimating the effect size of the QTL associated with the spotted caudal*). We used the following procedure:

1) Based on an individual's genotype at 17.1 and 22.4 Mb, we assigned that individual a probability of being spotted. We then determined that individual's phenotype by drawing from a random binomial weighted by the probability of being spotted.

2) Using these simulated phenotypes, we ran linear models as we had done with the real data. We determined the log likelihood difference between models with and without genotype included as an explanatory variable at each of the two focal loci (17.1 Mb, 22.4 Mb) and at the intervening ancestry informative markers.

3) We repeated this procedure 1,000 times.

Based on these simulations, we asked how frequently markers intermediate to 17.1 Mb and 22.4 Mb were found to be more strongly associated with the simulated trait. We found that in 93% of simulations, an intermediate marker (between 17.2-22.3 Mb) was significantly associated with the spotted caudal. An intermediate marker was more strongly associated with the spotted caudal than the marker at 17.1 Mb in 45% of simulations and more strongly associated than the marker at 22.4 in 63% of simulations. These results suggest that both the broad QTL interval and the location of the peak are unsurprising given features of the admixture mapping data and the arrangement of *xmrk* and *myrip* along the chromosome.

1.6.7 Localization of the melanoma risk signal

The admixture mapping peak associated with risk of melanoma contained only two genes within the genome-wide significant region: *cd97* and a long-chain fatty acid transporter. Such precise localization in an admixture mapping analysis is somewhat surprising, especially given the presence of other genes nearby (i.e. 14 genes within 100 kb of these genes). However, the local recombination rate in this region is unusually high, in the upper 10% of estimated recombination rates genome-wide, likely driving this unusually high resolution.

We wanted to ensure that precise the localization of this peak was robust to bootstrapping the individuals included in our analysis. To this end, we randomly subsampled 157 individuals (representing 75% of our sample) and re-ran our admixture mapping analyses for chromosome 5. For each simulation we recorded the intervals that exceeded the genome-wide significance threshold and we repeated this procedure 100 times. In all bootstrap resamples, a single significant peak was identified and the only genes overlapping with the significant peaks were *cd97* and the long-chain fatty acid transporter *slc27a1b*. Both genes were contained within the peak in 81% of bootstrap replicates.

1.6.8 Inferences about the history of admixture in the Chahuaco Falls hybrid population

Local ancestry information can be used to understand the history of hybridization in a population. We were particularly interested in estimating the time since initial admixture in Chahuaco falls because if admixture occurred very recently, this could explain the persistence of the melanoma hybrid incompatibility in the Chahuaco falls hybrid population.

Using the decay in admixture LD in our local ancestry data, we first estimated the time since initial admixture assuming a single pulse model of admixture (*58*, *59*). However, we note that this model is likely violated in Chahuaco falls hybrids given the variation in mixture proportion observed in individuals from this population (Fig. S4; *11*). To estimate admixture linkage disequilibrium as a function of genetic distance in our data, we first thinned our ancestry data by physical distance to retain 1 marker per 50 kb and then converted the data to plink format. We used plink (*33*) to quantify admixture LD between markers, including all markers separated by fewer than 10 Mb on the same chromosome. We converted the physical distance between markers to genetic distance based on the recombination map that we had previously developed for *X. birchmanni*. Following previous work (*58*, *59*), we fit an exponential curve to the decay of admixture LD over genetic distance using:

$$E(D) = ae^{-Tx} + c$$

where D is ancestry linkage disequilibrium, T is the number of generations since admixture, x is genetic distance in Morgans, *c* is a constant describing the value to which D decays, and *a* is a coefficient. We used the non-linear least squares function in R (`nls`) to predict *a*, T, and *c* using observed D in our data and known genetic distance. We excluded genetic distances of <0.1 cM due to concerns about the accuracy of the recombination map at this scale. In the best fit model, T is estimated to be 46 ± 1 generations (Fig. S34). We emphasize that this should be viewed as a lower bound on the time since initial admixture because recent migration will introduce long ancestry tracts that bias this estimate.

##### 1.7 Gene expression analysis of genes in the melanoma risk region

*X. malinche* ancestry in the region containing *cd97* and the long-chain fatty acid transporter *slc27a1b* is associated with melanoma risk in hybrids. Both genes have nonsynonymous substitutions between *X. birchmanni* and *X. malinche* (Fig. 3; Fig. S35, S36). However, *cd97* has known roles in cancer and overexpression of the gene has been implicated in tumor metastasis (*60–62*), while previous work has not identified involvement of *slc27a1b* in cancer pathways. This suggests that *cd97* is more likely to be the relevant gene in the associated interval. Interestingly, one of the mutations that distinguishes *X. birchmanni* and *X. malinche* is an amino acid substitution in a conserved calcium binding epidermal growth factor-like binding domain (Fig. 3).

Due to the link between *cd97* expression and metastasis, we wanted to examine whether there were gene expression differences between groups. To do so, we turned to gene expression data collected for this project from hybrids with invasive spots, normal spots, and unspotted caudal fin tissue (see <u>*1.2.2 Gene expression data collection*</u>). We trimmed RNAseq reads, mapped them to the *X. maculatus* reference genome, and evaluated differential expression with RSEM as described in section <u>*1.2.3 Gene expression and enrichment analysis*</u>. We found that *cd97* was differentially expressed between individuals with melanoma compared to individuals with normal and no spots (posterior probability of differential expression 0.92; FDR correction based on genes within 100 kb of the QTL peak). Taking the set of genes genome-wide with this level of differential expression and permuting our chromosome 5 interval onto the genome 1,000 times, we find that differentially expressed genes at this level are unlikely to localize within a permutated QTL interval by chance (p-value by permutation = 0.027). We did not see evidence for differential expression of *slc27a1b* (Fig. S37), nor any other protein-coding genes within 100 kb upstream or downstream of our QTL interval. To investigate this intriguing pattern further, we sought to characterize the expression of *cd97* as it related to spotted caudal phenotype and ancestry using a variety of approaches.

<u>1.7.1 Real-time quantitative PCR to evaluate *cd97* expression</u>

To quantify expression levels of *cd97* in a larger number of individuals of both parental species and hybrids, we used a real-time quantitative PCR approach. Using the *X. birchmanni* and *X. malinche* genomes, we designed primers that amplified from a conserved region of the gene in both species (i.e. primers that did not overlap divergent or polymorphic basepairs in either species). As a housekeeping gene, we used *efa1*, which has performed well in past qPCR experiments in swordtails (*63*) and does not show differential expression between species in RNAseq data (Fig. S38), nor allele specific expression in $F_1$ hybrids (see methods below, Fig.

S38). As with *cd97*, we designed primers that amplified a region of *efa1* that is identical between species. Primer sequences can be found in Table S2.

We tested both pairs of primers for specificity and efficiency. After confirming the presence of a single PCR product for both primer pairs and verifying this product with sanger sequencing, we evaluated the performance of our primers in qPCR. To do so, a serial dilution series of *X. birchmanni*, *X. malinche*, and $F_1$ cDNA and ran these in triplicate on a BioRad CFX384 C1000 touch real-time thermocycler machine (Biorad, Hercules, CA). All plates included no-template negative controls. The per-reaction mastermix included 1 µl cDNA, 0.4 µl of the forward and reverse qPCR primers, 5 µl of Maxima SYBR Green mastermix (ThermoFisher Scientific, Waltham, MA) and 3.2 µl of nuclease free water. The cycling conditions were as follows: 95 ºC for 10 minutes, 40 cycles of 95 ºC for 30 seconds, 60 ºC for 1 minute, followed by a read step. Each run included a final melt curve.

We found that the estimated efficiency of the primers ranged between 98-108% for each primer pair and genotype Table S2. Although such differences in efficiency could impact our results if they reflected systematic biases, point estimates of efficiency can vary due to noise across runs. We found no significant differences in slope as a function of primer target or template (Fig. S39, non-significant interaction term between template and slope in a linear model). Based on this information, we proceeded with these primers to quantify relative expression of *cd97* in the caudal fin tissue of *X. birchmanni*, *X. malinche*, $F_1$ hybrids, and natural hybrids. As described in the main text, we detected high levels of expression of *cd97* in *X. malinche* compared to *X. birchmanni*, while hybrids showed *malinche*-like expression patterns. This pattern was confirmed using RNAseq data generated from several tissues (see *1.7.2 Differences in expression at cd97 between X. malinche and X. birchmanni in other tissues*).

##### 1.7.2 Differences in expression at *cd97* between *X. malinche* and *X. birchmanni* in other tissues

To ask if higher expression of *cd97* in *X. malinche* is observed in other tissues, we reanalyzed previously collected RNAseq data for *X. birchmanni* and *X. malinche* from three available tissues (SRX2436597). To minimize the impacts of reference bias, we mapped reads to the outgroup *X. maculatus* reference genome with STAR (*3*), as described in section *1.2.3 Gene expression and enrichment analysis*. We then used the RSEM pipeline (*5*) to quantify expression of each transcript annotated in the *X. maculatus* reference genome and test for differential expression of *cd97* between groups with tissue as a covariate. This analysis suggested the presence of constitutively higher expression of the *cd97* gene in *X. malinche* (Fig. S16).

##### 1.7.3 Allele specific expression in $F_1$ hybrids

We wanted to determine whether the observed changes in *cd97* expression between *X. birchmanni* and *X. malinche* could be attributed to differences that have evolved in *cis*. To investigate this, we collected stranded RNAseq data from F1 hybrids generated between *X. birchmanni* and *X. malinche* for caudal fin tissue. We used two individuals with the spotted caudal trait and two without. This data is available through NCBI's sequence read archive (SRAXXXXX, pending).

To evaluate the evidence for allele-specific expression (ASE) of genes expressed in the caudal fin, we used a custom analysis pipeline, which is available on github (https://github.com/Schumerlab/ncASE_pipeline). Briefly, we mapped RNAseq data to both the *X. birchmanni* and *X. malinche* assemblies and counted the number of SNPs that supported the *birchmanni* and *malinche* allele for each individual. We excluded SNPs where counts based on the two references differed by more than 5%, as these are likely impacted by reference bias. For each gene, we averaged counts from the two references supporting the *X. birchmanni* and *X. malinche* alleles. We then used these counts along with the total number of mapped reads as input into an inverted beta binomial test (implemented with the ibb package in R, *64*) to evaluate differential expression of the *birchmanni* and *malinche* alleles in $F_1$ hybrids. The inverted beta binomial test appropriately accounts for non-independence between counts from the same sample. We do not see evidence for ASE of *cd97* using counts generated from either the *X. birchmanni* or *X. malinche* genomes (Fig. S14). This suggests that observed differences in expression in *X. malinche* and in hybrids are driven by changes in *trans*.

1.7.4 Predicted power to detect allele specific expression

The lack of observed ASE at *cd97* in RNAseq data from F1 hybrids could indicate the absence of *cis* regulatory modifiers of gene expression or low power to detect a *cis* regulatory effect. To investigate this, we performed simulations to evaluate whether we expect to have power to detect an allele-specific difference in expression between *X. malinche* and *X. birchmanni* if the difference in expression between species was explained entirely by ASE.

Across our qPCR and RNAseq results, we observe a 1.6-3.2 fold increase in expression of *cd97* in *X. malinche*. To be conservative, we simulated a 1.6 fold change in expression between the *birchmanni* and *malinche* alleles and a total expression level that matched the observed number of reads in RNAseq data from each of the four F1 hybrids. Specifically, we determined the number of reads that mapped to the *cd97* transcript for each F1 hybrid. Next, for each individual we drew from a random binomal distribution to determine the number of reads to generate from each species' reference genome, setting the probability of drawing a *malinche* allele to generate a 1.6 fold difference in expression. Using these numbers, we generated reads from both the *birchmanni* and *malinche* version of the *cd97* transcript using the wgsim program. For the purposes of read generation, we set the within-species polymorphism rate to species-specific values (0.12% for *birchmanni* and 0.03% for *malinche*). We simulated paired-end 75 bp reads, as in our real data. With these simulated reads, we repeated the allele-specific expression analysis pipeline described above (see *1.7.3 Allele specific expression in $F_1$ hybrids*) and asked whether we were able to detect ASE using the methods we applied to the real data. We performed 100 replicates of this simulation.

Results of these simulations indicated that we expect to have excellent power to detect ASE at *cd97* given the simulated fold-change in expression and coverage (Fig. S14). This suggests that the species-level differences in expression observed are likely due to a *trans* regulatory element.

Because of the known technical challenges associated with ASE analysis (*65*), we wanted to confirm that lack of higher expression of the *X. malinche cd97* allele using pyrosequencing techniques. We designed primers using the Qiagen Pyromark software. We tested several primer sets for performance using parental cDNA samples of both species. Briefly, we extracted RNA with the RNeasy kit (Qiagen) and used the GoScript Reverse Transcription system (#A5000, Promega Corporation, Madison, WI) without modifications to synthesize cDNA from total RNA. We then used the PyroMark PCR kit (Qiagen, Germantown, MD) following manufacturer's instructions with biotinylated primers and 45 cycles of amplification. We submitted our PCR reactions to the Protein and Nucleic Acid facility (Stanford University, CA) for pyrosequencing and used the Qiagen Pyromark software to analyze our results. Based on quality tests with parental cDNA, we selected a C/T ancestry informative site at basepair 529 in the cDNA sequence that had low signal for the alternative allele (2-3%) in three parental samples of both species. We use these primers to quantify allele specific expression in caudal fin samples from four lab-generated $F_1$ hybrids with the spotted caudal phenotype. Results for ASE based on pyrosequencing were consistent with results from RNAseq data (Fig. S14).

1.7.5 Consistency of QTL mapping and ASE results

We observe striking differences in expression of *cd97* between *X. birchmanni* and *X. malinche* and hybrids. However, this difference in expression is not explained by *cis* regulatory changes. This suggests that a difference in *trans* underlies our observation of higher expression in *X. malinche* and in hybrids.

This result is somewhat puzzling because if high expression levels of *cd97* drive melanoma, we would expect that mapping results would identify the *trans* regulatory region as the genetic basis of melanoma instead of, or in addition to, the region containing *cd97*.

One possibility is that we lack power to map this additional region with our limited sample size. As a first step in investigating this, we identified three genes that are annotated as interacting with *cd97* at high-confidence in the string database for *Danio rerio* (*56*) and determined their locations in the *X. birchmanni* genome assembly. We examined these genes for evidence of sub-significant associations with melanoma, reasoning that we might see a weak signal of association in these regions. The largest likelihood difference we observed in these regions was 2.4, which is modestly higher than the genome-wide background. However, we caution that incomplete protein interaction annotations among other issues complicates the interpretation of these analyses.

An alternate interpretation is that the observed amino acid changes between species may underlie susceptibility to melanoma, explaining the observed association between the *cd97* region and melanoma, or that both expression level of *cd97* and amino acid changes contribute to melanoma risk. These possibilities seem plausible given that we observe several amino acid substitutions between species, including a non-conservative amino acid substitution in a conserved binding domain (Fig. 3). Dissecting the precise molecular interactions involving *cd97* that increase melanoma risk will be an exciting direction for future work.

1.7.6 Determining the expression level of *xmrk*

We also wanted to evaluate expression of the *xmrk* and *myrip* genes, since our GWAS analysis indicated that these regions are associated with the spotted caudal, and as a result are associated with melanoma risk. Expression levels of *myrip* were analyzed with RSEM as described previously (*1.2.3 Gene expression and enrichment analysis*). As discussed in the main text, *myrip* has very low expression in adult caudal tissue, regardless of phenotype (Fig. S15).

To evaluate the expression of *xmrk* from RNAseq data we had to use a different approach. Since *xmrk* and *egfrb* are paralogs that are only 1.1% divergent on the sequence level within *X. birchmanni*, many RNAseq reads will not map uniquely to either copy, making it difficult to accurately quantify expression of each gene.

We used a pipeline that relied on SNP differences between the two orthologs to identify reads that originated from *xmrk* (rather than *egfrb*; Fig. S40). We mapped RNAseq reads to a version of the transcriptome containing only *xmrk* and separately to a version containing only *egfrb*. We required that informative sites be detected mapping against both copies and only retained informative sites where counts did not differ by more than 5% across the two versions of the transcriptome. Using this subset of 34 informative sites, we summed *xmrk*-specific counts for each individual (based on 14-34 SNPs per individual), and re-ran RSEM. These results are shown in Fig. S15.

#### 1.8 Evolutionary history of genes linked to the spotted caudal

Past work has suggested that *xmrk* arose in the common ancestor of two out of three of the extant swordtail groups (*31*). *Xmrk* appears to be absent from the third clade, the southern swordtails, which are more deeply diverged (*57*; Fig. S19). To better understand the evolutionary history of *xmrk*, we investigated its distribution among *X. birchmanni*, *X. malinche*, and their close relative, *X. cortezi*. Importantly, *X. cortezi* shares the spotted caudal trait with *X. birchmanni*.

##### 1.8.1 Whole genome sequencing of *X. cortezi*

We sequenced three *X. cortezi* individuals with the spotted caudal to >20X coverage. Genomic libraries were prepared following Quail et. al (*66*). Briefly, approximately 500 ng of DNA was sheared to ~400 basepairs using a Covaris sonicator. To repair the sheared ends, DNA was mixed with dNTPs and T4 DNA polymerase, Klenow DNA polymerase and T4 PNK and incubated at 25 $^{o}$C for 30 minutes. After the end-repair reaction, the DNA was purified using 18% SPRI beads. A-tails were added by mixing the purified repaired DNA with dATPs and Klenow exonuclease and incubating at 37 $^{o}$C for 30 minutes. After adding the A-tails, the DNA was purified using 18% SPRI beads and an adapter ligation reaction was performed. Purified DNA was amplified using indexed primers in an individual Phusion PCR reaction; between 11-12 PCR cycles were used. After amplification, libraries were purified using 18% SPRI beads. Libraries were quantified using a Qubit fluorimeter and library size distribution and quality were visualized on an Agilent Tapestation (Agilent, Santa Clara, CA). Libraries were sequenced on the Illumina NextSeq 4000 across four lanes to collect paired-end 75 basepair reads. These datasets are available through NCBI's sequence read archive (SRAXXXXX, pending).

##### 1.8.2 Evidence for presence or absence of *xmrk* across the swordtail phylogeny

Past work focused on *xmrk* has suggested that this gene arose in the common ancestor of two clades, the platyfish and northern swordtails, and has been lost many times (*57*). We combined the data described in the previous section with existing data from other species where we had at least 20X coverage (Table S3) to examine patterns of presence and absence of *xmrk* with whole genome data.

We used a coverage-based approach to understand whether there was evidence for a deletion of *xmrk* in a given species. Results of these analyses are summarized in Fig. S19. First, we mapped reads from each individual to the *X. birchmanni* reference genome, which contains both *xmrk* and *egfrb*. In species lacking *xmrk* we expect that the majority of reads will map to *egfrb*, resulting in lower coverage at *xmrk.* This should lead to lower normalized coverage in the region surrounding *xmrk* in the *X. birchmanni* genome, compared to coverage in species with both *xmrk* and *egfrb*. We summarized local coverage as ratio of average coverage in 10 kb windows versus average coverage genome-wide. We then normalized this data using local coverage in an *X. birchmanni* individual which underwent a PCR-free library preparation (*9*). This normalization is necessary because in species with both *xmrk* and *egfrb* some reads will not map uniquely to either copy given the similarity between the two sequences (~1% diverged at the nucleotide level; Fig. S40).

Based on this analysis, we see evidence that the *xmrk* region is absent in *X. malinche* but present in *X. cortezi* (Fig. S20). We confirmed these results by mapping to the *X. malinche* reference genome (which only contains *egfrb*) and verifying that *X. malinche* individuals had half the normalized coverage of other species at *egfrb*, as expected if reads from *xmrk* in *X. birchmanni* and *X. cortezi* mis-map to *egfrb*. Together, these results indicate that *xmrk* has been lost in *X. malinche* since its divergence from *X. birchmanni*.

##### 1.9 Selection in natural populations

In natural populations where melanoma is common, we observed fewer hybrid adult males with the spotted caudal phenotype than juvenile males (Fig. 4). By contrast, this shift is not observed in *X. birchmanni* or in a hybrid population where melanoma is uncommon (Fig. 4). Among spotted hybrid and *X. birchmanni* individuals raised in the lab, they are never observed to lose their spots, and spots in both groups tend to become more expanded over time (Fig. 1; see <u>1.1 Analysis of spotted caudal phenotype in parental species and hybrids</u>). This suggests that in high melanoma populations there is a survival bias, with spotted individuals being less likely to reach adulthood (Fig. 4).

<u>1.9.1 Evaluating possible effects of population structure on juvenile-adult male frequency shifts</u>

Shifts in juvenile and adult trait frequencies could be driven by differential survival of spotted individuals, or by differential survival of individuals as a function of their genome-wide ancestry. Although we do not see evidence for an overall correlation between genome-wide ancestry and spotted caudal phenotype (Pearson's correlation coefficient: -0.10, p=0.14), a strong genome-wide shift towards *X. malinche* ancestry would be predicted to decrease the number of spotted individuals.

To investigate this, we took fin clips for a subset of juvenile males (N=48) from the Chahuaco falls population, collected genome-wide ancestry data and compared this to ancestry data collected from adult males from this population (N=209). Only juvenile and adult males that were sampled in the same years were included in this analysis (collections from 2017 and 2019). We asked whether juveniles and adults differ in genome-wide ancestry, in total and as a function of their spotting phenotype, using an analysis of variance. We found no effect of phenotype or life stage on genome-wide ancestry (life stage: mean juvenile = 0.70, mean adult = 0.68, F=1.8, p=0.18; phenotype: mean spotted = 0.68, mean non-spotted=0.7, F=1, p=0.3). We also do not see an interaction effect between phenotype and life stage on ancestry (F=0.02, p=0.9).

To more systematically evaluate ancestry changes between juveniles and adults, we calculated the change in ancestry frequency between the two groups at ancestry informative markers genome-wide. In addition to differences that result from demographic processes and from selection, some differences in ancestry are expected simply as a result of sampling two groups. To compare observed shifts in ancestry to this null expectation, we calculated average ancestry in all 257 individuals and treated these averages as the true population mean at each marker. Next, we used the random binomial function in R to generate simulated juvenile and adult ancestry frequencies at each marker given the observed average ancestry and the number of individuals in each group. We then calculated the differences between juvenile and adult ancestry frequencies in the simulated dataset in which there were no true ancestry differences. This null distribution of juvenile-adult allele frequency differences is shown in Fig. S17.

Results of this analysis suggests that genome-wide, the observed allele frequency differences between juveniles and adults are similar to what is expected from binomial sampling, although in the real data we observe a slight shift towards *X. birchmanni* ancestry (median=0.02, Fig. S17). Although changes in population structure with age (or selection on ancestry genome-wide) that generated large shifts in ancestry could generate artifacts in tests for selection, the shifts we observe here are too slight to explain the shifts in spotted caudal frequency in our data. Moreover, we find a 21% shift towards *X. birchmanni* ancestry at the chromosome 5 melanoma risk region between juveniles and adults, which is in the top 1% of frequency shifts genome-wide (Fig. S17).

Together, these results suggest that changes in genome-wide ancestry are unlikely to explain the large shifts in the frequency of the spotting phenotype that we see between juvenile and adult male individuals in populations that have a high incidence of melanoma. Instead, we propose that the change in frequency is due to viability differences between individuals with melanoma. However, we caution that we have not excluded all possible alternative explanations for this pattern (such as a correlation between spotting phenotype and another, as of yet unknown, impact on viability).

1.9.2 Inferring viability selection using sampling of juvenile and adult males

To estimate the strength of viability selection consistent with the observed shifts in spotting frequency, we used an Approximate Bayesian Computation (ABC) approach. We emphasize two limitations of this analysis. First, not all hybrid individuals with the spotted caudal will develop melanoma (only an estimated 70±13% based on lab-raised crosses). However, due to the nature of our sampling and variation in the age at which melanoma develops, we are limited to considering juvenile-adult shifts in spotting phenotype. Second, the selection coefficients inferred in our ABC simulations should not be thought of as a true selection coefficient since they correspond to viability selection through development and not to fitness.

To perform ABC simulations, we treated the observed frequency of the spotted caudal in juvenile males as an estimate of the starting frequency of the trait. We used the rbinom function in R to generate a sample of individuals (equal to the number sampled in the real data) from a binomial distribution with a mean equal to the observed spotted caudal frequency in juveniles. We next drew a viability selection coefficient from a uniform prior between 0-1. Each simulated juvenile had a 100% probability of survival if they did not have a spot and a survival probability of $1 - s$ if they did. Our summary statistic for each simulation was the proportion of adult individuals with a spot. We accepted simulations where the adult frequency fell within two standard errors of the mean observed adult frequency in the real data. We performed these simulations separately for the two populations with high rates of melanoma (Chahuaco falls and Calnali low).

These simulations yielded well-resolved posterior distributions for viability selection coefficients in each population. The MAP estimate for the Chahuaco falls population was 0.19 (95% confidence intervals: 0.05-0.44; Fig. 4). For the Calnali low population the MAP estimate was almost identical (0.2; 95% confidence intervals: 0.04-0.38; Fig. 4), suggesting that the effect of the spotted caudal on viability is similar in the two populations.

1.9.3 Frequency of the spotted caudal in hybrid populations

One obvious puzzle raised by these analyses is how a trait under strong viability selection has persisted in some hybrid populations. To generate expectations for how quickly the trait should be purged from hybrid populations, we performed simulations using the selection coefficients inferred in ABC simulations and the hybrid population simulator admix'em (*67*). We assumed that *X. birchmanni* ancestry at the spotted caudal QTL was dominant and that only individuals with the trait experienced selection. We initiated simulations by randomly sampling a total population size from a uniform distribution ranging from 100 to 4,000 diploid individuals (*10*). We treated the observed ancestry frequency in the Chahuaco falls population as the true ancestry frequency and drew a starting frequency for the spotted caudal allele from a binomial distribution with that mean. After drawing a selection coefficient from the accepted values in ABC simulations, we allowed selection to occur for 45 generations (see *1.6.8 Inferences about the history of admixture in the Chahuaco Falls hybrid population*) and tracked the change in frequency of the simulated trait over time. We repeated this procedure 100 times.

These results of these simulations suggest that the persistence of the spotted caudal at high frequency in some hybrid populations is extremely surprising. We found that under selection coefficients consistent with the shifts in juvenile-adult frequency that we observe, in all simulations the trait was driven to low frequency in simulated populations within ~20 generations (Fig. S22). This confirms our intuition that the trait is at unexpectedly high frequency given its inferred impact on viability.

Because migration can prevent purging of deleterious alleles by constantly reintroducing them, we wanted to evaluate whether plausible levels of migration could maintain the trait at observed frequencies. Indeed, as mentioned previously, the observed ancestry distribution in the Chahuaco falls population suggests ongoing migration (Fig. S4; *11*). Given the difficulties of inferring appropriate parameters for simulations of a population with high rates of continuous migration, we treated approached these simulations as a qualitative look at the possible interplay between migration and selection on the spotted caudal. We performed simulations as described above but incorporated high levels of ongoing migration that reintroduced selected alleles each generation ($m$=0.05, 0.1, and 0.2 in separate simulations respectively).

Based on these simulations, we find that the simulated trait is maintained only in cases of high levels of continuous migration paired with low hybrid population sizes (Fig. S22). This could indicate that migration rates are extremely high in hybrid populations where melanoma is common. Alternately, it may signal that the impact of the spotted caudal melanoma on fitness is less extreme than its effects on viability imply, or that some other countervailing force is maintaining it in the population.

#### 1.10 Evaluating the impact of spotted caudal melanoma on swimming performance

One possible mechanism underlying the survival difference between spotted and unspotted individuals in populations with high rates of melanoma is a direct impact of melanoma on swimming performance. Individuals with advanced melanomas often experience degradation of a fin important to swimming and the growth of three-dimensional tumors (Fig. 4). We sought to test this in the lab using swim performance trials, evaluating both escape behavior (the "fast-start" response) and swimming endurance.

1.10.1 Fast-start startle response trials

In order to evaluate the impact of melanoma on the swim escape response we measured the swim velocity of the fast-start startle response reflex. This reflex is shared across teleost fish (*68*). Caudal fin shape is known to affect fast-start velocity (*69*, *70*).

We tested fast-start velocity following Johnson et al. (*71*). Individual fish originally collected from the Chahuaco falls population (n = 37) were placed in a 16x20 cm arena filled with water to a depth of 4 cm and allowed to acclimate for five minutes. The water temperature was maintained at 22 ± 0.2°C throughout the trial. After acclimation, we elicited the fast-start response by dropping a 2 cm glass marble from a height of 20 cm into the bottom left quadrant of the tank at a time point when the fish was not in that quadrant, was not moving, and was also at least two body lengths from any side of the tank. We recorded fast-start response using a high-speed digital video camera (Casio Exilim Pro EX- F1, Casio Computer Co., Tokyo, Japan) at 300 frames per second. Each fish was recorded three times and we used the average distance traveled in our analyses.

To standardize measures of fast-start velocity, we used ImageJ (*1*) to identify the midpoint of the fish's body length, which we used as a reference point. This coordinate was defined as the midpoint between the anterior most point of the fish and the end of the caudal peduncle. We then measured the distance traveled by tracking the path of the fish through individual frames, starting with the frame that preceded the introduction of the fast start stimulus through the following 15 frames. This corresponded to the distance the fish traveled over 53 milliseconds.

Because startle response trials included individuals with a variety of phenotypes besides the spotted caudal, we phenotyped individuals as previously described (*1.1 Analysis of spotted caudal phenotype in parental species and hybrids*) and included other phenotypes as covariates. We used the linear model function in R to analyze the distance an individual moved in 53 milliseconds after being startled as a function of invasion area, the presence or absence of 3-dimensional melanoma growth (e.g. Fig. 4), body length, sword length, and sex. Based on this analysis, we saw an effect of three-dimensional growth ($t=-2.6$, $p=0.014$) but no effect of invasion area.

1.10.2 Swimming exhaustion trials

We tested for evidence of differences in physiological swimming capacity between hybrid individuals with and without invasive melanoma patterns. We tested the ability of individuals to swim against a controlled current (n = 27). The experimental setup consisted of a rectangular tank (21 x 9 x 7 cm) covered with soft netting to catch the fish once it was exhausted. A mirror (9 x 7 cm) was attached to one side of the tank. In general, social fish are motivated to swim to exhaustion against a flow of water towards their reflection. Water flow was controlled with an attached aquarium pump positioned at the middle of the mirror (Micro Multifunction Pump, Moon's Aquariums). The swim cage was placed inside a 40L acrylic aquarium; the height of the water column (6 cm) was sufficient to allow the fish to swim without permitting it to escape from the swim cage

Before each trial, we covered the mirror with a piece of opaque acrylic (8.5 x 7 cm). We then placed the focal individual in the experimental tank to allow it to acclimatize. Ten minutes later, we removed the opaque acrylic and allowed the fish to see its mirror image for one minute. All tested fish responded to their mirror image by approaching it. After this exposure, we remotely started the aquarium pump which was set to exert a standardized rate of water flow.

Each trial was filmed and ended when 30 minutes had elapsed. After the trial, each fish was photographed on both sides with a scale reference. We used these photographs to measure the individual's standard length, sword length, body area, invasion area and presence/absence of three dimensional tumor growth using ImageJ (*1*).

Swim performance videos were scored for the frequency and duration of periods of resting and swimming. A fish was scored as resting if it was within one body length of the far end of the trial chamber (i.e furthest zone from the mirror and waterflow source). To analyze the data, we applied a general linear model with time spent in the zone furthest from the pump as the dependent variable and standard length, sex, sword length, and the presence or absence of three-dimensional growth as independent variables. This represents a model similar to those we used in the fast-start trials described above. We did not see a significant effect any independent variable on time spent resting; invasion ($t=-1.74$, $p=0.0962$; Fig. S41), three-dimensional melanoma growth ($t=0.220$, $p=0.8281$), and sword extension ($t=-0.319$, $p=.7527$) . Because few fish experienced exhaustion during trials, we speculate that more demanding swim performance trials might uncover differences between groups. However, we did not perform such tests as fish are unlikely to experience stronger flow rates in their natural environments

#### 2. Appendix of representative commands

*Note: scripts used here that are not part of pre-existing programs are available at [https://github.com/Schumerlab](https://github.com/Schumerlab)*

1.2.3 Gene expression and enrichment analysis

##### RSEM

```
rsem-prepare-reference --gtf Xiphophorus_maculatus_LG.Xipmac4.4.2.81.gtf --star --
star-path $STAR xma_washu_4.4.2-jhp_0.1_combined-unplaced-mito.fa xmac_ref

rsem-calculate-expression --strandedness reverse --star --fragment-length-mean 300 --
fragment-length-sd 50 --star-path $STAR --calc-ci --ci-memory 1024 -p 10 --star-
gzipped-read-file Coac_Wt_S19_bothlanes.fq.gz xmac_ref Coac_Wt_S19

rsem-generate-data-matrix $expsc1 $expsc2 $expsc3 $expsc4 $sc1 $sc2 $sc3 $sc4 $wt1
$wt2 $wt3 $wt4 > CHAF_samples_exp_sc_wt_RSEM_normalized_data_matrix.txt

rsem-run-ebseq CHAF_samples_exp_sc_wt_RSEM_normalized_data_matrix.txt 4,4,4
CHAF_samples_exp_sc_wt_RSEM_normalized_data_matrix.txt.results

rsem-control-fdr CHAF_samples_exp_sc_wt_RSEM_normalized_data_matrix.txt.results 0.05
CHAF_samples_exp_sc_wt_RSEM_normalized_data_matrix.txt.results.fdr
```

##### Enrichment analysis (in R)

```
library("GOstats")
library("GSEABase")
library("biomaRt")

mart <- useMart(biomart = "ensembl", dataset = "xmaculatus_gene_ensembl")

rsemanalysis<-
read.csv(file="~/Data/Chaf_sample_RSEM_matrix.results",sep="\t",head=TRUE)

results <- getBM(attributes =
c("go_id","external_gene_name","ensembl_gene_id","kegg_enzyme"),
filters=c("ensembl_gene_id"),values=as.character(c(rownames(rsemanalysis))), mart =
mart)

gene_universe<-subset(results,nchar(results$go_id)>0 &
nchar(results$external_gene_name) >0)
gene_universe$ensembl_id<-gene_universe[,3]
gene_universe[,3]<-as.numeric(as.factor(gene_universe[,3]))
gene_universe$Evidence<-rep("ISA",length(gene_universe[,3]))
colnames(gene_universe)<-
c("frame.go_id","frame.gene_name","frame.gene_id","frame.KEGG","frame.ensembl","frame.
Evidence")

goframeData =
data.frame(gene_universe$frame.go_id,gene_universe$frame.Evidence,gene_universe$frame.
gene_id)
goFrame=GOFrame(goframeData,organism="Xiphophorus")
goAllFrame=GOAllFrame(goFrame)
gsc <- GeneSetCollection(goAllFrame, setType = GOCollection())

universe = goframeData$gene_universe.frame.gene_id
```

```
rsemsig<-read.csv(file="~
/Data/Chaf_sample_RSEM_matrix.results.fdr",sep="\t",head=TRUE)
rsemsig<-subset(rsemsig,rsemsig$MAP=="Pattern4" | rsemsig$MAP=="Pattern5")

genes_sig<-row.names(rsemsig)
genes_match<-gene_universe[gene_universe$frame.ensembl %in% genes_sig,]
genes_match_sig = genes_match$frame.gene_id

params <- GSEAGOHyperGParams(name="Xiphophorus maculatus genes",
                              geneSetCollection=gsc,
                              geneIds = genes_match_sig,
                              universeGeneIds = universe,
                              ontology = "BP",
                              pvalueCutoff = 0.05,
                              conditional = FALSE,
                              testDirection = "over")

OverBP <- hyperGTest(params)

results_OverBP<-summary(OverBP)
```

##### 1.3.1 10X genomics and PacBio draft assemblies

###### 10X genomics

```
supernova run --id Xbirchmanni_10X_ref --fastqs
/home/ms857/data/Xbirchmanni_10Xchromium_Hudsonalpha_July2018_raw_data/ --description
run1 --maxreads 280000000

supernova mkoutput --
asmdir=/home/ms857/data/Xbirchmanni_10Xchromium_Hudsonalpha_July2018_raw_data/Xbirchma
nni_10X_ref/outs/assembly --outprefix=Xbirchmanni_10X_assembly --style=pseudohap
```

###### PacBio

```
canu -p Xbir_Canu_v1 -d Xbir_canu_v1 genomeSize=750m -pacbio-raw
Xbir_combined_pacbio_subreads.fastq.gz cnsErrorRate=0.25 -s xbir_canu_v1.spec
```

##### 1.3.3 Annotation of completed assemblies

###### BUSCO

```
BUSCO.py -c 30 -i ./assembly -o busco -l ~/data/actinopterygii_odb9/ -m geno –long
```

###### RepeatModeler and RepeatMasker

```
RepeatModeler -engine ncbi -pa 30 -database assembly.database

RepeatMasker -a -pa 30 -lib  ~/data/fishrepeats_from_Kang -dir . ./assembly
```

###### Gene models

```
nohup exonerate $_ localhost:12886 --model p2g --softmasktarget 1 --geneseed 250 --
showtargetgff yes --showvulgar no --showalignment yes --percent 30 --maxintron 200000
--bestn 3 >$_.exonerate

genblasta_v1.0.4_linux_x86_64 -q $_ -t ./2.repeatmodel_mask/assembly.masked -e 1e-5  -
c 0.6 -o $name.out

genewise fname.p.fa fname.d.fa -quiet -$strand -pseudo -genesf
```

```
hisat2 -p 16 -x genome.index -1 $left -2 $right -S hisat.sam >log.hisat 2

Trinity --genome_guided_bam ../hisat_stringite/hisat.sam.sorted.bam --
genome_guided_max_intron 10000 --max_memory 30G --CPU 10

Launch_PASA_pipeline.pl -c alignAssembly.config -C -R -g
./2.repeatmodel_mask/assembly.hm -t trinity_out_dir/Trinity-GG.fasta.clean -T -u
trinity_out_dir/Trinity-GG.fasta --ALIGNERS blat,gmap --CPU 2

snap -gff -quiet training/my-genome.hmm ./assembly >snap.gff

gmes_petap.pl --cores 15 --ET hisat.sam.sorted.bam.hint --sequence ./assembly

augustus --species=annopipe190909 --hintsfile=$_.hint --
extrinsicCfgFile=extrinsic.kang.cfg.RM  $_ --gff3=on > $_.gff3
```

#### 1.4.2 Identifying variants associated with the spotted caudal in *X. birchmanni*

##### Read mapping and filtering

```
bwa mem -M -R $RG $genome -t 3 $read1 $read2 > $sam

samtools fixmate -O bam $sam $bam

samtools sort $bam -o $sorted

samtools view -b -q 30 $sorted > $unique

samtools index $unique
```

##### Case control GWAS

```
$samtools_legacy_path/samtools-vlegacy mpileup -gR -r $scaff -f $genome1 $controls
$cases | $samtools_legacy_path/bcftools-vlegacy view -I -vcg -1 $num_controls - > $vcf
```

#### 1.5.1 Comparison of chromosome structure between *X. birchmanni* and *X. malinche*

##### MUMmer commands

```
nucmer ScHyZ96-1255-HRSCAF-1332.fa ScyDAA6-7-HRSCAF-50.fa -l 100 -c 200 --
prefix=ScHyZ96-1255-HRSCAF-1332_ScyDAA6-7-HRSCAF-50

delta-filter -m ScHyZ96-1255-HRSCAF-1332_ScyDAA6-7-HRSCAF-50.delta > ScHyZ96-1255-
HRSCAF-1332_ScyDAA6-7-HRSCAF-50.delta.m

mummerplot -large -layout ScHyZ96-1255-HRSCAF-1332_ScyDAA6-7-HRSCAF-50.delta.m --png -
p ScHyZ96-1255-HRSCAF-1332_ScyDAA6-7-HRSCAF-50
```

#### 1.6.2 Local ancestry analysis in natural hybrids

```
perl Ancestry_HMM_parallel_v5.pl hmm_configuration_file_CHAF.cfg
```

*Note: pipeline available at https://github.com/Schumerlab/ancestryinfer*

#### 1.6.3 Admixture mapping of two spotted caudal phenotypes

```
Rscript perform_glm_admixture_mapping_v3_binomialtrait.R $genotypes_file
$hybrid_index_file $phenotypes_file $focal_column $outfile_tag
```

#### 1.6.8 Inferences about the history of admixture in the Chahuaco Falls hybrid population

```
perl thin_genotypes_by_physical_distance.pl $genotypes 50000 $home_bin

perl convert_msg_genotypes_to_plink.pl $genotypes_thinned

plink --file $genotypes_thinned --recode --tab --geno 0.2 --maf 0.01 --allow-extra-chr
--out $genotypes_filtered

plink --file $genotypes_filtered --ld-window-kb 10000 -r2 d --hardy --hwe 0.000001 --
out CHAF_ld --ld-window-r2 0 --allow-extra-chr

Rscript convert_plink_LD_genetic_distance_xbir10x.R CHAF_ld.ld $focal_chrom
```

#### 1.7 Gene expression analysis of genes in the melanoma risk region

```
perl grep_list.pl surrounding_100kb_list Chaf_samples_RSEM_matrix.txt
surrounding_100kb_subset_Chaf_sample_RSEM_matrix.txt 1

rsem-run-ebseq surrounding_100kb_subset_Chaf_sample_RSEM_matrix.txt 4,4,4
surrounding_100kb_subset_Chaf_sample_RSEM_matrix.txt.results
```

#### 1.7.3 Allele specific expression in F1 hybrids

```
samtools mpileup -g --ignore-RG -f mal.ref.transcripts.fa $F1-1_maptomal_bam
| bcftools call -mO z -o $F1-1_maptomal_vcf

samtools mpileup -g --ignore-RG -f bir.ref.transcripts.fa $F1-1_maptobir_bam
| bcftools call -mO z -o $F1-1_maptobir_vcf

perl samtools_vcf_to_ASE_counts.pl $F1-1_maptomal_vcf
perl samtools_vcf_to_ASE_counts.pl $F1-1_maptobir_vcf

Rscript ncASE_pipeline_cmd.R $F1-1_maptomal_vcf $F1-1_maptobir_vcf $F1-2_maptomal_vcf
$F1-2_maptobir_vcf $F1-3_maptomal_vcf $F1-3_maptobir_vcf $F1-4_maptomal_vcf $F1-
4_maptobir_vcf F1_individuals_results_ncASE_counts
```

*Note: pipeline available at https://github.com/Schumerlab/ncASE_pipeline*

#### 3. Supplementary Figures

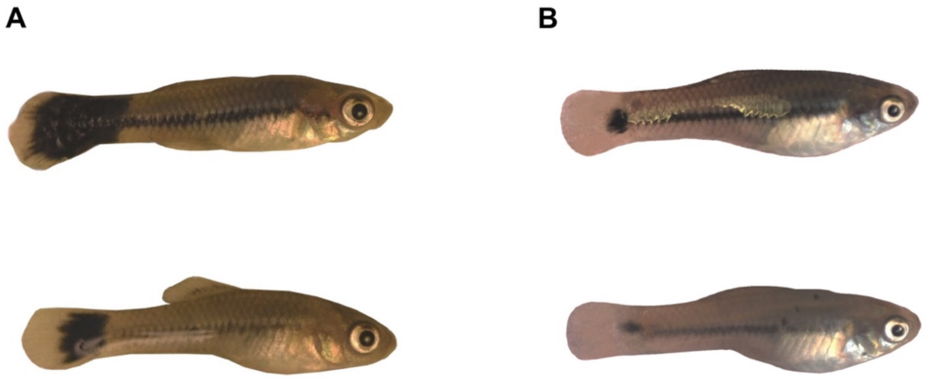

**Fig. S1.** Example spotted caudal phenotypes observed in lab reared hybrid (**A**) and pure *X. birchmanni* (**B**) juveniles. Hybrid individuals with the spotted caudal pattern sometimes develop melanoma before reaching adulthood as can be seen in the two individuals shown in **A**. In contrast, melanoma is extremely rare in *X. birchmanni*, even in older individuals (>2 years).

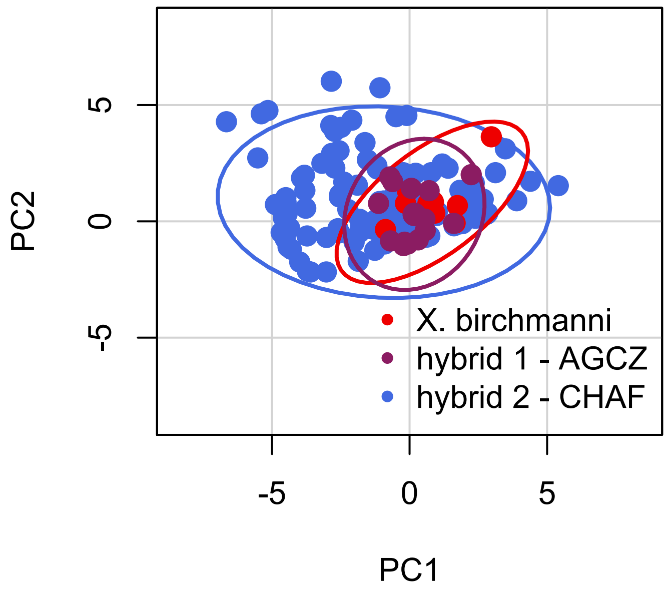

**Fig. S2.** Principal component analysis of different features of the spotted caudal phenotype. This analysis shows that there is greater variation in PC2 in individuals from the Chahuaco falls hybrid population (CHAF) than in *X. birchmanni*. PC2 is correlated with normalized spot area and invasion area. By contrast, individuals in the Aguazarca (AGCZ) hybrid population where melanoma is rare more closely resemble *X. birchmanni* phenotypes. Phenotypes included in this analysis were spot area within the caudal fin, spot area outside of the caudal fin, body length, 3D tumor growth (see Fig. 4), and whether individuals had a pale margin around the spot.

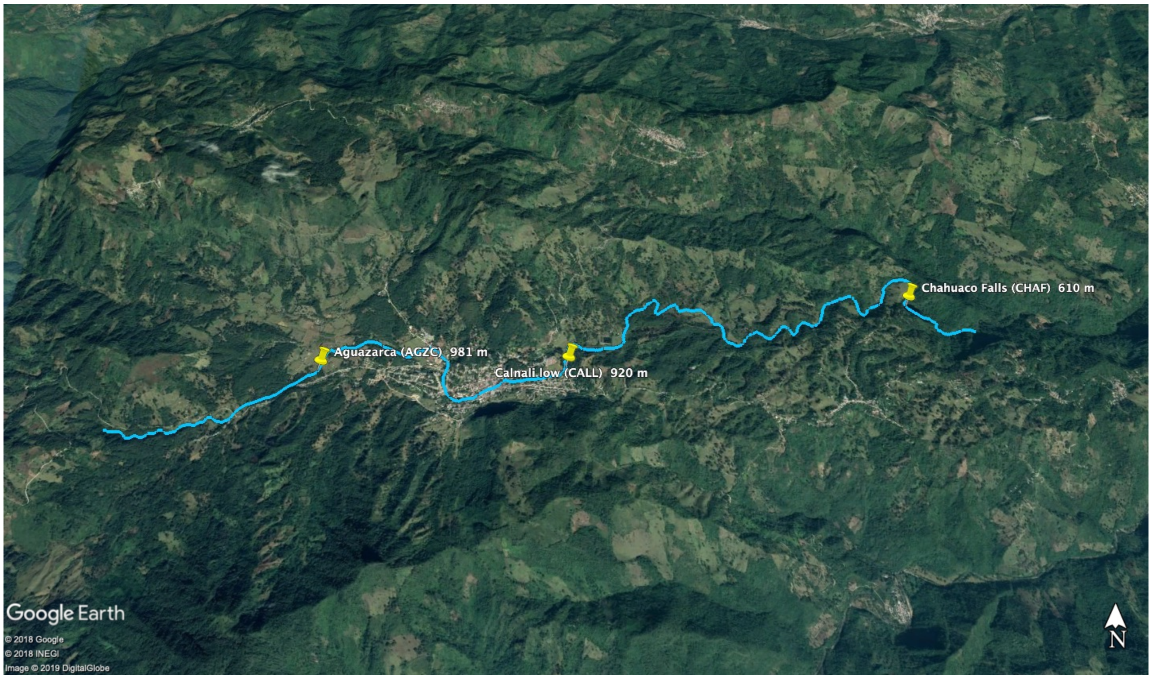

**Fig. S3.** Map of hybrid populations sampled for this study. All populations included in this study are found along the Río Calnali in the Sierra Madre Oriental of central Mexico. This image is modified from google earth.

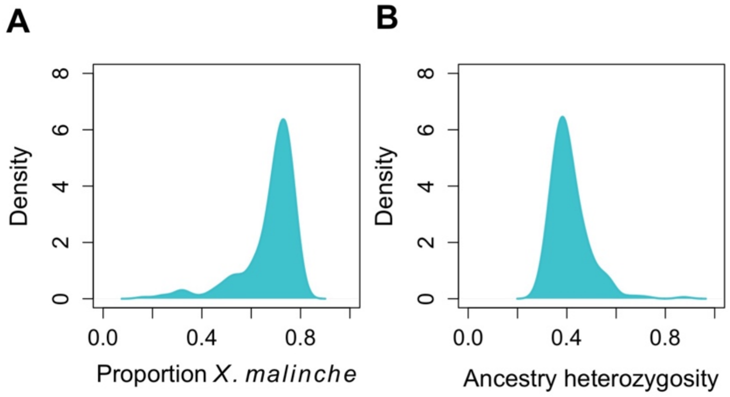

**Fig. S4.** Distribution of genome-wide admixture proportion (**A**) and the proportion of sites in an individual that are heterozygous for *X. birchmanni* and *X. malinche* ancestry (**B**) among 209 hybrid individuals collected from the Chahuaco Falls population.

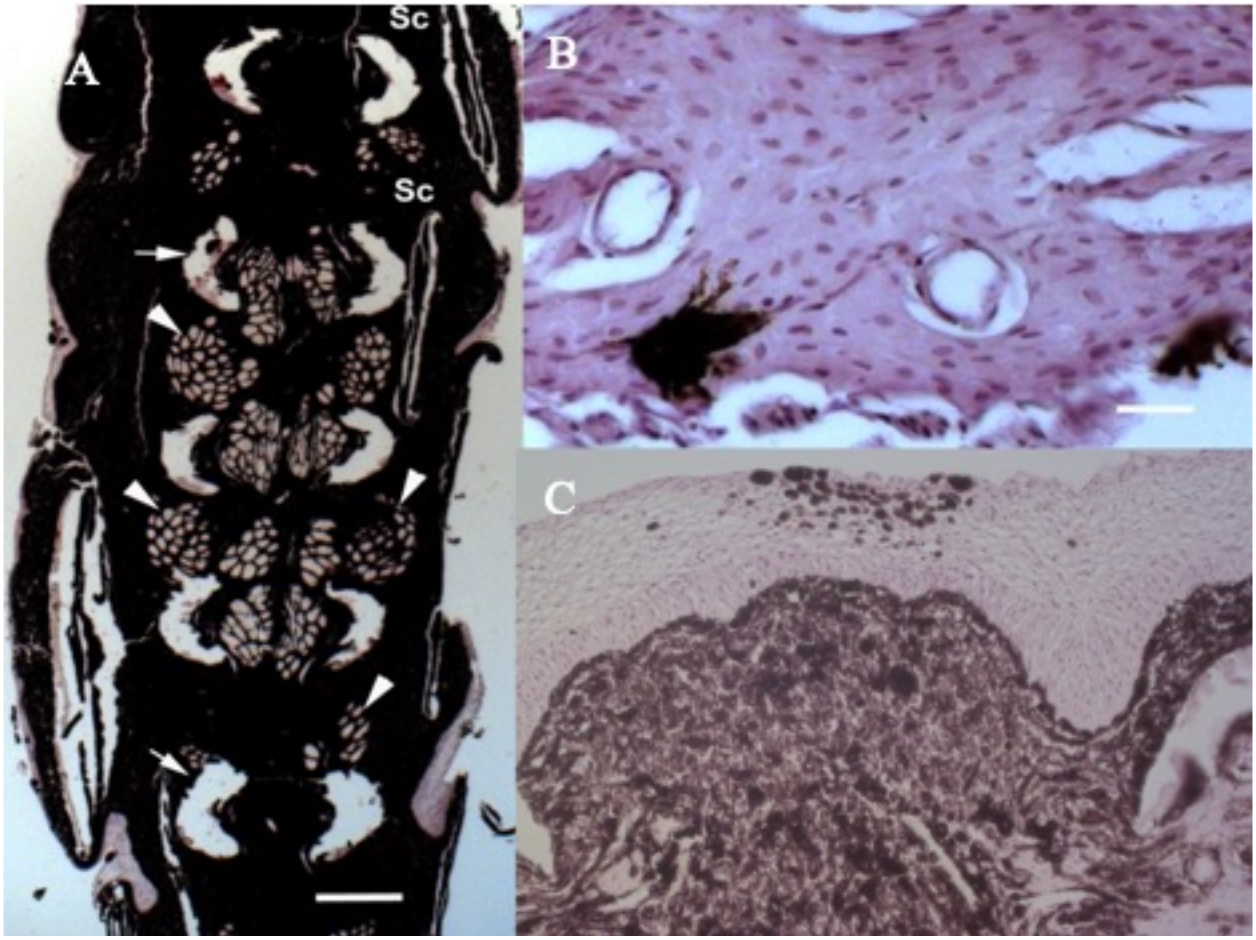

**Fig. S5.** Histology results from individuals with malignant melanoma from the Chahauco falls hybrid population. **A**) Transversal section of the distal caudal peduncle. The melanoma has replaced all tissues, and only small remnants of muscle bundles (arrow heads) are visible. Bones and cartilage of the peduncle are totally degenerated (arrows; sc – locations of scales). The white scale bar represents 300 µm. **B**) Nest of melanoma cells in the vertebral cartilage of the proximal peduncle. Here the primary tumor is not as space-filling as in the distal peduncle. Isolated melanoma cells indicate metastasis. The white scale bar represents 20 µm. **C**) Isolated nest of melanoma cells in the epidermis, with no connection to the primary tumor in the extra-epidermal compartment, indicating metastasis. Note the hyperplastic epidermis.

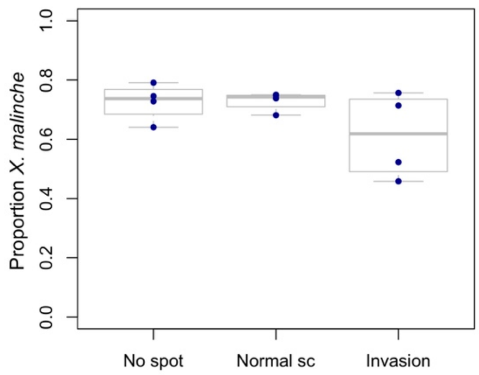

**Fig. S6.** Genome-wide ancestry of hybrid individuals from the Chahuaco Falls population used to generate RNAseq data for different spotted caudal phenotypes.

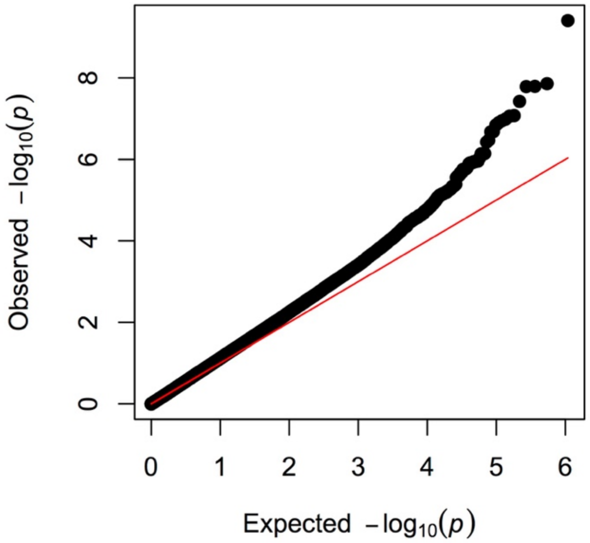

**Fig. S7.** A quantile-quantile plot of genome-wide p-values generated by case-control GWAS analysis comparing spotted and non-spotted individuals. Divergence from expectations (red line) occurs at lower p-value quantiles but matches expectations at higher p-values.

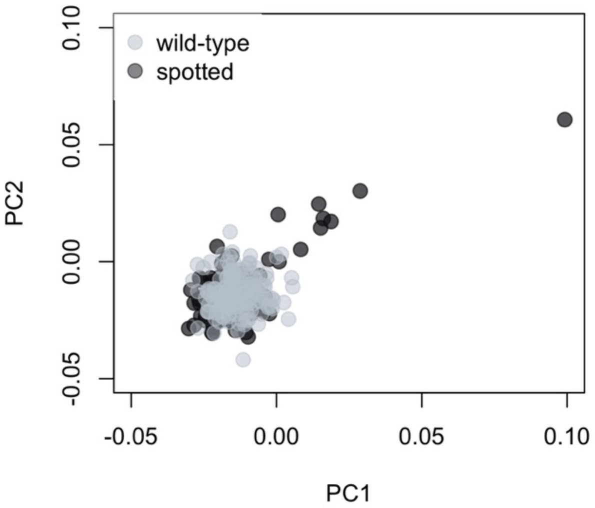

**Fig. S8.** Principle Component Analysis (PCA) identifies subtle genetic differentiation between unspotted (gray) and spotted (black) *X. birchmanni* individuals collected from the Coacuilco population. Out of the top 20 PCs, PC1 and PC2 were the only principal components correlated with phenotype (both $p<0.001$). Together, PC1 and PC2 account for approximately 1.5% of the variation in the data. See section *1.4.3 Evaluating the impact of population structure on GWAS results* for details of the PCA analysis.

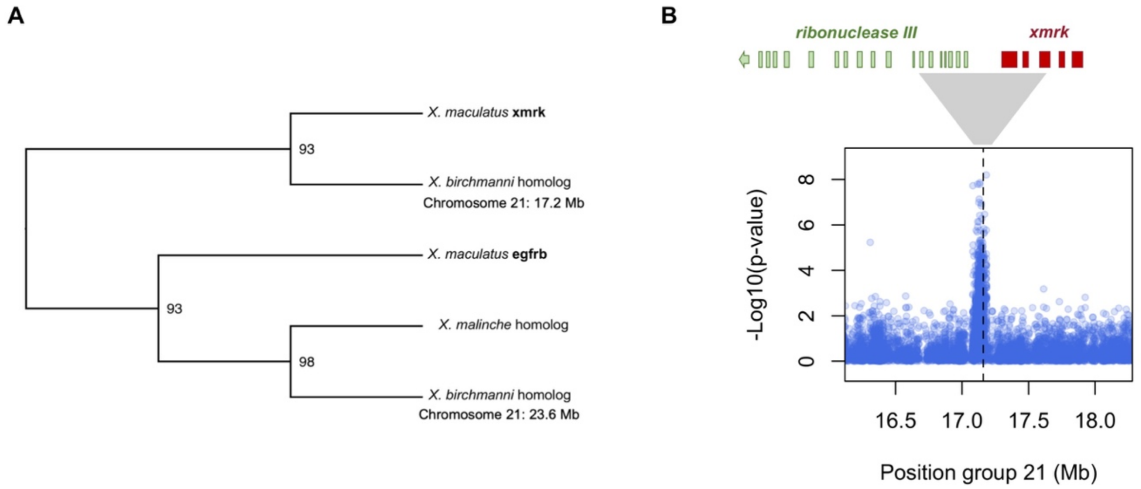

**Fig. S9. A**) Maximum likelihood phylogeny generated with RAxML shows the relationship between homologs of the epidermal growth factor b gene (*egfrb*) and its duplicate, *xmrk,* that were identified in the *X. birchmanni* and *X. malinche* genomes. Based on clustering with the well-characterized *X. maculatus* copies of these genes we concluded that the gene that falls within our GWAS hit for the spotted caudal is the *xmrk* gene. Numbers on the nodes of the phylogeny indicate the proportion of rapid bootstraps that supported the grouping. **B**) Close-up figure of the GWAS peak associated with *xmrk*. Gene models in the inset above reflect the approximate locations of exons belonging to the two genes overlapping this GWAS peak, *xmrk* (red) and ribonuclease 3 (green; the locations of remaining of the 10 exons of the ribonuclease 3 gene are indicated with the green arrow).

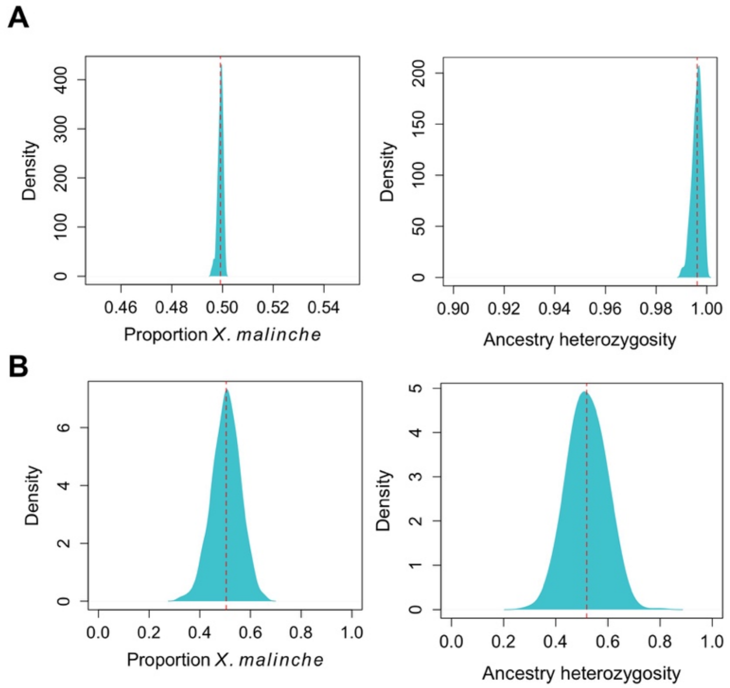

**Fig. S10.** Patterns of inferred ancestry in artificial hybrids. Inferred genome-wide ancestry (**A** – left) and heterozygosity at ancestry informative sites in $F_1$ hybrids (**A** – right) are consistent with a very low error rate in ancestry inference. Results for $F_2$ hybrids (**B**) also show expected patterns of genome-wide ancestry (left) and ancestry heterozygosity (right).

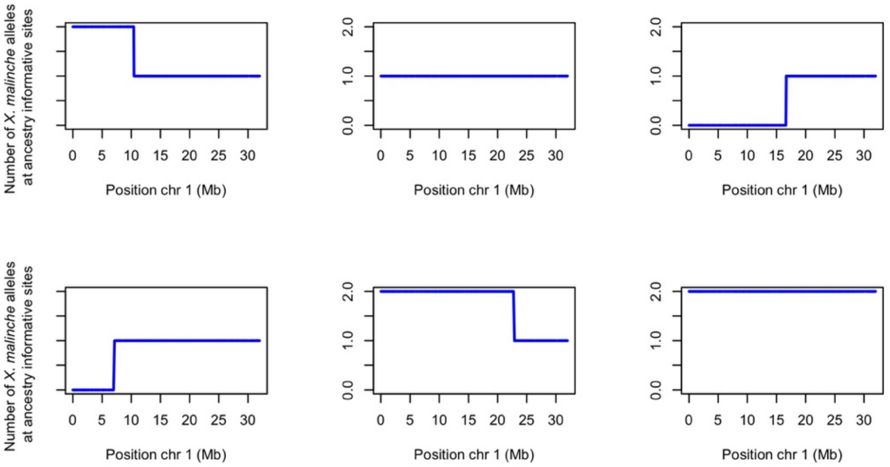

**Fig. S11.** Example results of local ancestry inference in $F_2$ hybrids. Inferred ancestry on chromosome 1 is plotted here for the first six hybrids in our dataset. Patterns of ancestry and the number of ancestry transitions mirror expectations for an $F_2$ intercross.

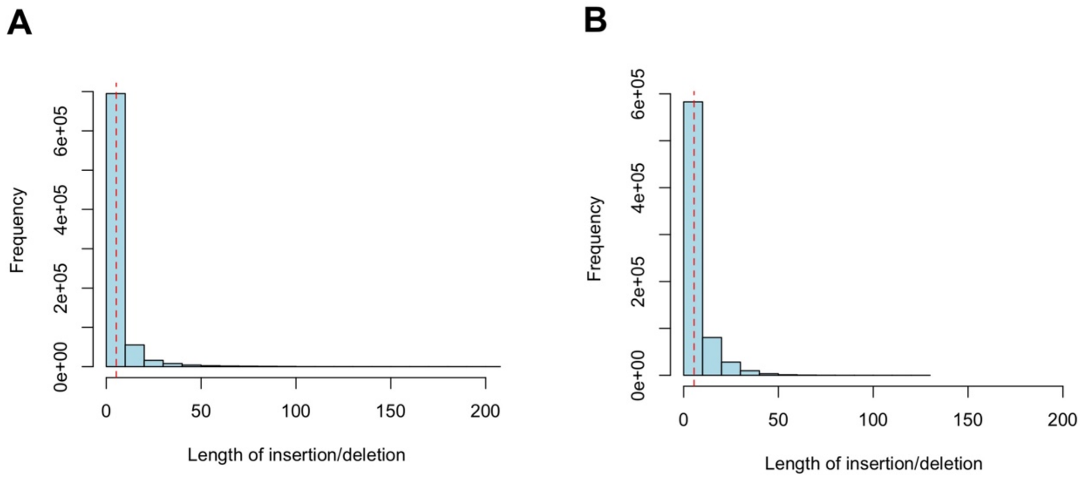

**Fig. S12.** Observed length distribution of small indels between *X. birchmanni* and *X. malinche* compared to the distribution used in simulations. **A**) Observed lengths of indels identified when mapping high coverage Illumina data from *X. malinche* to the *X. birchmanni* reference genome. **B**) Simulated indel distribution using *wgsim*. We modified the indel extension parameter in *wgsim* to roughly match the indel distribution observed our data. Dashed red lines indicate the mean length.

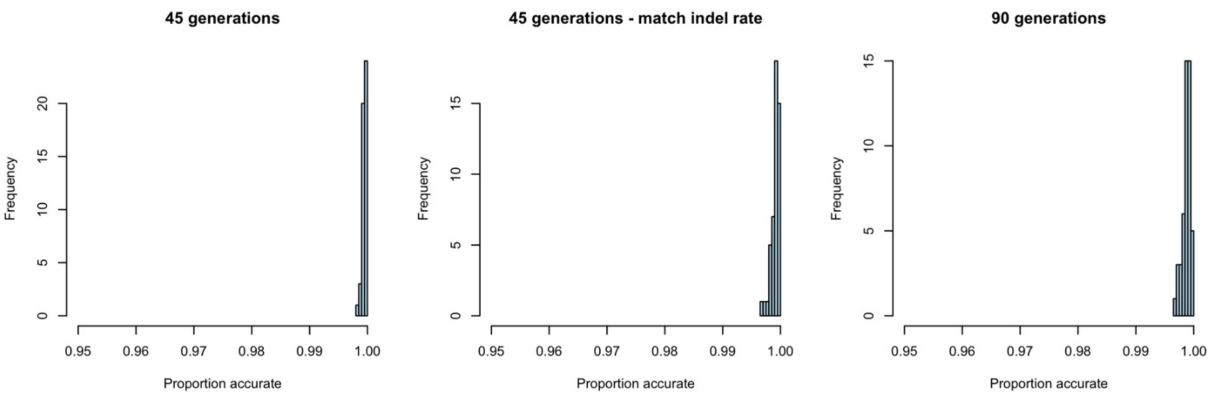

**Fig. S13.** Results of simulations evaluating predicted accuracy of our local ancestry inference approach under different scenarios. **A**) Simulations matching the observed mixture proportions for the Chahuaco falls populations and simulating 45 generations of admixture (consistent with estimates for this population; *1*) indicate that we expect to have high accuracy in this demographic scenario. Shown here is the distribution of individual level accuracy in 100 simulated individuals. **B**) We repeated these simulations with the observed indel rate and length distribution between species (Fig. S12). We found that this does not have a strong impact on accuracy (see Supporting Information *1.6.1 Accuracy of local ancestry inference approach using de novo assemblies*). **C**) Time since admixture can also impact accuracy since ancestry tracts become smaller with an increasing number of generations since initial hybridization. We repeated simulations doubling the time since initial admixture and found that accuracy was still high under this scenario.

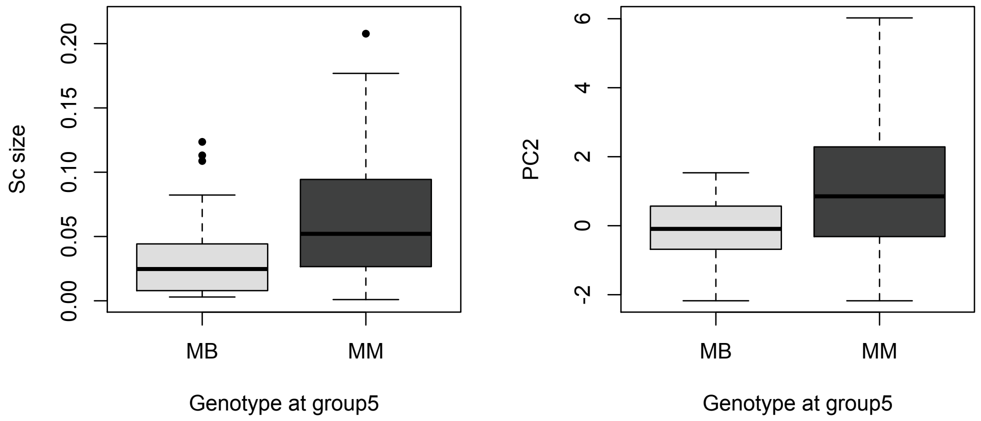

**Fig. S14.** Individuals that are heterozygous at the QTL peak on chromosome 5 have a lower probability of melanoma than individuals homozygous for *malinche* ancestry in this region (Fig. 3). We asked whether these individuals also had smaller spots (normalized for body size). Plotted here are only spotted individuals from the Chahuaco falls hybrid population, split by genotype at the chromosome 5 QTL peak. Individuals heterozygous for *malinche* ancestry have smaller spots (left, t-test of log-transformed data: p=0.007). Similarly, these individuals had significantly different distributions in their PC2 loadings (right; t-test p-value: $3x10^{-5}$) which correlates with both spot size and invasiveness (Fig. S2).

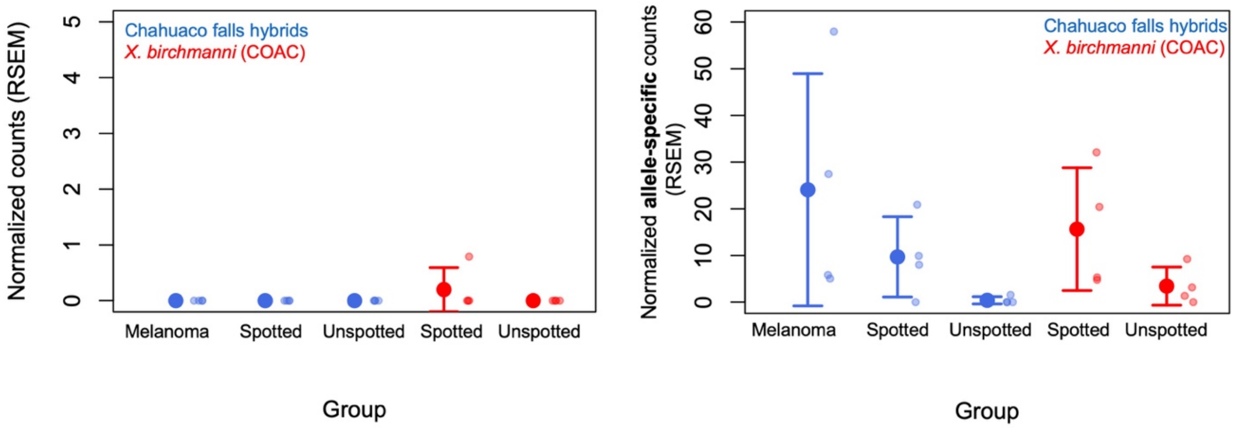

**Fig. S15.** Expression of GWAS hits *myrip* and *xmrk* in caudal tissue of *X. birchmanni* and natural hybrids. **A**) *myrip* has undetectable expression in caudal tissue of adult hybrids and low expression in *X. birchmanni* (regardless of spotting phenotype), suggesting that it is not expressed at the right time to be involved in the transition to melanoma. **B**) In contrast, although there is large variation in expression, *xmrk* is differentially expressed in caudal tissue between spotted and unspotted individuals (RSEM posterior probability of differential expression = 0.999). The most likely expression pattern inferred by RSEM is of three expression states, with the highest expression in spotted individuals regardless of group (hybrid or *X. birchmanni*) and lower expression in unspotted individuals, with significant differences in expression between unspotted hybrid and *X. birchmanni* individuals. Solid dots show the mean expression and whiskers show two standard errors of the mean. Transparent dots show raw data for each individual. Note that because of sequence similarities between *xmrk* and *egfrb* we used a modified approach to generate counts at informative sites for RSEM analysis, see details in Supporting Information *1.7.6 Determining the expression level of xmrk.*

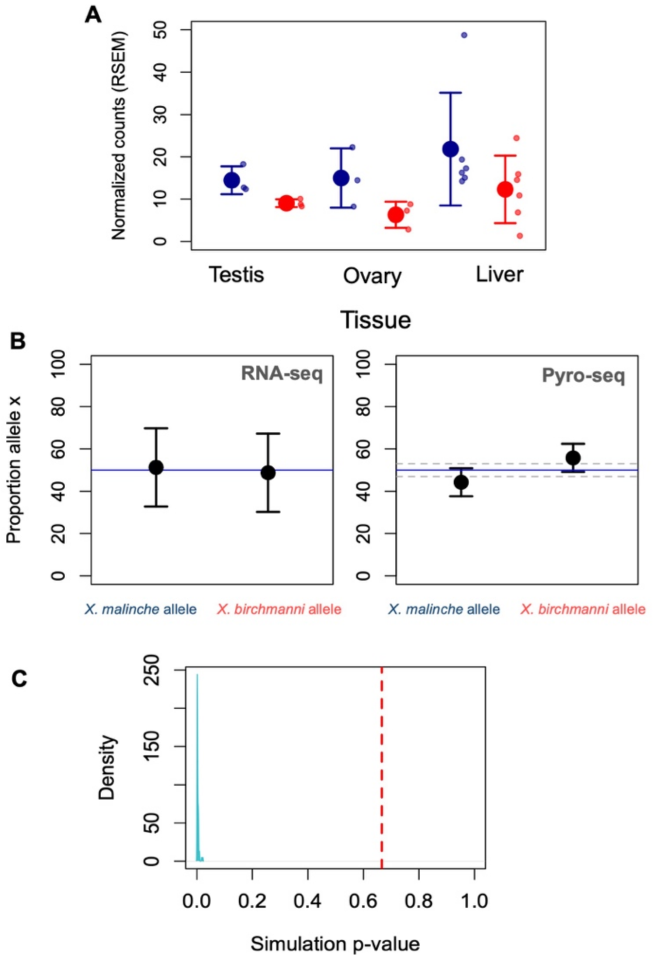

**Fig. S16. A)** Higher expression of *cd97* in *X. malinche* is not specific to the caudal tissue, and does not appear to be driven by *cis* regulatory changes (**B, C**). **B**) Estimates of allele specific expression of the *birchmanni* and *malinche* alleles in caudal fin tissue of four $F_1$ hybrids at *cd97* based on RNAseq (left) and pyrosequencing (right) data. Blue line indicates expectation of 50-50 expression of the two alleles, error bars show two standard deviations. Gray lines in pyrosequencing results indicate the variance expected in allelic estimates from pure parental individuals (see *1.7.3 Allele specific expression in F1 hybrids*). **C**) Despite the lack of signal in RNAseq data, simulations suggest that we expect to have excellent power to detect allele specific expression in our data. Shown here is a p-value distribution from 100 replicate simulations of allele specific expression at *cd97*, using the same analysis pipeline applied to the real data and mimicking other features of our dataset. The red line indicates the observed p-value for allele specific expression at *cd97* in the real data. See *1.7.3 Allele specific expression in F1 hybrids* and *1.7.4 Predicted power to detect allele specific expression* for more information. Solid dots and whiskers in **A** and **B** indicate means ± 2 s.e.m. Transparent dots in **A** indicate raw data for each individual.

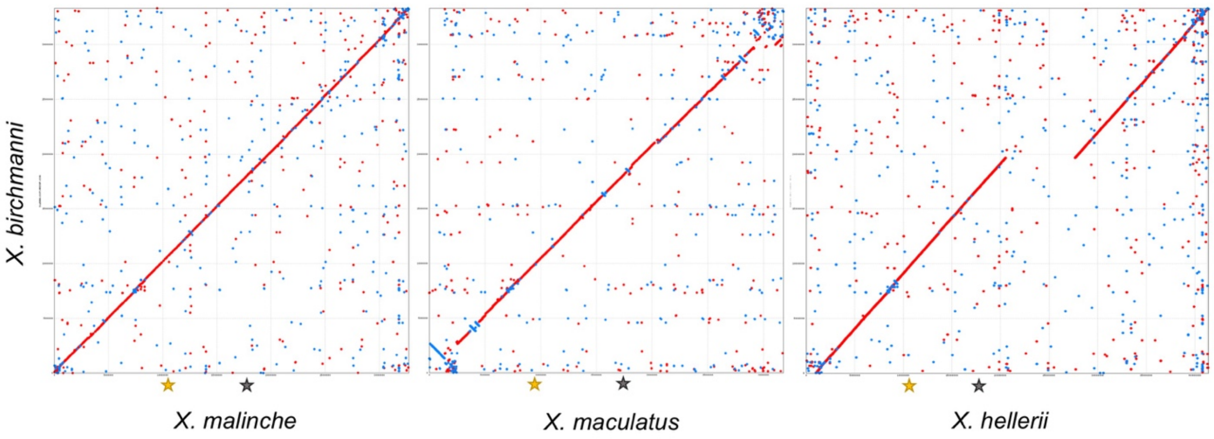

**Fig. S17.** Mapping results indicated that the interacting locus generating the melanoma incompatibility in crosses between *X. maculatus* and *X. hellerii* (region surrounding *cdkn2a/b*) is distinct from the locus in crosses between *X. birchmanni* and *X. malinche* (*cd97* region). However, the two regions both occur on chromosome 5. We examined whether these regions differed in their structure or relative position along the chromosome in any of these species by aligning chromosome 5 of each species to the *X. birchmanni* assembly with MUMmer. The yellow star indicates the location of *cd97* in each assembly and the gray star indicates the location of *cdkn2a/b*. The two genes are separated by ~7 Mb in each assembly.

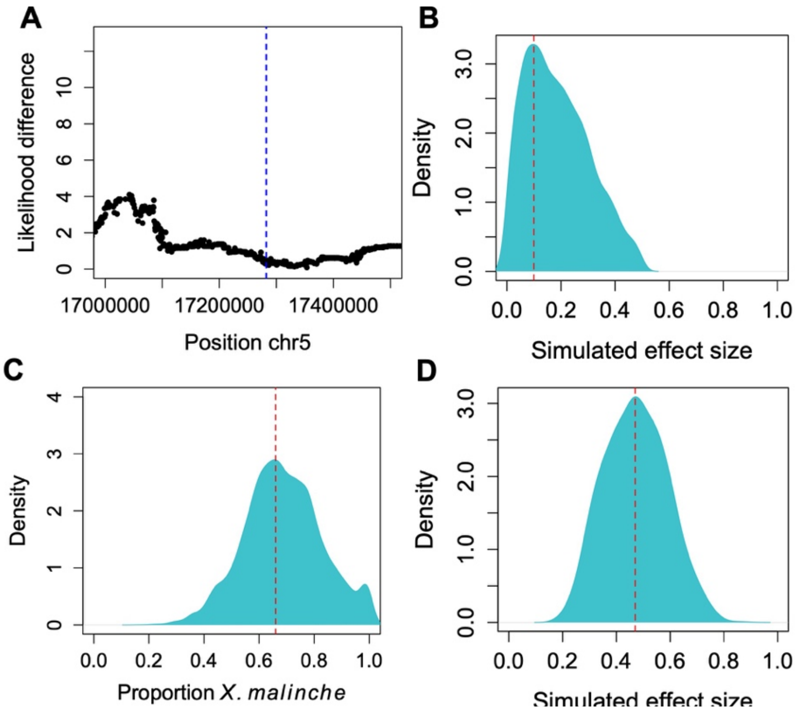

**Fig. S18.** Evaluation of the possible role of *cdkn2a/b* in the *X. birchmanni* - *X. malinche* melanoma. **A**) Zooming in on the region of chromosome 5 containing this gene (blue line) we do not see evidence for a sub-significant signal associated with melanoma. **B**) Posterior distribution of ABC simulations asking what effect sizes of *cdkn2a/b* are consistent with the observed likelihood difference in this region. The MAP estimate shown by the red line is 0.1 (95% confidence intervals: 0.013-0.43). **C**) Moreover, there is no evidence that this region has skewed ancestry (red line – ancestry at *cdkn2a/b*) compared to the genome-wide ancestry distribution (blue distribution). **D**) By contrast, the MAP estimate for the melanoma risk region we identified containing *cd97* is 0.5 (95% confidence intervals: 0.26-0.7). Note that these simulations accounted for the impact of the winner's curse on effect size estimates, see *1.6.5 No evidence for involvement of a previously mapped melanoma risk region* for details.

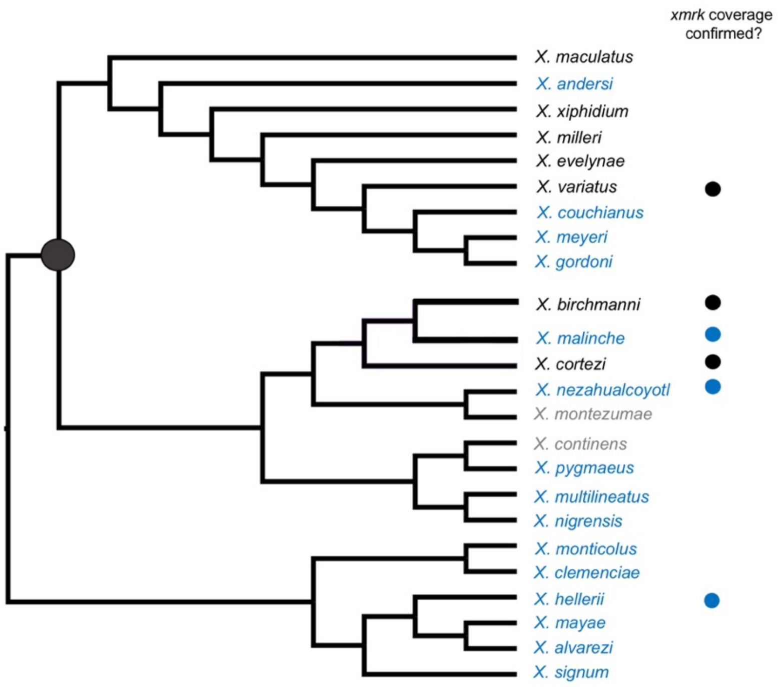

**Fig. S19.** Modified nuclear phylogeny of *Xiphophorus* including all described species except *X. kallman*i and *X. mixei* (*4*). *Xmrk* presence (black) or absence (blue) predicted from previous studies is indicated by colors in the species names. See *51* for a summary of these results. Cases where no data is available are indicated in gray. Cases where we were able to confirm previous reports using local coverage analysis (Fig. S20) are marked with colored circles (black for confirmed presence, blue for confirmed absence).

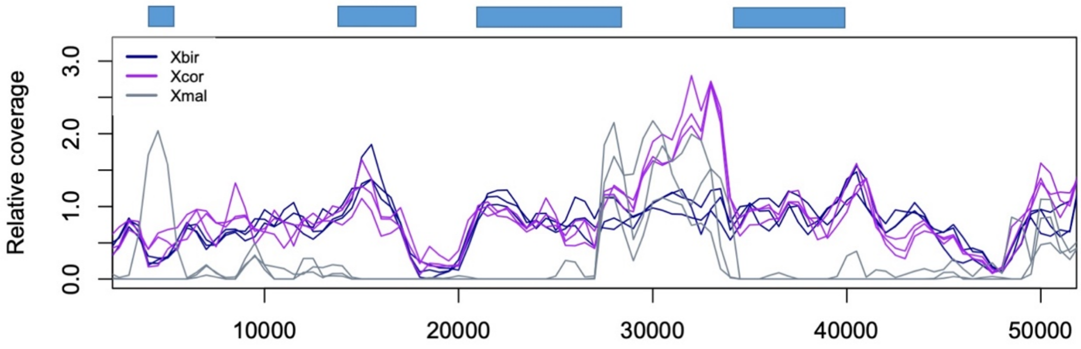

**Fig. S20.** Relative coverage in the *xmrk* region in different species based on mapping whole genome data from three individuals per species to the *X. birchmanni* (blue) reference genome suggests that *xmrk* has been deleted in *X. malinche* (gray) but is present in close relative *X. cortezi* (purple). Blue rectangles show the approximate locations of coding basepairs in the *xmrk* gene.

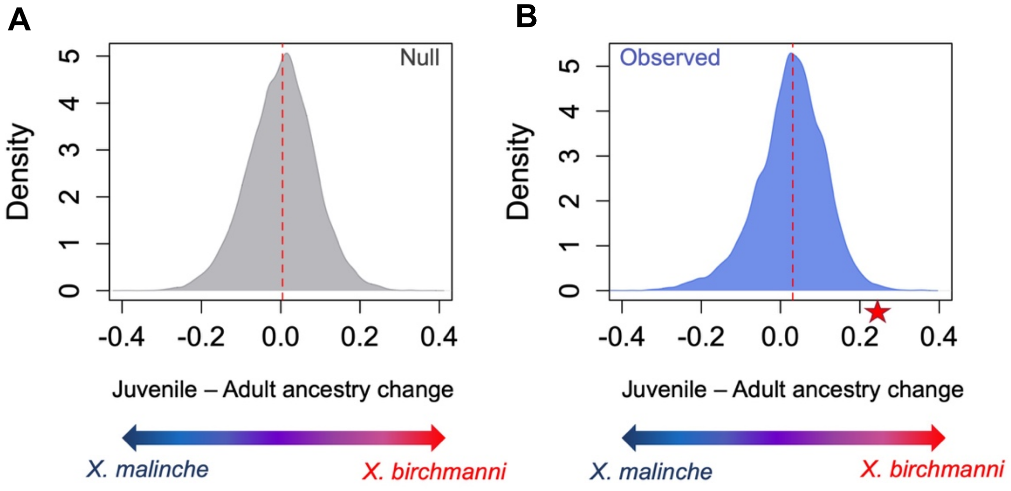

**Fig. S21.** We see strong shifts in spotting phenotype between the juvenile and adult stage in males from the Chahuaco falls population over multiple collection years (Fig. 4). These shifts in phenotype could be the product of selection on spotted individuals with melanoma or the result of genome-wide shifts in ancestry that coincidentally change the frequency of the spotted caudal. To investigate this, we analyzed ancestry data from adult and juvenile males collected from the same years. In **A**, we plot simulation results showing the expected distribution of differences in allele frequency given binomial sampling of individuals (see Supporting Information *1.9.1 Evaluating possible effects of population structure on juvenile-adult male frequency shifts*). The red line shows the median difference genome-wide. In **B**, we plot the observed distribution of estimates of juvenile-adult allele frequency differences. Compared to null expectations, we see a slight shift towards *X. birchmanni* ancestry (median = 0.02; indicated by red line), but little evidence for major shifts in genome-wide ancestry over time. Moreover, the melanoma-risk locus we mapped, indicated by the red star, is an outlier in terms of frequency changes between juveniles and adults (upper 1%). This locus shows strong shifts towards *X. birchmanni* ancestry in our dataset.

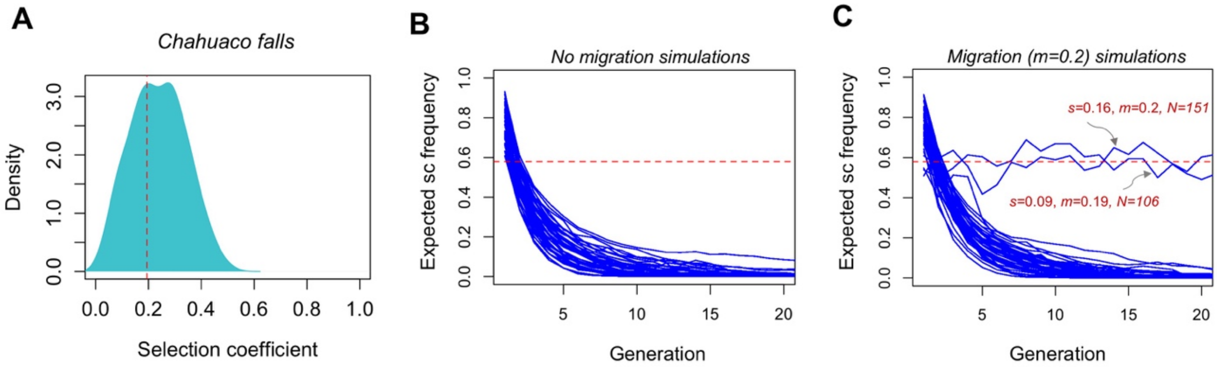

**Fig. S22.** Simulations suggest that under viability selection coefficients inferred by ABC simulations, the spotted caudal is not expected to persist in hybrid populations under most conditions. **A)** Posterior distribution of inferred viability selection coefficients inferred for the Chahuaco falls population. **B)** In the absence of migration, the trait is expected to be quickly purged from the population. Shown here are the results of admix'em simulations of hybrid populations sampling from the accepted selection coefficients plotted in **A**. **C)** Only in the presence of extremely high migration rates and in simulations with weaker selection coefficients do we see the trait occasionally persisting at high frequency.

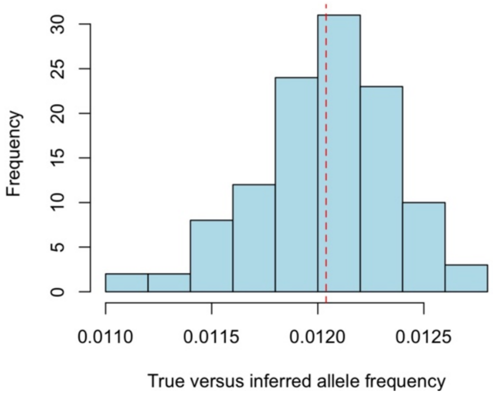

**Fig. S23.** Difference between true and inferred allele frequency in simulations using samtools-legacy to estimate allele frequency, applying the same approach used with our real data. Simulations used *X. birchmanni*-specific parameters and the average per-individual coverage observed in our real data. Plotted here is the average of the absolute value of allele frequency differences between the true frequency and inferred frequency in each of 100 simulations. The dashed red line indicates the mean difference across simulations.

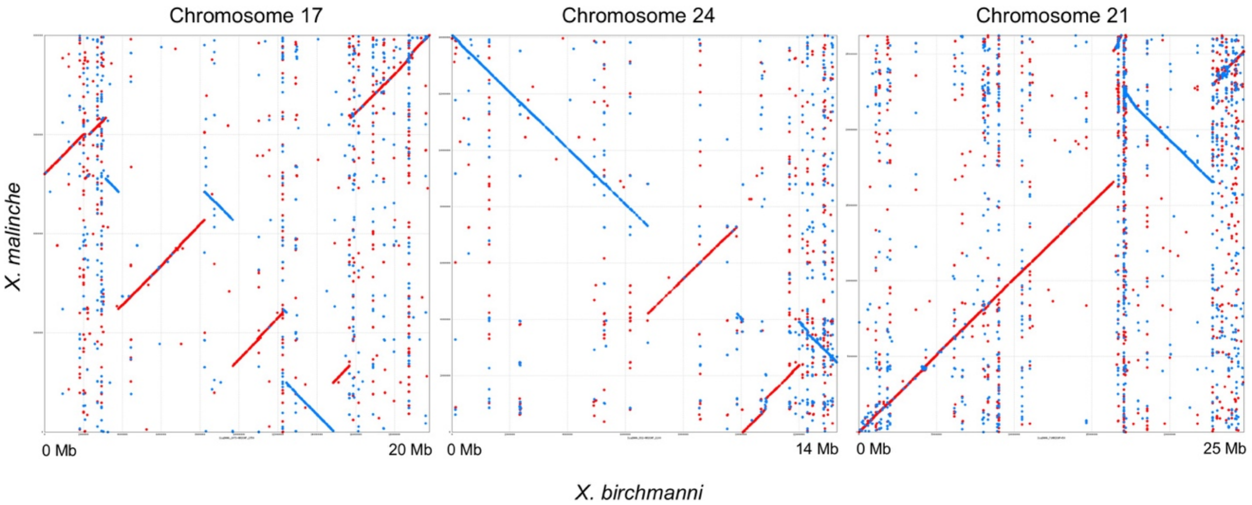

**Fig. S24.** MUMmer alignment of *X. birchmanni* and *X. malinche de novo* assemblies identifies structural differences between species, including previously known inversions on chromosome 17 and 24, and a newly identified structural rearrangement on chromosome 21. Further analysis suggests that this inversion is likely segregating at moderate frequencies within *X. birchmanni* (see <u>*1.5.1 Comparison of chromosome structure between X. birchmanni and X. malinche*</u>).

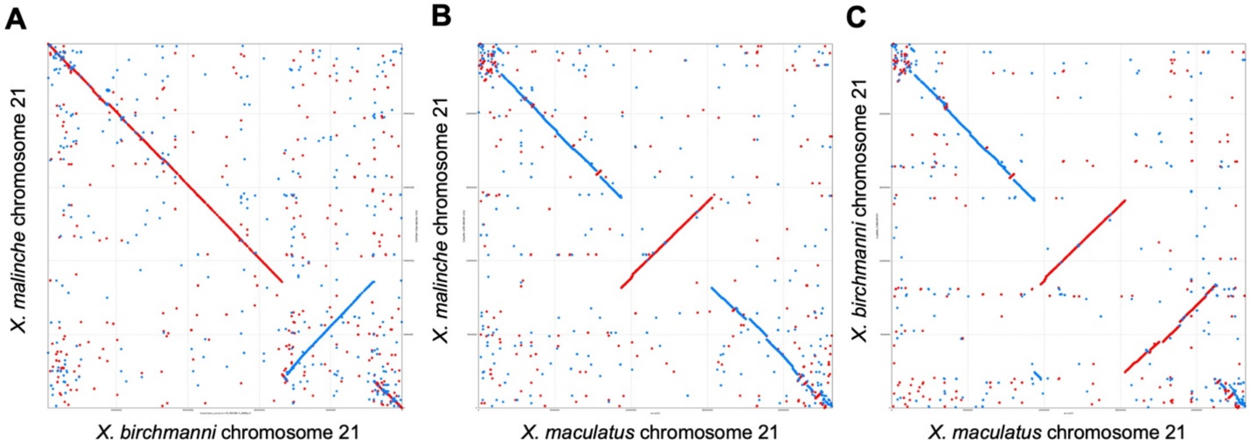

**Figure S25.** The assembled *X. birchmanni* and *X. malinche* genomes differ by a large inversion on chromosome 21, which can be visualized by the MUMmer plot in panel A. We aligned chromosome 21 from *X. birchmanni* and *X. malinche* to an outgroup assembly, *X. maculatus*, to determine which arrangement was shared with the outgroup. Interestingly, this reveals a rearrangement between both species and *X. maculatus* elsewhere in the chromosome (**B**, **C**). However, these alignments suggest that the inversion identified when comparing the *birchmanni*-*malinche* assemblies is derived in *X. birchmanni*.

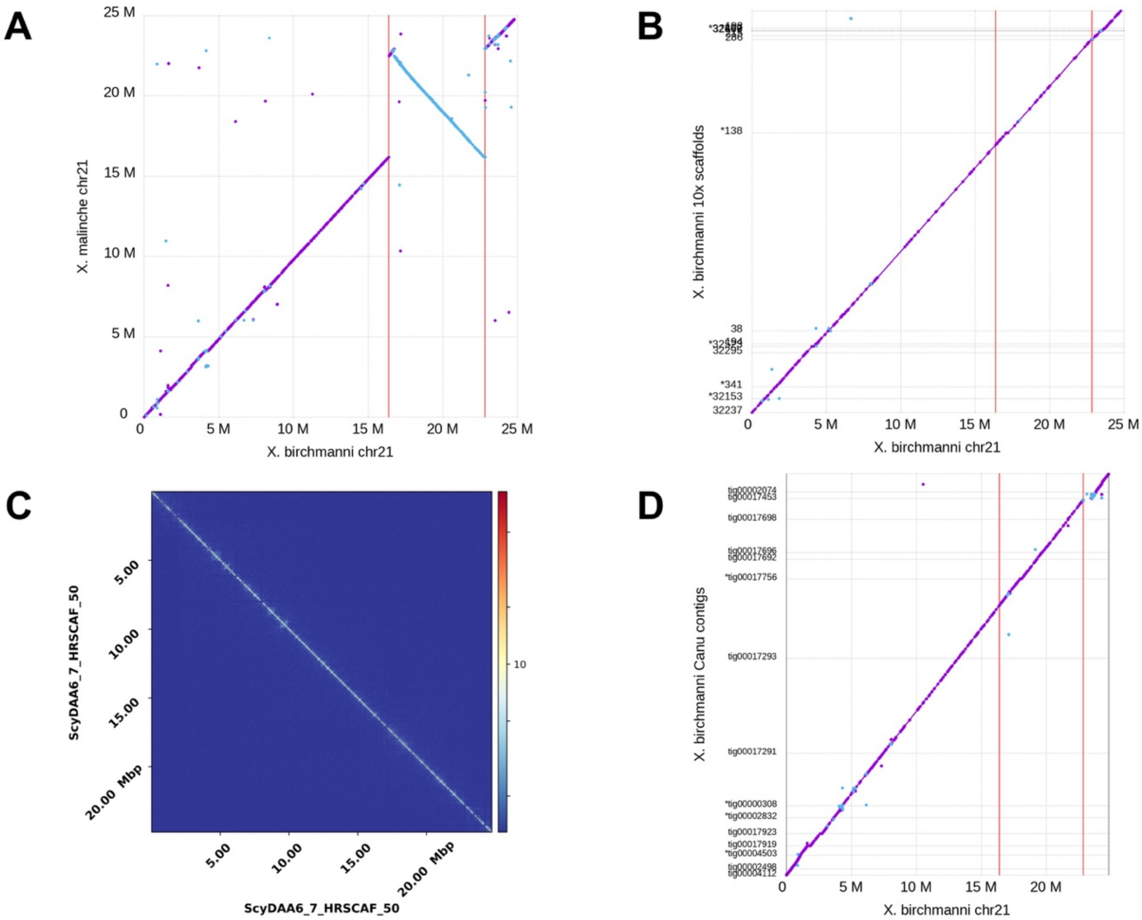

**Fig. S26.** Support for a chromosome 21 inversion in *X. birchmanni*: **A**) MUMmer alignment of chromosome 21 of *X. malinche* (y-axis) and *X. birchmanni* (x-axis), red vertical bars denote the inversion breakpoints. **B**) MUMmer alignment of 10x Supernova scaffolds (y-axis) against the final *X. birchmanni* assembly (x-axis). Note that although the left inversion breakpoint is spanned an assembled scaffold, the right breakpoint is not spanned. **C**) Hi-C contact map of *X. birchmanni* chromosome 21, showing no off-diagonal signal of the inversion. This indicates that the Hi-C individual was homozygous for the inverted (i.e. non-*X. malinche*) arrangement. **D**) MUMmer alignment of Canu contigs (y-axis) against the final *X. birchmanni* assembly (x-axis).

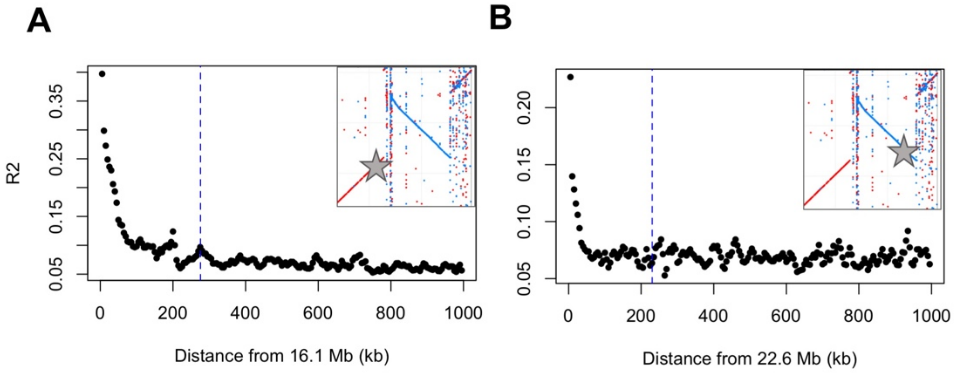

**Fig. S27.** Linkage disequilibrium surrounding the putative inversion breakpoints in population samples of *X. birchmanni*. **A**) Parental population LD decay is shown near the five-prime edge of the putative segregating inversion on chromosome 21. There is weak evidence of unusual LD patterns in this region. In contrast, there are no clear deviations observed at the three-prime edge of the putative inversion (**B**). The blue dashed lines indicate the inversion breakpoints estimated by LUMPY but may be imprecise. Stars in insets indicate the approximate position of the zero coordinate on the x-axis.

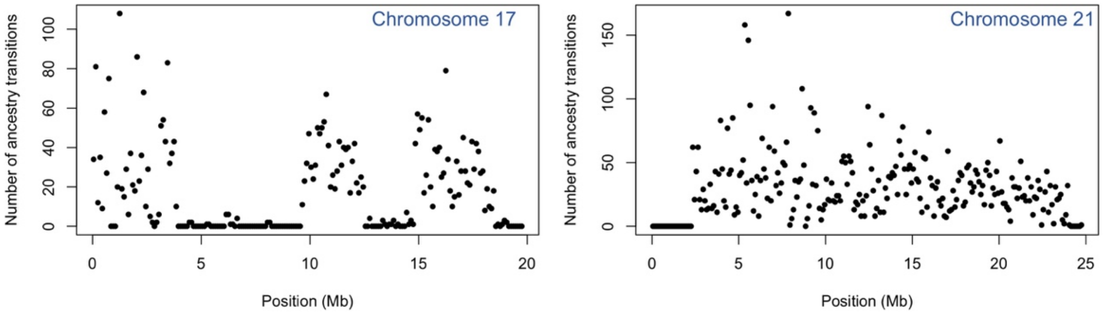

**Fig. S28.** Ancestry transitions observed on chromosome 17 and chromosome 21 in Chahuaco Falls hybrids. As expected if inversions cause recombination suppression in heterozygotes and the inversion is fixed, we observe few ancestry transitions in hybrids in the inverted region on chromosome 17. However, the same is not the case for the inversion identified on chromosome 21, consistent with other data suggesting that it is unlikely to be a fixed difference between species. See *1.5.1 Comparison of chromosome structure between X. birchmanni and X. malinche* for more information.

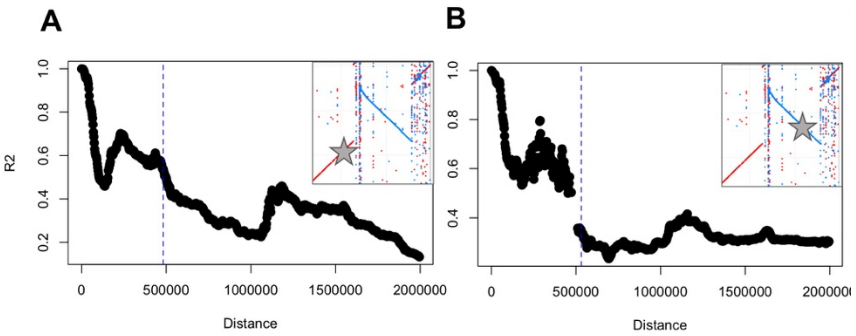

**Fig. S29.** Patterns of admixture LD decay over physical distance in the Chahuaco falls population in regions near the five-prime (**A**) and three-prime (**B**) edges of the putative segregating inversion on chromosome 21. The blue dashed lines indicate the inversion breakpoints estimated by LUMPY but may be imprecise. Stars in insets indicate the approximate position of the zero coordinate on the x-axis. Average patterns genome wide can be seen in Fig. S34.

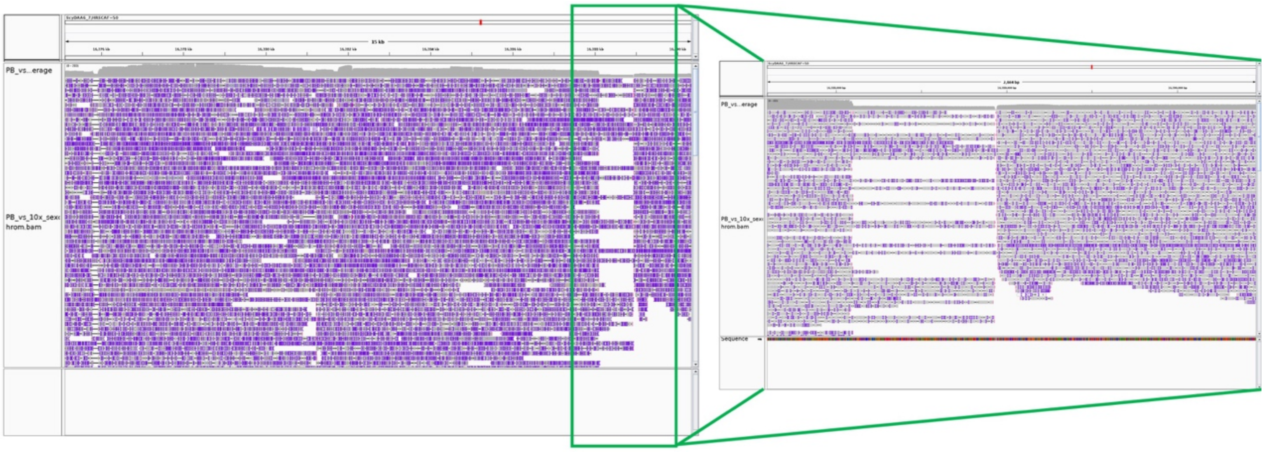

**Fig. S30.** Integrative Genomics Viewer snapshot of PacBio read aligned to the *X. birchmanni* reference assembly at the left breakpoint of the chromosome 21 inversion. This shows that the individual for which PacBio data was collected is heterozygous for the *X. birchmanni* arrangement, since a number of PacBio reads span the breakpoint. The image on the left spans the left inversion breakpoint, and inset on the right provide a closer view of the putative breakpoint. The right breakpoint is not shown since we were unable to identify PacBio reads that spanned this gap.

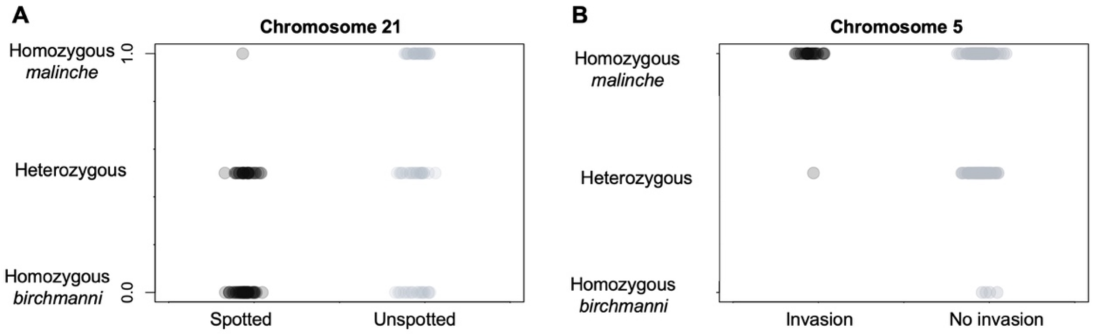

**Fig. S31.** Individual genotypes as a function of phenotype at QTL peaks on chromosome 21 and chromosome 5. **A**) Nearly all spotted individuals have *X. birchmanni* ancestry at chromosome 21, whereas unspotted individuals are more likely to be homozygous for *X. malinche* ancestry. Note that because the spotted caudal is polymorphic in *X. birchmanni* it is not surprising that some unspotted individuals have *X. birchmanni* ancestry at this region. **B**) Individuals with melanocyte invasion are more likely to be homozygous for *X. malinche* ancestry at the chromosome 5 QTL peak.

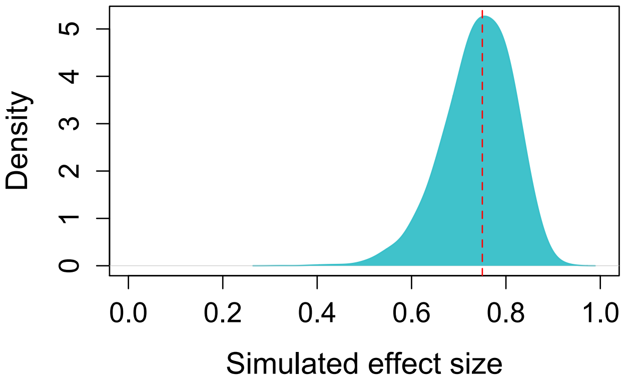

**Fig. S32.** Posterior distribution from ABC simulations estimating the effect size of *X. birchmanni* ancestry at the chromosome 21 QTL on spotting phenotype. The MAP estimate shown by the red line is 0.75 (95% confidence intervals: 0.57 - 0.86). These simulations are described in Supporting Information section *1.6.4 Estimating the effect size of the QTL associated with the spotted caudal.*

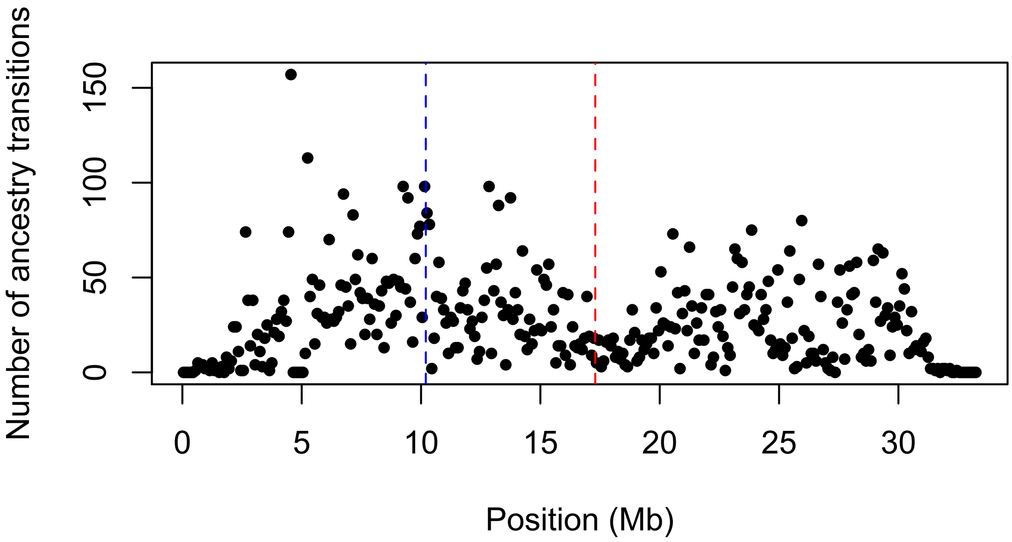

**Fig. S33.** Frequency of ancestry transitions per 100 kb window along chromosome 5 in 209 Chahuaco falls hybrid individuals. The blue and red lines show the locations of the *cd97* and *cdnk2a/b* genes respectively, indicating substantial overall recombination between these regions in hybrids.

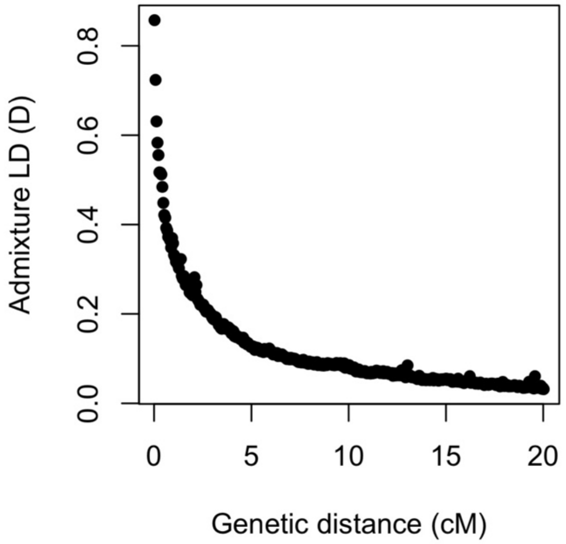

**Fig. S34.** Observed average decay in admixture linkage disequilibrium over genetic distance in the Chahuaco falls hybrid population. Fitting an exponential model to this curve, we recovered an estimate of time since initial admixture of 46 ± 1 generations. However, we caution that this is likely an underestimate because of ongoing migration.

```
xbirchmanni     MGFGKGLLILGFVCALQRPVFGCRNGFQNNDGKCEDINECNDPDFCGNHAKCINNNGSYY    60
xmalinche       MGFGKGLLILGFVCALQRPVFGCRNGFQNNDGKCEDINECNDPDFCGNHAKCINNNGSYY    60
                ************************************************************

xbirchmanni     CECSAGFHNVKGNPNFTVEAGQCTDINECLDNNTTCGPHGTCNNVIGHYKCVCNAGYVSA    120
xmalinche       CECSAGFHNVKGNPNFTVEAGQCTDINECLDNNTTCGPHGTCNNVIGHYKCVCNAGYVSA    120
                ************************************************************

xbirchmanni     DVSNKTAHCRDIDECKEHKDVCGEFGVCENLNGSYRCICEAGHTNYGQEKTRCSKLECDS    180
xmalinche       DASNKTAHCRDIDECKEHKDVCGEKGVCENLNGSYRCICEAGHTNYGQEKTRCSKLECDS    180
                *.******************* **************************************

xbirchmanni     FHTDSDPGKSFEGLADMLSMMRNSCLALSDPSYSAGGKADGDALLEKLLTATDTILSPAH    240
xmalinche       FHADSDPGKSFEGLADMLSMMRNSCLALSDPSYSAGGKADGDALLEKLLTATDTILSPAH    240
                **:*********************************************************

xbirchmanni     VENSKGVSGLLTAVEKSILLIGPQLKANDTKMETKETEAKITVRRGASRPTGQIHLTSEH    300
xmalinche       VENSKGVSGLLTAVEKSILLIGPQLKANDTKMETKETEAKITVRRGASRPTGQIHLTSEH    300
                ************************************************************

xbirchmanni     ADLNTDWTTAAGEGPYPGFALAALLSYKNLEESVNRSFEDLAEKEKDAVSFQVFSKVVSV    360
xmalinche       ADLNTDWTTAAGEGPYPGFALAALLSYKNLEETVNRSFEDLAEKEKDAVSFQVFSKVVSV    360
                ********************************:***************************

xbirchmanni     VVSNPSTQNLHRPVNITFRHLKDIKQTPEVSYICAYWDERGVWSPDGCSQELSNDTHTVC    420
xmalinche       VVSNPSTQNLHRPVNITFRHLKDIKQTPEVSYICAYWDERGVWSPDGCSQELSNDTHTVC    420
                ************************************************************

xbirchmanni     SCEHLSSFAVLMALYPMKTSFHLQLLTQIGLTISLVCLVLSILTFKFCRSIQGTRTTIHL    480
xmalinche       SCEHLSSFAVLMALYPMKTSFHLQLLTQIGLTISLVCLVLSILTFKFCRSIQGTRTTIHL    480
                ************************************************************

xbirchmanni     HLCICLFMADLVFLVGISRTRPEGGCKFVAAMLHFFFLGVMSWMLLEGVQLYRMVVLVFN    540
xmalinche       HLCICLFMADLVFLVGISRTRPEGGCKFVAGMLHFFFLGVMSWMLLEGVQLYRMVVLVFN    540
                ******************************.*****************************

xbirchmanni     ATIRPLYLYLTGYGIPLGIVIISVIIRPGGYGTKDHCWLSLEHGLIWSFFGPVCFIIALN    600
xmalinche       ATIRPLYLYLTGYGIPLGIVIISVIIRPGGYGTKDHCWLSLEHGLIWSFFGPVCFIIALN    600
                ************************************************************

xbirchmanni     VFFFIVTVWKLAQKFATLNPDLSKLHKIKAFTVTAIAQMCILGLMWVFGAFLFSEGMTAV    660
xmalinche       VFFFIVTVWKLAQKFATLNPDLSKLHKIKAFTVTAIAQMCILGLMWVFGAFLFSEGMTAV    660
                ************************************************************

xbirchmanni     AYIFTVLNSLQGALVFIMHCVLSKQVRDEYVQFLSCVCTPQKKRYSDFSSTNPSSSQSQG    720
xmalinche       AYIFTVLNSLQGALVFIMHCVLSKQVRDEYVQFLSCVCTPQKKRYSDFSSTNPSSSQSQG    720
                ************************************************************

xbirchmanni     SRSGQITGESQI   732
xmalinche       SRSGQITGESQI   732
                ************
```

**Fig. S35.** Clustal alignment of translated nucleotide sequences of *X. birchmanni* and *X. malinche* at the *cd97* gene. Stars indicate places where the two sequences are identical; colons, periods and blanks indicate spaces where an amino acid change has occurred and dashes indicate possible insertions or deletions.

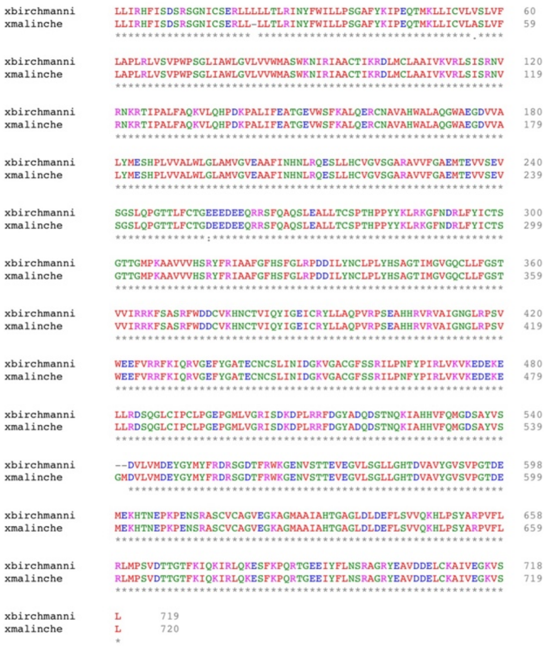

**Fig. S36.** Clustal alignment of translated nucleotide sequences of *X. birchmanni* and *X. malinche* at the *slc27a1b* gene. Stars indicate places where the two sequences are identical, colons, periods and blanks indicate spaces where an amino acid change has occurred and dashes indicate possible insertions or deletions. Note the non-canonical lysine start codon; this start codon was confirmed with sanger sequencing.

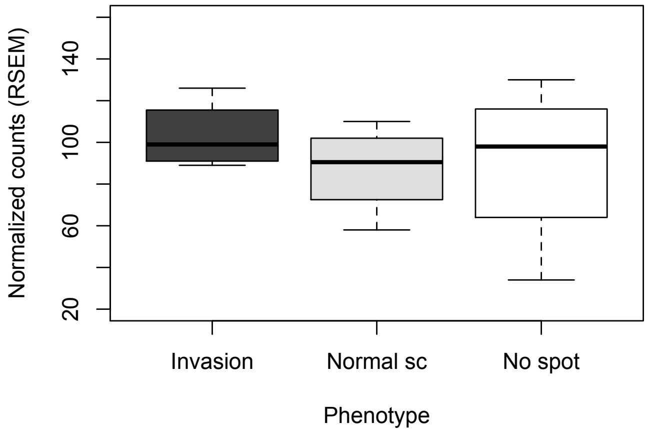

**Fig. S37.** Expression of *slc27a1b* across different spotted phenotypes in Chahuaco falls hybrid individuals. Plotted here are normalized counts produced by RSEM for *slc27a1b* expression from four Chahuaco falls hybrids of each phenotype. In contrast to the other gene in the QTL interval, *cd97*, we see no evidence of differential expression between individuals with and without melanoma (p=0.3).

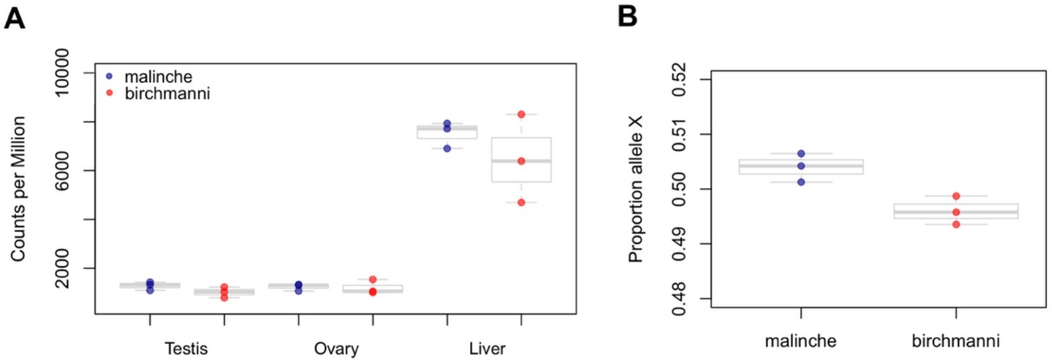

**Fig. S38.** Expression level of the housekeeping gene used in qPCR, *efa1*, is similar in *X. birchmanni* and *X. malinche* in a number of tissues (**A**), suggesting that it is an appropriate housekeeping gene for our purposes. The data shown here reflect expression level determined from RNAseq data derived from several tissues. Similarly, there is not evidence of allele specific expression of *efa1* based on our analysis (**B**), (see *1.7.4 Predicted power to detect allele specific expression*).

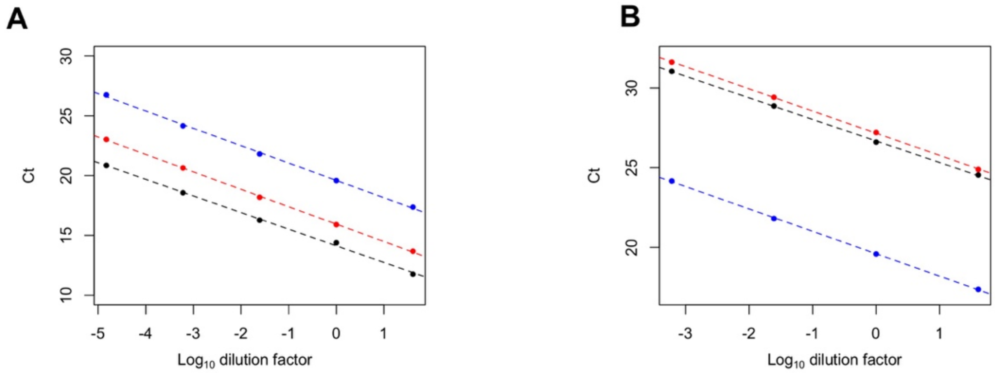

**Fig. S39.** Comparison of slopes across biological conditions for serial dilution tests to estimate qPCR efficiency for *efa1* (**A**) and *cd97* (**B**) qPCR primers. Although there are slight differences in the point estimates of efficiency across conditions (Table S2), there is not evidence for differences in slope (see *1.7.1 Real-time quantitative PCR to evaluate cd97 expression*). Note that the Ct of *X. birchmanni* for *cd97* reflects a high overall expression level because a PCR-product was used in efficiency dilutions due to low concentrations of *cd97* in *X. birchmanni*.

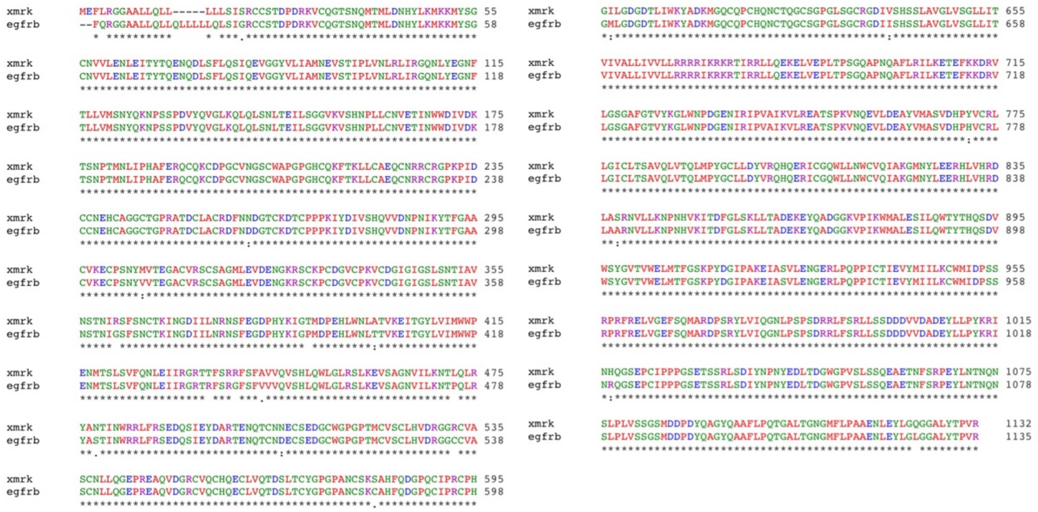

**Fig. S40.** Clustal alignment of *xmrk* and *egfrb* protein sequences from *X. birchmanni*. Stars indicate places where the two sequences are identical; colons, periods and blanks indicate places where an amino acid change has occurred and dashes indicate possible insertions or deletions.

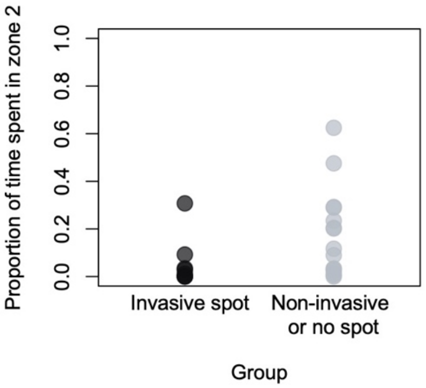

**Fig. S41.** Performance in endurance swim trials as a function of melanoma state in hybrids. Shown here for visualization is the time spent in zone 2 by individuals with and without melanoma. Zone 2 is the region of the tank furthest from the pump and fish that spent time in this zone were deemed to have experienced swimming exhaustion (see *1.10 Evaluating the impact of spotted caudal melanoma on swimming performance*). We detected no significant impact of spotting phenotype on exhaustion in a general linear model (p=0.1), but note that most individuals experienced little or no exhaustion during the course of the trials.

#### 4. Supplementary Tables

**Table S1.** Complete gene ontogeny results based on GOstats analysis of genes differentially expressed between melanotic and normal caudal tissue. A version of this analysis thinned with REVIGO is shown in Figure S1.

| GOBPID | P-value | Odds Ratio | Expected Count | Count | Size | Term |
|---|---|---|---|---|---|---|
| GO:0006397 | 0.000 | 5.3 | 2.6 | 12 | 98 | mRNA processing |
| GO:0016071 | 0.000 | 4.5 | 3.6 | 14 | 132 | mRNA metabolic process |
| GO:0008380 | 0.000 | 5.5 | 2.1 | 10 | 79 | RNA splicing |
| GO:0000398 | 0.000 | 6.5 | 1.5 | 8 | 54 | mRNA splicing, via spliceosome |
| GO:0000375 | 0.000 | 6.5 | 1.5 | 8 | 54 | RNA splicing, via transesterification reactions |
| GO:0000377 | 0.000 | 6.5 | 1.5 | 8 | 54 | RNA splicing, via transesterification reactions with bulged adenosine as nucleophile |
| GO:0048730 | 0.001 | Inf | 0.1 | 2 | 2 | epidermis morphogenesis |
| GO:0000387 | 0.004 | 12.2 | 0.3 | 3 | 12 | spliceosomal snRNP assembly |
| GO:0006396 | 0.005 | 2.4 | 6.4 | 14 | 237 | RNA processing |
| GO:0001837 | 0.008 | 8.4 | 0.4 | 3 | 16 | epithelial to mesenchymal transition |
| GO:0010467 | 0.009 | 1.6 | 31.1 | 44 | 1152 | gene expression |
| GO:0050808 | 0.009 | 5.4 | 0.8 | 4 | 31 | synapse organization |
| GO:0022607 | 0.012 | 1.9 | 10.8 | 19 | 401 | cellular component assembly |
| GO:0006334 | 0.014 | 14.6 | 0.2 | 2 | 7 | nucleosome assembly |
| GO:0015914 | 0.018 | 6.1 | 0.6 | 3 | 21 | phospholipid transport |
| GO:0030318 | 0.018 | 6.1 | 0.6 | 3 | 21 | melanocyte differentiation |
| GO:0006333 | 0.018 | 12.1 | 0.2 | 2 | 8 | chromatin assembly or disassembly |
| GO:0016226 | 0.018 | 12.1 | 0.2 | 2 | 8 | iron-sulfur cluster assembly |
| GO:0051262 | 0.018 | 12.1 | 0.2 | 2 | 8 | protein tetramerization |
| GO:0031497 | 0.018 | 12.1 | 0.2 | 2 | 8 | chromatin assembly |
| GO:0031163 | 0.018 | 12.1 | 0.2 | 2 | 8 | metallo-sulfur cluster assembly |
| GO:0072583 | 0.018 | 12.1 | 0.2 | 2 | 8 | clathrin-dependent endocytosis |
| GO:0007064 | 0.018 | 12.1 | 0.2 | 2 | 8 | mitotic sister chromatid cohesion |
| GO:0008152 | 0.018 | 1.4 | 103.5 | 118 | 3835 | metabolic process |
| GO:0035556 | 0.019 | 1.7 | 13.0 | 21 | 480 | intracellular signal transduction |
| GO:0044085 | 0.019 | 1.7 | 12.2 | 20 | 452 | cellular component biogenesis |
| GO:0015748 | 0.020 | 5.8 | 0.6 | 3 | 22 | organophosphate ester transport |
| GO:0006869 | 0.021 | 3.0 | 2.2 | 6 | 80 | lipid transport |
| GO:0065003 | 0.021 | 2.1 | 5.5 | 11 | 203 | protein-containing complex assembly |
| GO:0016043 | 0.021 | 1.5 | 27.5 | 38 | 1019 | cellular component organization |
| GO:0034622 | 0.022 | 2.3 | 4.1 | 9 | 152 | cellular protein-containing complex assembly |
| GO:0000956 | 0.023 | 5.5 | 0.6 | 3 | 23 | nuclear-transcribed mRNA catabolic process |

| | | | | | | |
|---|---|---|---|---|---|---|
| GO:0071704 | 0.024 | 1.4 | 93.1 | 107 | 3450 | organic substance metabolic process |
| GO:0006468 | 0.024 | 1.7 | 13.3 | 21 | 493 | protein phosphorylation |
| GO:0010876 | 0.026 | 2.8 | 2.3 | 6 | 84 | lipid localization |
| GO:0007276 | 0.026 | 5.2 | 0.6 | 3 | 24 | gamete generation |
| GO:0048609 | 0.026 | 5.2 | 0.6 | 3 | 24 | multicellular organismal reproductive process |
| GO:0032504 | 0.026 | 5.2 | 0.6 | 3 | 24 | multicellular organism reproduction |
| GO:0043966 | 0.027 | Inf | 0.0 | 1 | 1 | histone H3 acetylation |
| GO:0070054 | 0.027 | Inf | 0.0 | 1 | 1 | mRNA splicing, via endonucleolytic cleavage and ligation |
| GO:0021772 | 0.027 | Inf | 0.0 | 1 | 1 | olfactory bulb development |
| GO:0090522 | 0.027 | Inf | 0.0 | 1 | 1 | vesicle tethering involved in exocytosis |
| GO:0003347 | 0.027 | Inf | 0.0 | 1 | 1 | epicardial cell to mesenchymal cell transition |
| GO:0021988 | 0.027 | Inf | 0.0 | 1 | 1 | olfactory lobe development |
| GO:0098743 | 0.027 | Inf | 0.0 | 1 | 1 | cell aggregation |
| GO:0001502 | 0.027 | Inf | 0.0 | 1 | 1 | cartilage condensation |
| GO:0032285 | 0.027 | Inf | 0.0 | 1 | 1 | non-myelinated axon ensheathment |
| GO:0032233 | 0.027 | Inf | 0.0 | 1 | 1 | positive regulation of actin filament bundle assembly |
| GO:0070534 | 0.027 | Inf | 0.0 | 1 | 1 | protein K63-linked ubiquitination |
| GO:1903391 | 0.027 | Inf | 0.0 | 1 | 1 | regulation of adherens junction organization |
| GO:1903392 | 0.027 | Inf | 0.0 | 1 | 1 | negative regulation of adherens junction organization |
| GO:0060389 | 0.027 | Inf | 0.0 | 1 | 1 | pathway-restricted SMAD protein phosphorylation |
| GO:0051496 | 0.027 | Inf | 0.0 | 1 | 1 | positive regulation of stress fiber assembly |
| GO:0140238 | 0.027 | Inf | 0.0 | 1 | 1 | presynaptic endocytosis |
| GO:0048488 | 0.027 | Inf | 0.0 | 1 | 1 | synaptic vesicle endocytosis |
| GO:0099022 | 0.027 | Inf | 0.0 | 1 | 1 | vesicle tethering |
| GO:0010961 | 0.027 | Inf | 0.0 | 1 | 1 | cellular magnesium ion homeostasis |
| GO:0071840 | 0.027 | 1.5 | 28.9 | 39 | 1069 | cellular component organization or biogenesis |
| GO:0007528 | 0.028 | 9.1 | 0.3 | 2 | 10 | neuromuscular junction development |
| GO:0051091 | 0.028 | 9.1 | 0.3 | 2 | 10 | positive regulation of DNA-binding transcription factor activity |
| GO:0034728 | 0.028 | 9.1 | 0.3 | 2 | 10 | nucleosome organization |
| GO:0007062 | 0.028 | 9.1 | 0.3 | 2 | 10 | sister chromatid cohesion |
| GO:0016070 | 0.029 | 1.5 | 27.2 | 37 | 1007 | RNA metabolic process |
| GO:0048762 | 0.029 | 3.1 | 1.7 | 5 | 64 | mesenchymal cell differentiation |
| GO:1902905 | 0.029 | 5.0 | 0.7 | 3 | 25 | positive regulation of supramolecular fiber organization |
| GO:0061564 | 0.030 | 2.3 | 3.7 | 8 | 136 | axon development |
| GO:0051495 | 0.032 | 4.8 | 0.7 | 3 | 26 | positive regulation of cytoskeleton organization |
| GO:0048706 | 0.033 | 2.7 | 2.4 | 6 | 89 | embryonic skeletal system development |
| GO:0014044 | 0.034 | 8.1 | 0.3 | 2 | 11 | Schwann cell development |
| GO:0014037 | 0.034 | 8.1 | 0.3 | 2 | 11 | Schwann cell differentiation |

| | | | | | | |
|---|---|---|---|---|---|---|
| GO:0050931 | 0.035 | 4.6 | 0.7 | 3 | 27 | pigment cell differentiation |
| GO:0006323 | 0.040 | 7.3 | 0.3 | 2 | 12 | DNA packaging |
| GO:0044089 | 0.042 | 4.2 | 0.8 | 3 | 29 | positive regulation of cellular component biogenesis |
| GO:0043933 | 0.045 | 1.9 | 6.2 | 11 | 229 | protein-containing complex subunit organization |
| GO:0051090 | 0.047 | 6.6 | 0.4 | 2 | 13 | regulation of DNA-binding transcription factor activity |

**Table S2.** Primer sequences used for qPCR analysis of *cd97* expression in caudal fin tissues from *X. birchmanni*, *X. malinche*, and F1 hybrids.

| Target gene | Forward | Reverse | Efficiency |
|---|---|---|---|
| *cd97* | TGTGGGAACCACGCAAAGT | TCCAGTTTCAGCACAGTCGG | *X. malinche* - 105%<br>*X. birchmanni* - 98%<br>$F_1$ - 108% |
| *efa1* (house-keeping) | CCCCTAACCTGACCACTGAA | GTGGGTCGTTCTTGCTGTCT | *X. malinche* - 100%<br>*X. birchmanni* - 98%<br>$F_1$ - 103% |

**Table S3.** High coverage data from previous studies used in population genetic and phylogenetic analyses. Average per basepair coverage when mapped to the *X. birchmanni* reference genome is listed.

| **Species** | **SRA accession** | **Coverage** |
|---|---|---|
| *X. hellerii* | SRR7532852 | 85 |
| *X. maculatus* | SRR7532852 | 91 |
| *X. montezumae* | SRR3086791 | 21 |
| *X. nezahualcoyotl* | SRR3086878 | 24 |
| *X. malinche* | SRR6649369 | 36 |
